# HLA binding of self-peptides is biased towards proteins with specific molecular functions

**DOI:** 10.1101/2021.02.16.431395

**Authors:** Vadim Karnaukhov, Wayne Paes, Isaac B. Woodhouse, Thomas Partridge, Annalisa Nicastri, Simon Brackenridge, Dmitrii Scherbinin, Dmitry M. Chudakov, Ivan V. Zvyagin, Nicola Ternette, Hashem Koohy, Persephone Borrow, Mikhail Shugay

## Abstract

Human leukocyte antigen (HLA) is highly polymorphic and plays a key role in guiding adaptive immune responses by presenting foreign and self peptides to T cells. Each HLA variant selects a minor fraction of peptides that match a certain motif required for optimal interaction with the peptide-binding groove. These restriction rules define the landscape of peptides presented to T cells. Given these limitations, one might suggest that the choice of peptides presented by HLA is non-random and there is preferential presentation of an array of peptides that is optimal for distinguishing self and foreign proteins. In this study we explore these preferences with a comparative analysis of self peptides enriched and depleted in HLA ligands. We show that HLAs exhibit preferences towards presenting peptides from certain proteins while disfavoring others with specific functions, and highlight differences between various HLA genes and alleles in those preferences. We link those differences to HLA anchor residue propensities and amino acid composition of preferentially presented proteins. The set of proteins that peptides presented by a given HLA are most likely to be derived from can be used to distinguish between class I and class II HLAs and HLA alleles. Our observations can be extrapolated to explain the protective effect of certain HLA alleles in infectious diseases, and we hypothesize that they can also explain susceptibility to certain autoimmune diseases and cancers. We demonstrate that these differences lead to differential presentation of HIV, influenza virus, SARS-CoV-1 and SARS-CoV-2 proteins by various HLA alleles. Finally, we show that the reported self peptidome preferences of distinct HLA variants can be compensated by combinations of HLA-A/HLA-B and HLA-A/HLA-C alleles in frequent haplotypes.

## Introduction

T cells detect pathogen-infected and abnormal (e.g. tumour) cells by monitoring cell-surface-displayed short peptides presented by the human leukocyte antigen (HLA) complex. HLA molecules are highly specific in terms of the peptide sequences they are able to present, and peptides not presented by HLAs remain invisible to the immune system (1). HLA class I (HLA-I) and HLA class II (HLA-II) molecules present peptides that are typically recognised as a complex by CD8 and CD4 T cells, respectively; HLA-I-peptide complexes are also engaged by activating and inhibitory receptors on innate lymphocyte subsets such as natural killer (NK) cells. The three classical HLA-I genes expressed in all nucleated cells in humans are HLA-A, HLA-B, and HLA-C. HLA-I molecules present peptides derived from intracellular proteins. The intracellular antigen presentation pathway involves cleavage of proteins in the cytosol by proteasomes, translocation to the endoplasmic reticulum (ER) lumen, trimming by ER-resident aminopeptidases, loading onto HLA and presentation at the cell surface. Each cell’s HLAs present multiple different peptides at varying peptide-HLA copy numbers per cell. HLA-II genes (HLA-DR, HLA-DP and HLA-DQ) are constitutively expressed in only a subset of cells specialized for antigen presentation, such as dendritic cells, B cells, and macrophages, but expression can also be induced in additional cell types, e.g. in response to cytokine stimulation. HLA-II molecules present peptides derived from extracellular proteins taken into cells via endocytosis and phagocytosis, and intracellular proteins that access the HLA-II processing pathway via autophagy.

HLA-I molecules typically bind peptides of 8-12 amino acids (aa) in length. The HLA-I peptide-binding cleft is closed at both N-and C-terminal ends, and optimal length preferences are often biased towards binding of 9-mer peptides; longer peptides frequently bulge out of the cleft to be accommodated (2). For most HLA-I alleles the most abundant peptide length is 9 aa, but fine length preferences differ between alleles -in particular, some bind almost exclusively 8-and 9-mers (e.g., HLA-B* 51:01) while others have a relatively high frequency of ligands of length 12-13 aa (e.g. HLA-A* 01:01) (3). By contrast, HLA-II molecules possess an ‘open’ peptide-binding cleft and can therefore accommodate longer peptides than HLA-I. They frequently present nested sets of peptides that have a common “core” with N-and C-terminal extensions of varying length (4). High affinity ligands for a given HLA allele usually share a common amino acid motif with relatively strict preferences in anchor positions (for HLA-I usually the second (P2) and last (PΩ), for HLA-II -P1, P4, P6 and P9), which form specific interactions with residues of corresponding HLA binding pockets (4, 5). The HLA locus is the most polymorphic in the human genome with tens of thousands alleles described to date (6). HLA variants that differ in peptide-contacting residues differ in the repertoire of peptides they present. The diversity of HLA alleles in the population is an important evolutionary mechanism for defense against diverse pathogens, e.g. rapidly mutating viruses. Different HLA alleles are associated with the severity and outcomes of viral infections. For example, the HLA-C* 15:02 allele is associated with protection against SARS-CoV-1 (7), and HLA-B57 is highly associated with efficient HIV-1 control and long-term non-progressive infection in the absence of antiretroviral therapy (8).

Large-scale *in vitro* binding assays and recent advances in mass spectrometry (MS) have enabled generation of large datasets of ligands for many HLA alleles (5). The largest database of HLA ligands, IEDB (9), contains ∼750,000 peptidic epitopes presented by 830 MHC alleles (as of August 2020).

Experimental HLA ligandome data is used for the training of artificial neural networks for prediction of HLA ligands and T cell epitopes (reviewed in (10, 11)). Different tools, such as NetMHCpan (12) and MHCflurry (13) allow HLA-I ligand prediction with high accuracy and allow predictions even for HLA variants with no experimental data available (14). For HLA-II predictions of peptide binding are complicated by substantial variation in length of presented peptides and currently available HLA-II binding predictors have limited accuracy (4).

Comparison of MS-eluted HLA ligands and decoys predicted as HLA binders that were not observed in MS data enables the development of antigen processing predictors (13). The combination of antigen processing and HLA binding predictors in the MHCflurry tool resulted in significantly higher performance compared to HLA binding prediction only (13). Experimental HLA ligandome data (15) is also useful for the investigation of properties of proteins that are more likely to give rise to HLA ligands. It was recently shown that helical regions are significantly enriched in the ligands, suggesting different proteolytic resistance depending on the secondary structure and size of the initial protein fragment (16).

Apart from that, protein length and expression level, rate of proteasomal degradation, mRNA translation efficacy, presence of proteolytic signals, and sites of ubiquitination also influence the presence of protein-derived peptides in HLA ligandome (17, 18).

Several studies have employed gene ontology (GO) analysis to characterize functions of proteins that frequently serve as HLA ligands sources (17–23). These studies found enrichment of mitochondrion, ribosome, and nucleosome cellular component terms (17) and DNA-, RNA-and protein-interaction molecular function terms (18) and relative depletion of membrane and extracellular matrix proteins (21). However, these results may reflect differences in the expression level of the corresponding genes, rather than enrichment of HLA ligands within them. Abelin *et al*. (24) demonstrated that after correction for expression, enrichment in HLA ligands is observed only for proteins associated with the late endosome, although in the absence of the correction proteins with other localizations were also enriched (ER, mitochondria, nucleus, secreted) or depleted (cell membrane, cytoplasm). It was recently shown that most foreign MHC-I-displayed peptides are immunogenic (25). Additionally, recent work by the Cerundolo lab suggests that mitochondria-localized proteins are more immunogenic than other human peptides (26), which has implications for cancer immunotherapy. However, the studies mentioned above were based on aggregated datasets containing ligands from many distinct HLA alleles, and corresponding analyses were not focused on exploring differences between alleles. The datasets in question were also relatively small, e.g. in the largest of them (18) only 59% of human genes gave rise to at least one HLA ligand, while as it was recently shown by Sarkizova *et al*. (3) all human proteins may serve as sources of HLA ligands.

In this study, we investigated HLA binding preferences in terms of functions of presenting genes. In order to remove gene expression biases and focus on HLA presentation only, we used HLA ligandomes predicted *in silico* by the commonly used tool NetMHCpan-4.0, and focused mostly on HLA-I alleles due to significantly lower accuracy prediction for HLA-II. We performed HLA binding predictions for all possible peptides of length 8-12 derived from the human proteome for a set of HLA alleles with different binding motifs.

For all protein-coding genes, enrichment in HLA ligands was computed, and GO enrichment analysis was performed for sets of genes depleted or enriched in HLA ligands. Our results demonstrate that HLA alleles have a tendency to present peptides derived from proteins with specific molecular functions. These propensities are different for HLAs with different binding motifs, but similar for alleles with similar anchor residue preferences, which is explained by HLA preferential presentation of proteins enriched in amino acids that are favourable anchor residues for that allele.

Using experimental data from the HLA ligand atlas (15), we observe substantial differences between HLA class I and class II alleles, with class I alleles tending to present intracellular proteins and class II -membrane transport proteins.

Differences in functions of proteins preferentially presented by different HLA variants may be important for antiviral immunity. We demonstrate that HIV-protective HLA-B* 57:01 is more likely to present proteins from GO categories corresponding to viral genes as compared with non-protective HLA-B* 08:01.

We also hypothesized that HLA presentation bias towards proteins with specific functions may be compensated for in haplotypes. We demonstrate that HLA-A/HLA-B and HLA-A/HLA-C allele pairs from frequent HLA haplotypes are significantly more different in their GO enrichment profiles of the presented proteins than random allele pairs.

## Methods

### Predicting HLA ligands

All possible 8-12 mers were cut from the human proteome and supplied to the software NetMHCpan v 4.0 (12) in order to predict putative HLA ligands. A proof-of-concept analysis was initially performed in this study for an *ad hoc* selected set of 6 HLA alleles (HLA-A* 02:01, HLA-A* 11:01, HLA-B* 07:02, HLA-B* 27:05, HLA-C* 02:02, HLA-C* 15:02), all having different anchor residue preferences (**Supplementary Figure S1**).

For exploration of differences in presentation of viral genes we expanded this set to 12 alleles with the addition of HLA-A* 01:01, HLA-A* 03:01, HLA-B* 08:01, HLA-B* 57:01, HLA-C* 07:02 and HLA-C* 08:01. Viral HLA-binding peptides were predicted using NetMHCpan v 4.0 software in the same way as human-derived ligands.

Conclusions from the smaller dataset were supported upon repetition of analysis using an extended set of 93 HLA alleles (see **Supplementary Table 3**) covering 95% of individuals worldwide. For analysis of compensation of HLA presentation bias in haplotypes, ligandome predictions were also made for alleles corresponding to frequent haplotypes but not presented in the set of 93 alleles. In total, 133 HLA alleles were surveyed in the current study.

The NetMHCpan software was run using default parameters, and both strong and weak binders (Rank < 2) were used as the list of putative human-derived ligands for each allele. Complementary analysis was performed using the MHCflurry software (13) with default parameters. In order to confirm that results were not biased by using a specific HLA binding prediction algorithm we selected HLA ligands based on the “Affinity percentile” column from MHCflurry. We also used the “Presentation score” column to account for antigen processing biases.

### HLA ligand enrichment analysis

Human proteins were assayed based on the number of observed and expected ligands for each HLA allele as follows. We first counted the number of predicted ligands *N*_*i*_ of length *l* coming from protein *i*. The average number of presented ligands for each HLA allele was computed as *ρ* =< *N*_*i*_ >/< *L*_*i*_ >, where *L*_*i*_ = *length of protein* − *l* is the corrected protein length and <·> denotes the average over the proteome. The probability of observing a given number of ligands from each gene and the odds are computed using Binomial distribution as *P*(*N*_*i*_) = *P*_*binom*_(*N*_*i*_|*ρ, L*_*i*_) and *log Odds* = *log* (*N*_*i*_ / *ρL*_*i*_). These values were used to define sets of HLA ligand-enriched and -depleted proteins (HLEPs and HLDPs).

### Experimentally validated HLA ligands

HLA ligands for both class I and class II alleles were extracted from the HLA Ligand Atlas dataset (15) that lists peptides obtained from publicly available MS HLA elution experimental data.

### Mass spectrometry-based profiling of peptides presented on single HLA-I allele-expressing cell lines

HLA-I-deficient CD4-expressing 721.221 cells (originally obtained from Prof Masafumi Takaguchi, Kumamoto University, Japan) were stably transfected with HLA-A* 02:01. Transfectants were expanded by growth in RPMI 1640 medium (Thermo Fisher) containing 10% fetal bovine serum (FBS), 2 mM L-glutamine, 100 U/mL penicillin, and 100 μg/mL streptomycin (R10), and 2 x 10^8^ cells harvested for HLA-I bound peptide profiling. Mass spectrometry-based immunopeptidome profiling of HLA-A* 11:01-transfected CD4.221 cells was reported previously (27); the same methodology was employed here for immunopeptidome profiling of HLA-A* 02:01-expressing CD4.221 cells.

Briefly, cells were washed once in PBS, pelleted and 1 ml IGEPAL buffer [0.5% IGEPAL 630, 50 mM Tris pH8.0, 150 mM NaCl and 1 tablet Complete Protease Inhibitor Cocktail EDTA-free (Roche) per 10 ml buffer] was added per 0.5–1 × 10^8^ cells, and cells were lysed by mixing for 45 min at 4°C. Cell lysates were cleared by two centrifugation steps, 2000 × g for 10 min followed by 20,000 × g for 30 min at 4°C. HLA-peptide complexes were immunoprecipitated from the cell lysates on W6/32-coated Protein A-Sepharose beads overnight at 4°C. W6/32-bound HLA-peptide complexes were sequentially washed with 20 mL of wash buffer 1 (0.005% IGEPAL, 50 mM Tris pH 8.0, 150 mM NaCl, 5 mM EDTA), wash buffer 2 (50 mM Tris pH 8.0, 150 mM NaCl), wash buffer 3 (50 mM Tris pH 8.0, 400 mM NaCl) and finally wash buffer 4 (50 mM Tris pH 8.0). Peptide-HLA complexes were eluted from the beads in 5 mL of 10% acetic acid, and samples were dried down prior to resuspension in 120 μL loading buffer (0.1% TFA, 1% acetonitrile in ultragrade HPLC water). Samples were loaded onto a 4.6 × 50 mm ProSwift RP-1S column (Thermo Fisher Scientific) and eluted using a 500 μL/min flow rate over 10 min from 2 to 34% buffer B (0.1% TFA in acetonitrile) in buffer A (0.1% TFA in water) using an Ultimate 3000 HPLC system (Thermo Scientific). Alternate odd and even HPLC fractions were pooled and dried down prior to resuspension in 20 μL LC-MS/MS loading buffer (0.1% TFA in water).

For LC-MS/MS analysis, 9 ul of each sample was injected onto a Dionex Nano-Trap precolumn (Thermo Scientific), before separation with a 60 min linear gradient of acetonitrile in water of 2-25% across a 75 µm × 50 cm PepMap RSLC C18 EasySpray column (Thermo Scientific) at 40°C and a flow rate of 250 nl/min, resulting in an approximate average pressure of 600 bar. LC solvents contained 1%(v/v) DMSO and 0.1%(v/v) formic acid. Peptides were introduced using an Easy-Spray source at 2000V at to a Fusion Lumos mass spectrometer (Thermo Scientific). The ion transfer tube temperature was set to 305°C. Full MS spectra were recorded from 300-1500 m/z in the Orbitrap at 120,000 resolution with an AGC target of 400,000. Precursor selection was performed using TopSpeed mode at a cycle time of 2 s. Peptide ions with a positive charge between 1-4were isolated using an isolation width of 1.2 amu and trapped at a maximal injection time of 120 ms with an AGC target of 300,000. Singly charged ions were deprioritised to other ion species during acquisition. Higher-energy collisional dissociation (HCD) fragmentation was induced and fragments were analysed in the Orbitrap. LC-MS/MS data was analysed using PEAKS v8.0 (Bioinformatic Solutions) software.

### Gene ontology enrichment analysis

Sets of proteins enriched and depleted in HLA ligands (HLEPs and HLDPs) were assayed for over-representation of certain Gene Ontology (GO) categories as follows. GO enrichment test was performed using GOANA method from Limma R package (28) and top enriched GO terms coming from molecular function (MF), biological process (BP) and cellular component (CC) were selected for visualization. Additional verification of GO enrichment trends was performed with DAVID web tool (29). Note that while sets of HLEPs and HLDPs were used for GO analysis for *in silico* predicted ligands datasets, sets of proteins containing at least one HLA ligand were assayed for experimental data as most of those datasets contain too few ligands to perform ligand enrichment test.

### Analysis of compensation for HLA presentation bias in haplotypes

Data for HLA-A/HLA-B/HLA-C haplotypes with the highest frequency in 19 populations of different ethnic origin (listed in **Supplementary Table 4**, all from USA National Marrow Donor Program) was taken from the “Allele frequencies” database (http://www.allelefrequencies.net/) (30). Filtering for haplotypes with a frequency higher than 0.01% resulted in multiple entries for each of the populations (mean 42, range 28 −89) which were merged to an aggregated dataset of 806 HLA-A/HLA-B/HLA-C haplotypes.

Further, these haplotypes were split into pairs of corresponding HLA-A/HLA-B, HLA-A/HLA-C, and HLA-B/HLA-C alleles (the resulting dataset is referred to as “Haplotypes”). The “Control” dataset included all possible HLA-A/HLA-B, HLA-A/HLA-C, and HLA-B/HLA-C combinations of alleles from the “Haplotypes” dataset with the exclusion of those pairs that were identical to pairs from the “Haplotypes” dataset within the first two digits in allele names.

For each pair of alleles, the euclidean distance between GO term enrichment profiles was calculated and the distributions of that distance for “Haplotypes” and “Control” datasets were compared. GO term enrichment profile is a vector composed of enrichment folds for each of the analyzed GO terms (wherein fold value was taken with a positive sign for enriched GO terms, for depleted terms with a negative sign, and for not significantly changed terms fold was set to 0).

### Data analysis code availability

Custom R scripts were used for data processing, analysis and plotting. R code used for the analysis is stored as an R markdown notebook that was tested under R v3.6.2. Scripts and smaller datasets are available at corresponding GitHub repository https://github.com/antigenomics/hla-go-ms.

## Results

### Exploring differences in HLA ligand incidence across human proteins

We started our analysis (see **Figure 1** for overview) by running a large-scale *in silico* prediction of HLA ligands in the entire human proteome using NetMHCpan software. For our exploratory analysis we predicted 9-mer ligands for an *ad hoc* set of 6 HLA alleles: two HLA-A alleles (HLA-A* 02:01 and HLA-A* 11:01), two HLA-B alleles (HLA-B* 07:02 and HLA-B* 27:05) and two HLA-C alleles (HLA-C* 02:02 and HLA-C* 15:02). Our predictions yielded ∼5×10^5^ HLA ligands for each allele (**Supplementary Table S1**) in line with previous estimates of the number of 9-mers a single HLA can present (31).

**Figure 1.**
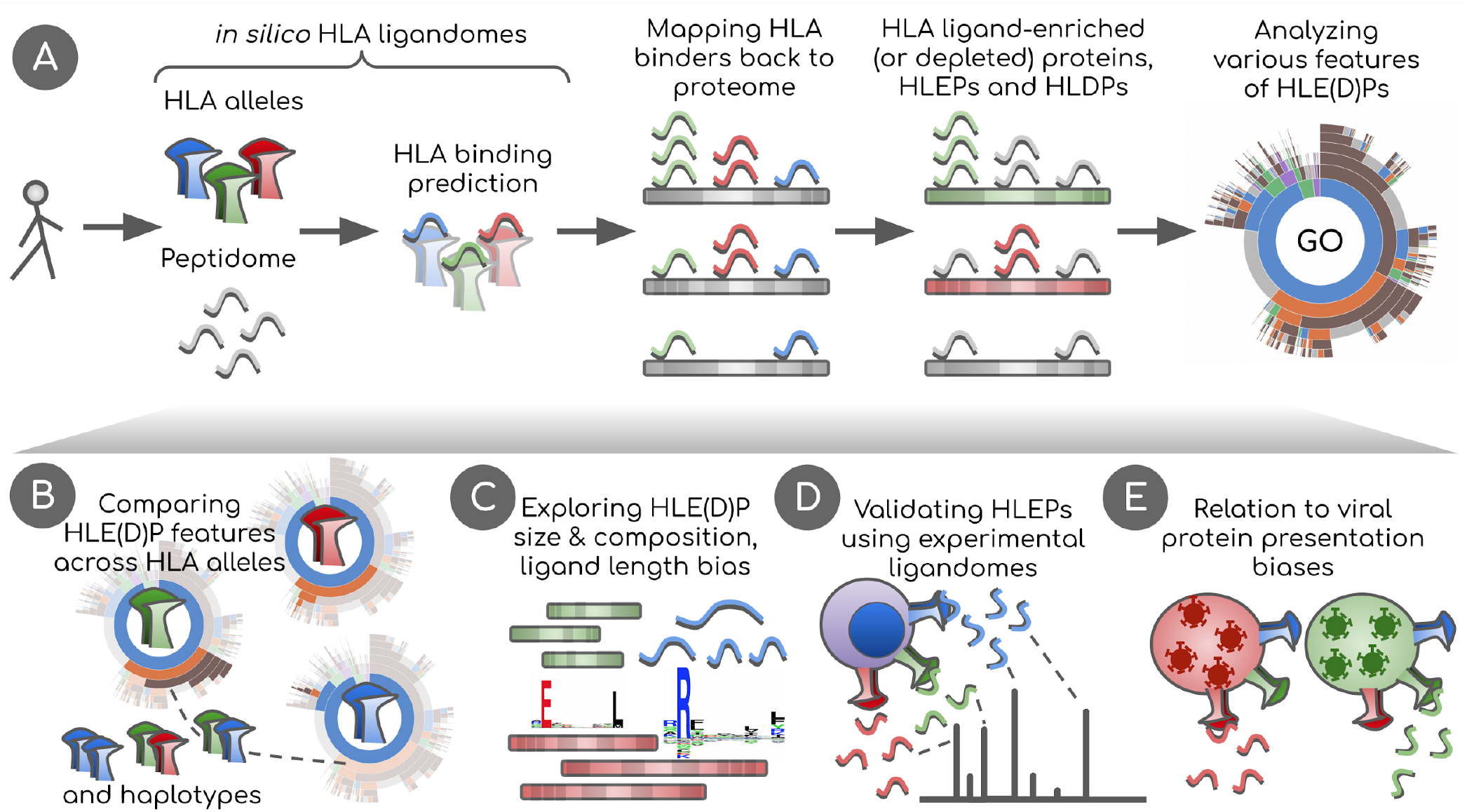
Overview of the study. **A**. *In silico* HLA ligandomes are generated by running HLA binding prediction software for the human peptidome (8-to 12-mers). HLA binders are then mapped back onto their parent proteins. Statistical analysis is performed to define sets of human proteins enriched or depleted in HLA ligands, HLEPs and HLDPs respectively. Functional analysis of these gene sets is performed using Gene Ontology (GO) category enrichment tests. **B**. GO annotation results are used to perform comparative analyses of HLA alleles, defining characteristic features of HLE(D)Ps. We show that preferred GO categories are clearly distinct between HLA alleles defining groups alleles with specific GO annotation profiles. These differences are however balanced and compensated by non-random selection of HLAs in HLA haplotypes observed in the population. **C**. Potential biases that shape HLE(D)P sets are explored, such as protein length, protein amino acid composition, together with the length of HLA ligands and HLA anchor residue types. **D**. Results are validated using real HLA ligandomes obtained from mass spectrometry data. **E**. Differences are identified in non-self peptide presentation by various HLAs by studying HLA presentation preferences of viral proteins. We link viral and human peptide presentation by demonstrating the relation between self-and non-self protein presentation preferences for various HLAs.

In order to identify proteins in which ligands are either enriched (HLA ligand-enriched proteins, HLEPs) or depleted (HLDPs) for every surveyed HLA allele we computed the number of ligands mapping to every human protein and estimated the expected number of ligands for each protein as the proteome-average ligands-per-amino acid frequency multiplied by the length of the protein. HLEPs and HLDPs were then selected based on a fixed P-value threshold (computed using Binomial distribution and adjusted for multiple testing, see **Materials and Methods**) and the observed-to-expected ratio of mapped ligand counts (**Figure 2A**). For all HLA alleles surveyed we observed many HLEPs (an average of 877 across all alleles) and HLDPs (565 on average), but the exact number of such proteins varied greatly across alleles (range from 55 to 1946 for HLEPs and from 106 to 1557 for HLDPs, see **Supplementary Table S1**).

**Figure 2.**
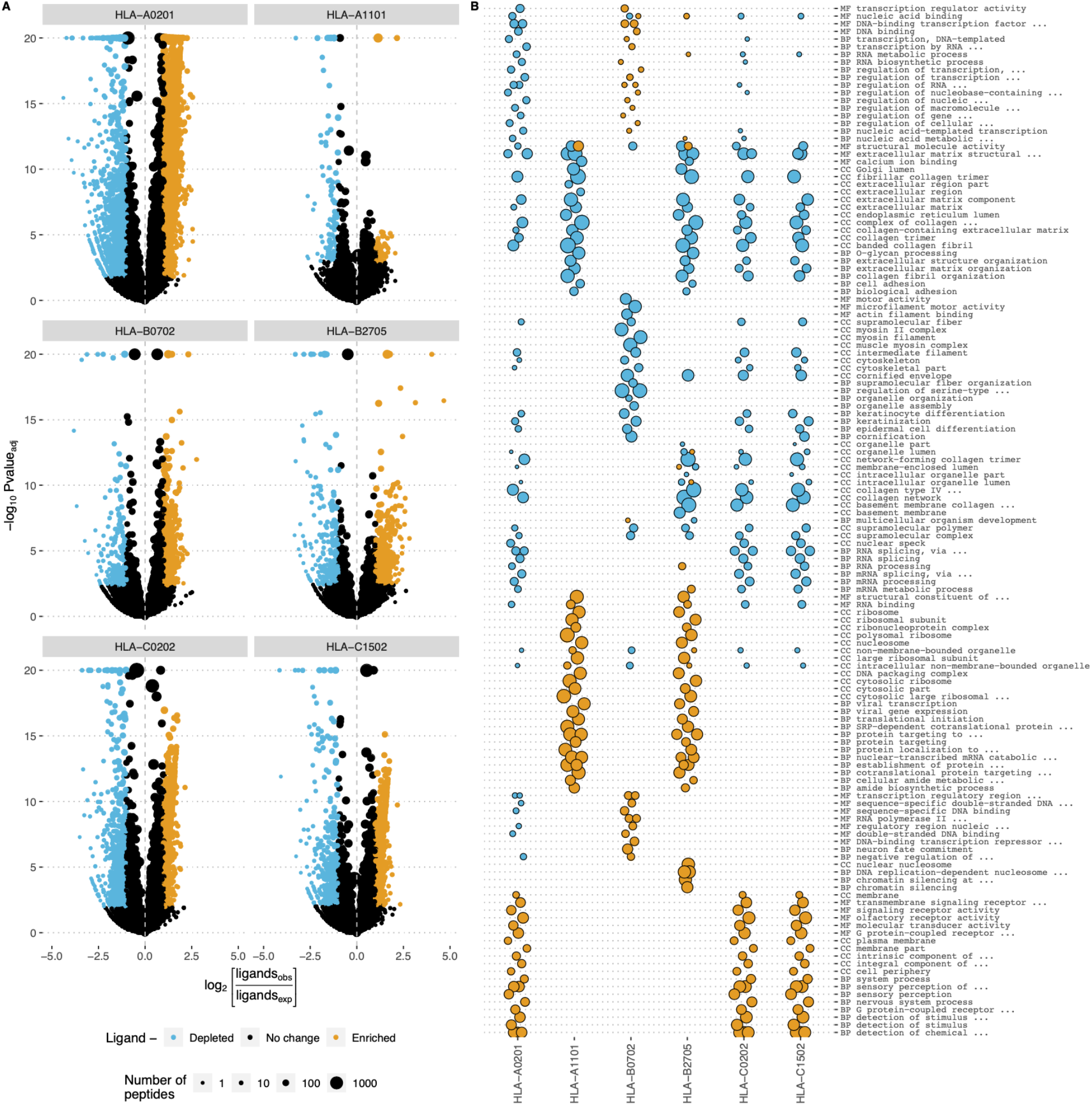
Human genes enriched and depleted in HLA ligands and their associated Gene Ontology (GO) categories. **A**. Volcano plots showing the log of the ratio of observed and expected number of HLA ligands for each human gene plotted against enrichment P-value computed using binomial test. Point size shows number of predicted HLA ligands, point colour highlights genes enriched and depleted in ligands according to at least 2-fold increase or decrease in the number of ligands and adjusted P-value of < 0.05. Data for 6 selected HLA alleles are shown as separate plots. **B**. GO term enrichment analysis for human genes differentially presented by different HLAs. Point size represents the GO enrichment fold for genes enriched (yellow) and depleted (blue) in HLA ligands for each of 6 surveyed HLA alleles. An adjusted P-value threshold of 0.01 was used as a threshold, Y axis lists the union of sets of top 20 GO categories for both ligand-enriched and ligand-depleted genes for each HLA allele. GO term names are preceded by either CC (cellular component), MF (molecular function) or BP (biological process) ontology name.

### HLAs differentially present human proteins associated with certain gene ontology categories

We next analyzed the length distribution HLEPs, HLDPs and the remaining proteins that do not show any difference in HLA ligand counts (**Supplementary Figure S2**). While one might expect that longer proteins would provide more statistical power to infer differences in the number of HLA ligands, we found that proteins of any length can feature differences in HLA presentation. More specifically, we observed that longer proteins are more likely to be depleted in ligands while shorter ones are enriched in presented peptides.

In order to characterize human genes encoding proteins that are more or less likely to be presented by HLAs we ran a gene ontology (GO) term enrichment analysis as shown in **Figure 2B**. We observed prominent enrichment of certain “molecular function” (MF), “biological process” (BP) and “cellular component” (CC) GO categories of genes coding for HLEPs and HLDPs, and found differences between GO category profiles across surveyed HLAs. In general, genes coding for HLDPs are more likely to encode extracellular matrix components, collagen and myosin, which is in line with the observation mentioned above as those are typically longer proteins. It is thus hard to decouple potential length bias from gene function in this case, as all HLA alleles show similar disfavoring of this set of genes.

On the other hand, genes coding for HLEPs display a diverse set of associated GO categories. For example, HLA-A* 02:01, HLA-C* 02:02 and HLA-C* 15:02 are more likely to present ligands from genes encoding membrane proteins and those involved in receptor signalling such as G-coupled receptor and olfactory receptor signalling. HLEPs for HLA-A* 11:01 and HLA-B* 27:05 are involved in translation and gene expression, while HLA-B* 07:02 ligands derive from proteins involved in regulation of transcription and HLA-B* 27:05 presents ligands from genes involved in DNA replication and chromatin silencing. It is also necessary to note that HLA-A, -B and -C genes do not show much similarity and alleles of different HLA-I genes can have similar preferences.

### Human proteins are differentially represented within HLA ligands of different length

In order to check for differences in human HLEP and HLDP set composition for HLA ligandomes corresponding to different peptide lengths we surveyed 8-mer to 12-mer predictions for the HLA-A* 11:01 allele as described above (**Supplementary Figure S3**). HLA-A* 11:01 predominantly presents 9-and 10-mers, although longer and shorter peptides are also known to be presented by this allele (32). We found genes that were either enriched or depleted for HLA ligands for all surveyed peptide lengths; the total number of ligands of each length was around 10^5^ (**Supplementary Table S2**). Analysis of genes enriched within HLA-A* 11:01 ligands of each length reveals a number of GO categories that are associated with longer and shorter HLA ligands (**Supplementary Figure S4**). GO categories characteristic of genes depleted in HLA-A* 11:01 ligands are similar across all ligand lengths and correspond to genes coding for extracellular proteins and collagen, in line with general trends observed for the 6 HLA-I alleles described above. GO categories of genes coding for HLA-A* 11:01 HLEPs are, however, distinct across peptide lengths: 9-and 10-mer peptides are linked to genes involved in transcription and translation processes, while 12-mers are linked to genes associated with mitochondrial and transporter genes. As observed biases in HLEP features may be due to differences in *in silico* ligand prediction accuracy for different lengths (for example, there are far more training examples of 9-mer ligands than 11-mers (12)), we performed additional validation of these results using experimental HLA ligandomes and alternative software tools as described in the next section.

### Amino acid composition of proteins enriched or depleted in HLA ligands

To explore the molecular basis for differences in gene presentation profiles between HLA alleles, we compared the amino acid composition of HLEPs and HLDPs. The results presented in **Figure 3A** demonstrate that HLEPs are enriched in amino acids which are good anchor residues for the particular HLA allele (see **Supplementary Figure S1** for motifs of presented peptides). For example, for HLA-A* 02:01, HLA-C* 02:02, and HLA-C* 15:02, which require hydrophobic anchors, HLEPs have a higher frequency of hydrophobic and lower frequency of charged residues. The amino acid frequency profile for HLA-B* 27:05 HLEPs is very close to that of the human proteome except for a higher frequency of arginine, which is strictly preferred by the allele as an anchor residue in the P2 position.

**Figure 3.**
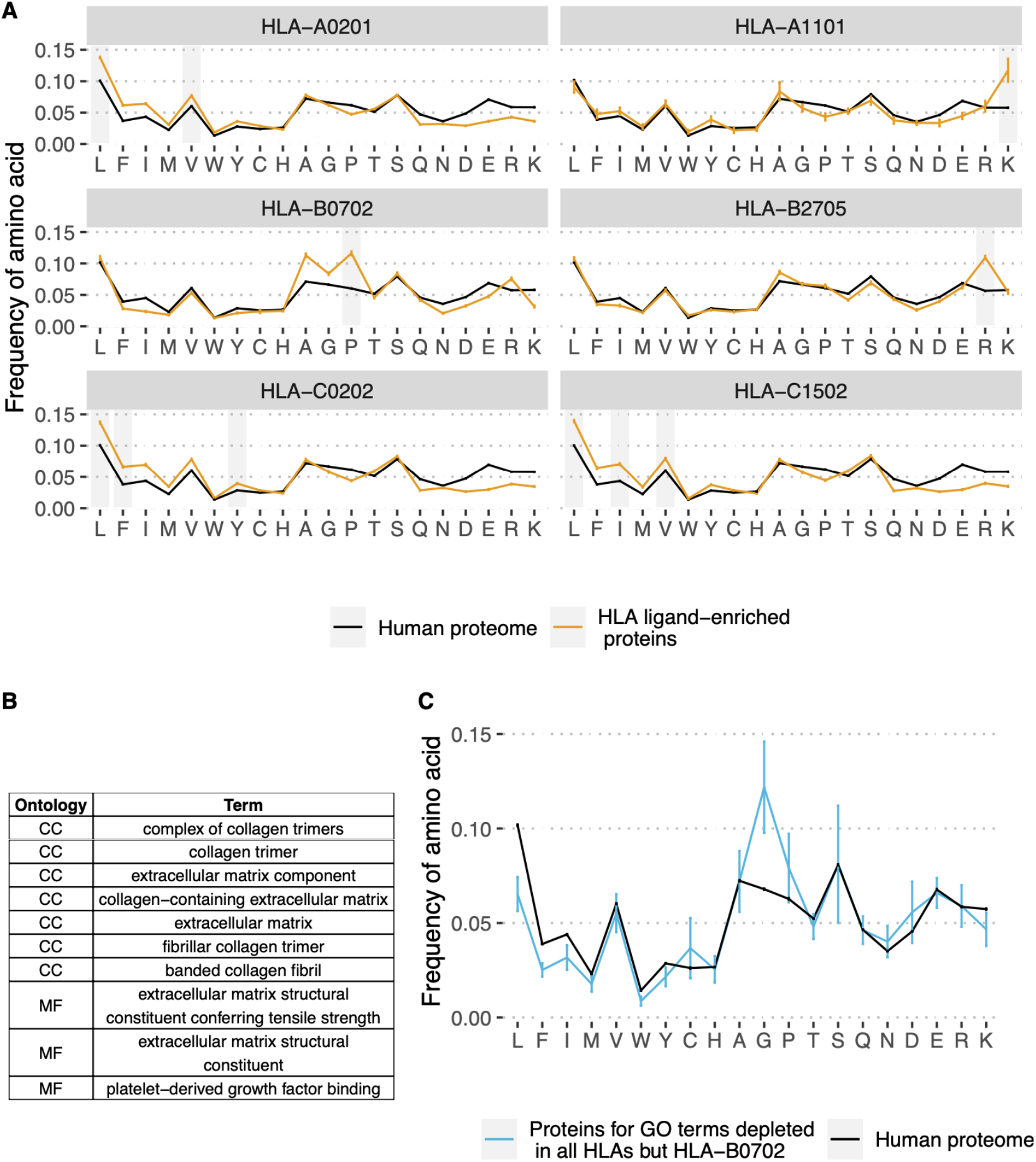
Amino acid composition of HLEPs and HLDPs for different HLA alleles. **A**. Comparison of amino acid composition of HLEPs for the 6 selected HLA alleles and all proteins of the human proteome. Grey bars mark amino acids preferred by the allele in the anchor positions (P2 and/or P9) according to **Supplementary Figure 1**. Note that HLEPs tend to have a higher frequency of amino acids that are good anchors for this allele. Error bars show 95% confidence interval for the mean value. **B**. GO categories associated with HLDPs for 5 of 6 selected alleles (all but HLA-B* 07:02). Ontology names: CC -cellular component, MF -molecular function, BP -biological process. **C**. Comparison of amino acid composition of proteins corresponding to GO categories from **B**. (only proteins which are in HLDPs for at least 1 allele were considered) and human proteome. Error bars show 95% confidence intervals for the mean value. Amino acids which are enriched in proteins corresponding to “commonly depleted” GO categories are glycine which can’t be used as anchors for most HLA alleles, and proline which dramatically affect peptide backbone conformation. HLA-B* 07:02 represents a special case as this allele strictly prefers proline as anchor residue in P2.

The observed bias in the amino acid composition of HLEPs and HLDPs leads to differences in the GO categories enriched in these gene sets. Thus, for HLA-A* 02:01, HLA-C* 02:02 and HLA-C* 15:02, which are prone to present more hydrophobic proteins, GO categories enriched in HLEPs are mainly associated with membrane proteins (**Figure 2B**) which have a relatively high frequency of hydrophobic residues. HLA-A* 11:01 and HLA-B* 27:05, which require lysine and arginine as anchor residues, are more likely to present proteins involved in interaction with DNA (**Figure 2B**) and that have a relatively high frequency of positively charged amino acids.

Comparing GO enrichment analysis results for different alleles we found that several GO categories are enriched in HLDPs for all surveyed alleles except HLA-B* 07:02 (**Figure 2B, Figure 3B**). Genes corresponding to these GO terms are enriched in glycine and proline residues as shown in **Figure 3C**. These GO categories are mostly associated with the extracellular matrix (**Figure 2B, Figure 3B**) and include fibrous proteins such as collagen. Glycine and proline residues are critically important for collagen ternary structure formation. Glycine is a bad anchor residue for almost all HLAs, and proline dramatically affects peptide conformation preventing its binding with HLA. It can be suggested that proteins enriched in G and/or P are poorly presented by multiple HLAs. HLA-B* 07:02 is the exception as this allele strictly requires proline as the peptide P2 anchor.

### Exploration of ligand presentation bias on an extended dataset of HLA alleles

To check that the conclusions from the analyses in the previous sections are not artifacts of the selection of surveyed HLA alleles, we performed *in silico* predictions of HLA ligands and GO enrichment analysis for an extended set of 93 alleles (**Supplementary Table S3)** using NetMHCpan software, in the same manner as for the initial set of 6 alleles. Alleles were selected to cover at least one of the HLA-A, HLA-B, and HLA-C alleles in 95% of individuals worldwide, based on allele frequencies reported by Sarkizova *et al*. (3).

First, we re-examined our observation of the existence of GO categories associated with HLDPs for most alleles. We observed that there is a group of GO terms related to extracellular matrix and collagen which are associated with HLDPs for up to 95% (89 out of 93) of surveyed alleles (**Supplementary Figure S5A**). Proteins corresponding to these terms are enriched in glycines and prolines (**Supplementary Figure S5C**). Exceptional alleles, for which these “universally depleted” terms are not associated with HLDPs are the ones that strictly require proline residues as P2 anchors (**Supplementary Figure S5B**). Overall, all the conclusions reached from the analyses performed with the initial set of 6 alleles, i.e. depletion of proteins enriched in glycines and prolines which correspond to GO terms related to structural functions (**Figure 3BC**, with the exception of HLA-B* 07:02 having proline anchor residues), remain valid when a broader range of alleles are considered.

Further, we explored HLA presentation preferences for proteins of different lengths. To better understand the protein length bias for HLEPs and HLDPs we compared the amino acid composition of human proteins of different lengths. Preferences in the length of presented proteins may be explained by the differential amino acid composition of proteins from different length quartiles (**Supplementary Figure S6A**). As shown in the figure, the frequency of hydrophobic amino acids is highest for proteins in the second length quartile (Q2) and lowest for Q4 proteins. In accordance with this trend Q2 proteins constitute the highest fraction of HLEPs for HLA-A* 02:01, HLA-C* 02:02, and HLA-C* 15:02 alleles featuring hydrophobic anchor residues. For HLA-A* 11:01 and HLA-B* 27:05, which utilise positively charged anchor residues, Q1 proteins constitute more than half of HLEPs, in line with the observation that these proteins have the highest frequency of arginines and lysines. For the majority of alleles, the highest fraction of HLEPs is composed of smaller proteins from the second length quartile (Q2) and most HLDPs are longer proteins from Q4 (**Supplementary Figure S6B**). Hydrophobic amino acids which are required as anchor residues for the majority of alleles (see **Supplementary Table S3**) are enriched in Q2, so for these alleles the length distribution of HLEPs peaks in Q2. There are some exceptions where the distribution of length of HLEPs is not maximal in Q2 (**Supplementary Figure S6C)**, but these exceptions are also explained by the amino acid composition of proteins in different length quartiles. Alleles for which length distribution of HLEPs peaks in Q1 require positively charged anchor residues (arginine and lysine) which are enriched in Q1 proteins. Alleles that prefer to present Q4 proteins require negatively charged glutamic acid as an anchor which is enriched in Q4 proteins (**Supplementary Figure S6A**).

### Validation of HLA allele and ligand length biases

To ensure our observations are not an artifact of the HLA ligand prediction model used by NetMHCpan software, we re-ran the same analysis using an alternative software tool MHCflurry (13). In addition to peptide-HLA binding, MHCflurry also takes into account antigen processing and can provide the combined score (“Presentation score”), which performs better to predict HLA ligands. We performed the analysis using both affinity and presentation scores of MHCflurry to enable evaluation of the impact of software and potential antigen processing biases. In both cases the analysis revealed nearly identical results to **Figure 2** in terms of ligand enrichment scores and P-values, as well as the list of associated GO categories, as can be seen in **Supplementary Note 1, Figure SN1**. Also, the bias in the number of HLA ligands reported for proteins of different lengths (**Supplementary Figure S2**) holds true when applying MHCflurry as a HLA ligand prediction method (**Supplementary Note 1, Figure SN2**).

In addition, to determine whether the interpretation of our findings would be affected by use of an alternative GO annotation strategy, we re-annotated HLA ligand-enriched and -depleted gene sets using DAVID, another commonly used web tool (29). DAVID analysis showed clustering of GO categories enriched and depleted for genes of interest that was highly in line with the results described above (**Supplementary Note 1, Figure SN3**). Annotation clusters (groups of GO categories) identified by DAVID as being over-represented are also supportive of the general trends mentioned above, e.g. depletion of proteins representing extracellular matrix components, collagen and myosin in HLA ligands.

We also independently validated our results with experimentally-determined ligandomes of HLA-A* 02:01 and HLA-A* 11:01 alleles (see **Methods** section). Analysis of GO categories enriched in parent proteins of HLA-A* 02:01 peptides compared to HLA-A* 11:01-presented proteins and vice-versa revealed consistency with the *in silico* results reported above, and protein GO categories matched HLEPs of corresponding HLA alleles. As can be seen from **Figure 4A**, GO categories that are enriched in either HLA-A* 02:01 or HLA-A* 11:01 according to *in silico* data analysis are also more common in proteins that feature HLA ligands of corresponding alleles in experimental data, supporting the observed difference between functions of HLA-A* 02:01 and HLA-A* 11:01 HLEPs. Moreover, GO categories enriched in HLEPs of HLA-A* 11:01 allele ligands of various lengths are highly correlated with GO categories enriched for ligands of corresponding lengths in experimental data (**Figure 4B**).

**Figure 4.**
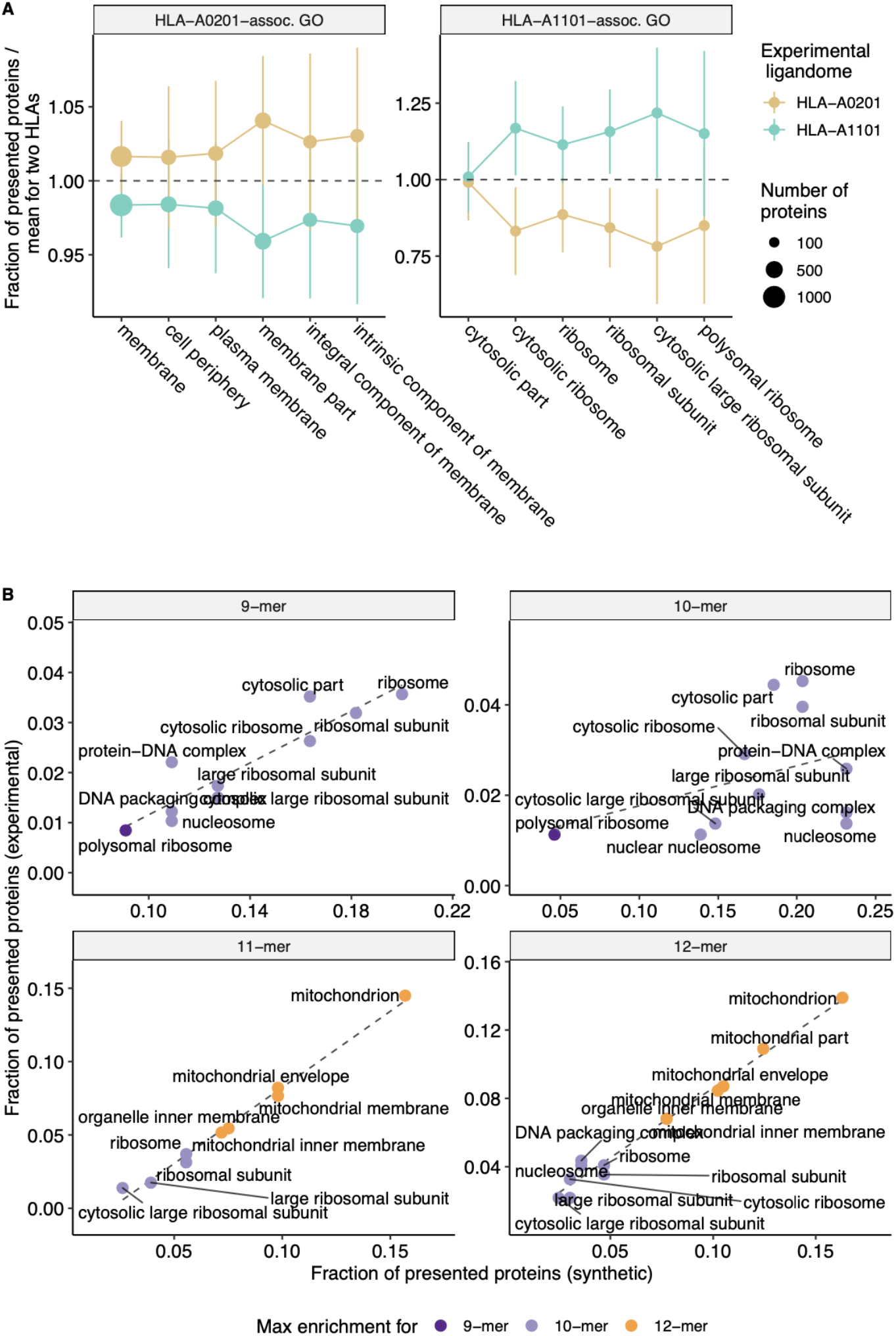
Experimental validation of biased selection of self proteins presented by different HLA alleles and different HLA ligand lengths. **A**. Fraction of proteins presented by experimentally obtained HLA-A* 02:01 and HLA-A* 11:01 ligandomes that correspond to a given GO category. The fraction is normalized by mean value for two alleles to highlight differences between proteins related to HLA-A* 02:01 and HLA-A* 11:01 alleles. Cellular component (CC) GO categories associated with proteins which are frequently presented by either HLA-A* 02:01 (left panel) or HLA-A* 11:01 (right panel) according to *in silico* predictions were selected to match those in **Figure 2B**. Error bars show 1 standard deviation of fractions. **B**. Scatterplot compares the fraction of proteins presented by HLA-A* 11:01 ligands of various lengths that correspond to a given GO category between *in silico* predictions and experimental data. CC GO categories associated with 9-12-mer ligands were selected to match those in **Supplementary Figure 5**. Colour shows ligand length at which maximum fold enrichment is reached for a given GO category.

### HLA ligand atlas analysis and difference between class I and class II HLAs

To provide further verification of our results on real HLA ligand datasets and compare them to *in silico* HLA ligand predictions we explored the HLA ligand atlas dataset (15). We ran GO enrichment analysis for sets of genes corresponding to peptides presented by each HLA allele in the dataset and compared profiles of enriched categories across alleles. Note that we used every gene that has at least one reported HLA ligand, and did not use an enrichment test for the number of ligands per gene as the size of the database is too small to ensure good coverage of all human genes. Using HLA ligand atlas allowed us to independently validate the phenomenon of preferential presentation of genes with specific functions by different HLA alleles.

Principal component analysis was used to visualize differences between HLAs based on functions of genes they tend to present as shown in **Figure 5A**. The plot shows clear separation between genes providing a source of ligands presented by HLA class I and class II, with co-clustering of HLA-C and HLA-B alleles and a notable differentiation between HLA-DQ versus HLA-DR. Notably, while HLA class I and class II alleles are clearly separable, it is hard to tell HLA class I genes apart based on GO enrichment profiles (**Figure 5B**).

**Figure 5.**
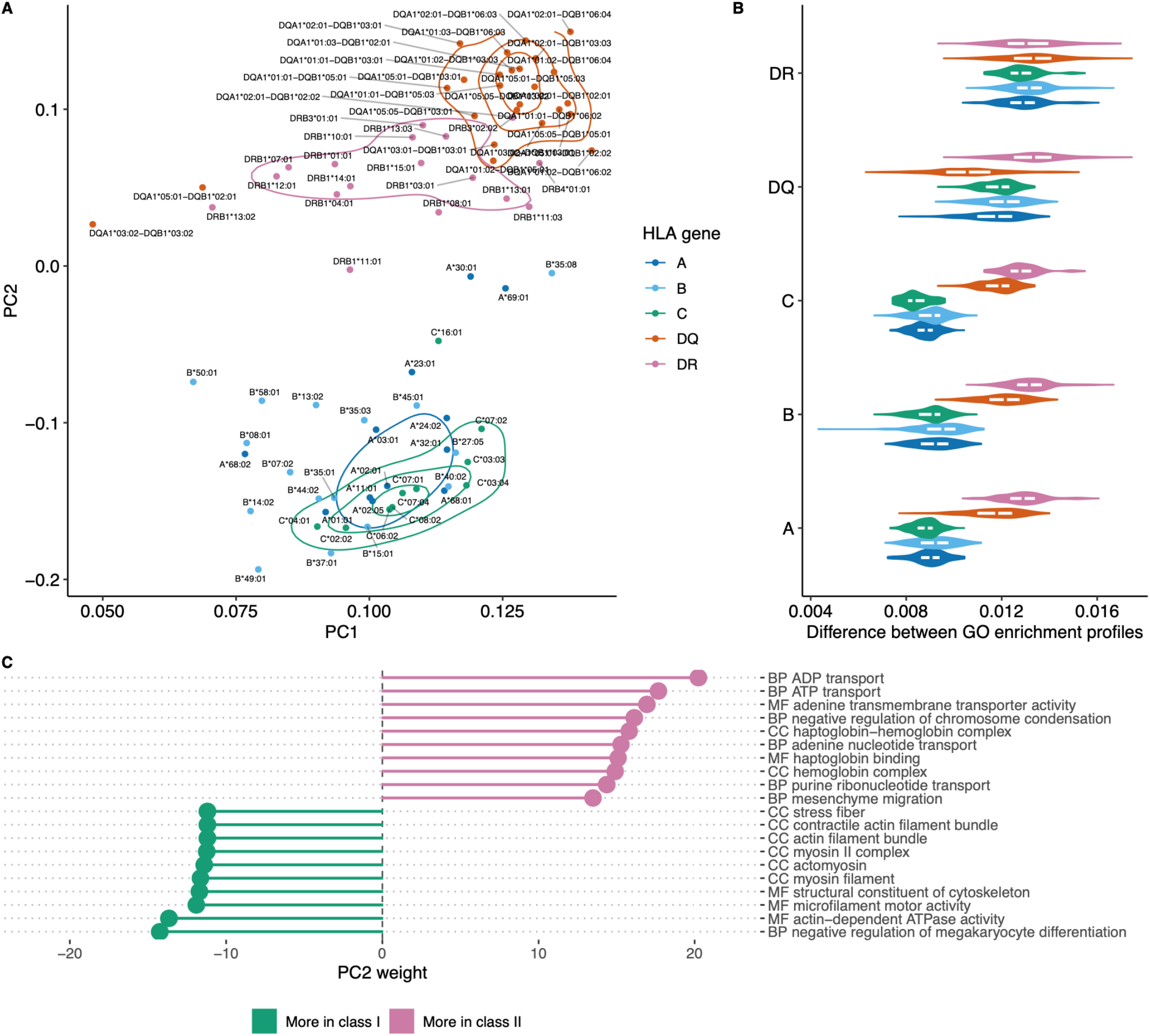
Visualizing similarities between HLA alleles based on enriched GO categories of genes they tend to present. **A**. Principal component analysis (PCA) results for GO category fold enrichment profiles of various HLAs. GO enrichment profiles are computed based on gene sets obtained by mapping HLA ligands from the HLA ligand atlas dataset as the logarithm of observed to expected fraction of genes representing a given GO category. Colour shows HLA gene: A/B/C for class I and DR/DQ for class II. **B**. Distribution of pairwise distances between GO enrichment profiles of HLA alleles of the same and distinct HLA genes. Y axis corresponds to the HLA gene of the first HLA allele in each pair, the gene of the second allele is indicated by colour (same as in **A**.). Euclidean distance divided by the total number of GO categories is used. **C**. List of the top 10 GO categories that have most absolute weight in PC2 (see panel **A**.). Negative weight corresponds to dominance in class I HLA alleles, while positive weight corresponds to dominance in HLA class II. GO term names are preceded by either CC (cellular component), MF (molecular function) or BP (biological process) ontology name.

For in-depth exploration of genes that are differentially presented between class I and class II alleles we took the PC2 component from **Figure 5A** that linearly separates HLA classes and analyzed GO categories having highest weights in these components (**Figure 5C**). We observed that class II HLA alleles are more likely to present membrane transport proteins, while class I alleles are prone to present components of intracellular structural proteins.

### Analyzing presentation of viral genes by different HLA alleles

As the spectrum of viral protein functions should be highly specific we suggest that our observation of differential presentation of peptides derived from human proteins with differing functions may be extrapolated to presentation of viral peptides by distinct HLAs. Thus, we suggested that there may be substantial differences in presention of viral peptides and different human HLA alleles will favor certain viral proteins. We calculated the presentation odds for each viral gene-HLA pair as the ratio of the observed number of ligands divided by expected value computed under the assumption of independence between an HLA allele and the number of ligands it presents from a given gene. Comparative analysis of presentation odds of viral peptides by human HLAs (**Figure 6A**) reveals co-clustering of viral genes with similar functions and certain HLAs (e.g. HLA-B* 07:02 and HLA-B* 27:05, HLA-A* 03:01 and HLA-A* 11:01).

**Figure 6.**
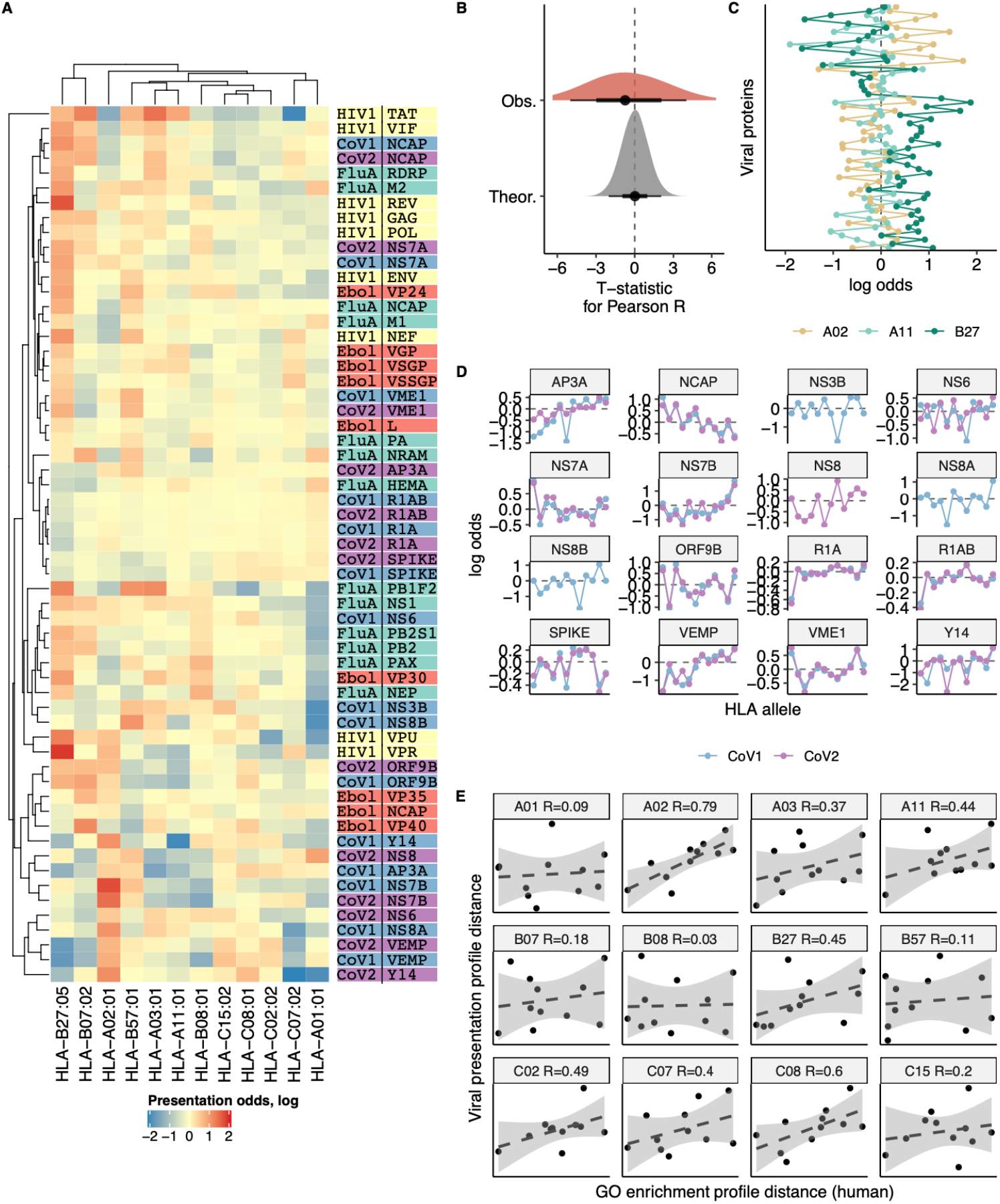
Differences in the number of ligands coming from viral genes presented by 12 different HLA alleles. **A**. Heatmap showing the logarithm of the ratio of observed to expected number of HLA ligands (presentation odds). Expected number of ligands for each gene and HLA pair was estimated as the sum of corresponding row and column of the matrix divided by the total number of ligands in the matrix. Dendrograms show results of hierarchical clustering of gene-wise and HLA-wise presentation profiles performed using complete linkage algorithm with Euclidean distance measure. Proteins from HIV (yellow), Influenza (blue) and SARS-CoV-1 (red) are shown. **B**. Absolute values of T-statistic for observed pairwise correlation coefficients between viral gene presentation profiles of different HLAs (dark grey) compared to theoretical distribution (n=1000 random samples from null distribution with same number of degrees of freedom, light grey). **C**. An example comparison of viral gene presentation profiles between HLA-A* 01 and HLA-A* 02 and HLA-B* 27 alleles. Two-tailed paired T-test yields T statistic of 3.3 and P-value of 0.002. **D**. Comparison of SARS-CoV-1 and SARS-CoV-2 protein presentation odds across different HLA alleles. **E**. Correlation between distance in viral protein presentation odds profiles (Y axis) and distance in GO category profiles of HLA ligand-enriched human genes (X axis) for all HLA alleles. Each panel shows distances from a given HLA profile to profiles of each of the remaining 11 HLA alleles. Allele name and Pearson correlation coefficient are shown in panel title.

However, surveyed HLA alleles mostly feature contrasting presentation odds profiles, and the distribution of correlation coefficients for these profiles is shifted to negative values (**Figure 6B**). For example a pair of HLA-A alleles, HLA-A* 02:01 and HLA-A* 11:01, appear to have distinct preferences for presenting viral proteins (**Figure 6C**), in line with their difference in preferences for presenting human proteins with certain functions reported above (**Figure 2B**). On the other hand, HLA-A* 11:01 and HLA-B* 27:05, which tend to present human proteins of similar functions (**Figure 2B**) are also similar in terms of viral protein presentation odds profiles (**Figure 6C**).

Considering similar viral proteins, we observe little difference in the way they are presented by the same HLA. When comparing presentation odds across all 12 surveyed HLAs between proteins of SARS-CoV-1 and SARS-CoV-2 strains we observe nearly perfect correlation for almost all proteins (**Figure 6D**).

Finally, we performed a direct comparison to test the hypothesis that tendency to present self-peptides with certain functions is intrinsically linked to the variability in viral protein presentation by HLAs. We tested if distance between self-peptide GO profiles was correlated with distance in viral presentation profiles, testing each of 12 HLA alleles against the remaining 11 (**Figure 6E**). All alleles show a positive correlation between these two distances, and for the majority the correlation was substantial (R > 0.3 for 7 out of 12 alleles). The overall correlation coefficient for all 66 possible distance pairs is R = 0.32, P = 0.009.

We can speculate that inter-allele differences in preferences for binding of peptides derived from viral as well as human proteins could be among the factors contributing to the differential association of particular alleles with protection/pathogenesis in different infections (see **Discussion**).

### Haplotype compensation of bias in HLA presentation of proteins with different molecular functions

In previous sections, it was noted that HLA alleles with similar anchor residue preferences have similar profiles of GO terms enrichment and viral gene presentation odds, while alleles with different anchor preferences are more likely to have contrasting profiles (see **Figure 2B** and **Figure 6ABC**). Considering that HLA alleles are inherited not individually but in haplotypes, one may hypothesize that haplotypes composed of HLA alleles that are prone to present proteins with different molecular functions might be evolutionarily advantageous as they would be able to present a more diverse set of peptides to the immune system.

To test this we collected a dataset of HLA class I haplotypes (combinations of HLA-A, HLA-B, and HLA-C alleles) which have the highest frequencies in populations of different ethnic origin (**Supplementary Table S3**, for details see Methods). Haplotypes were divided into pairs of alleles of different genes (HLA-A/HLA-B, HLA-A/HLA-C, and HLA-B/HLA-C). As a control set we reshuffled alleles from the haplotype set to make up random pairs. In both sets, for each resulting pair of HLA alleles, we computed GO terms enrichment profiles distance. A comparison of corresponding distributions (**Figure 7**) between these two sets demonstrates that pairs of HLA-A/HLA-B and HLA-A/HLA-C alleles associated with frequent haplotypes are significantly different in terms of the distance between corresponding GO terms enrichment profiles from control allele pairs. Thus, commonly observed haplotypes consist of more divergent pairs of HLA alleles in terms of the proteins they tend to present peptides from. This result may be interpreted as indicating that the haplotype composition is focused on compensating “holes” in the immunopeptidome that are the result of non-uniform proteome presentation by various HLAs.

**Figure 7.**
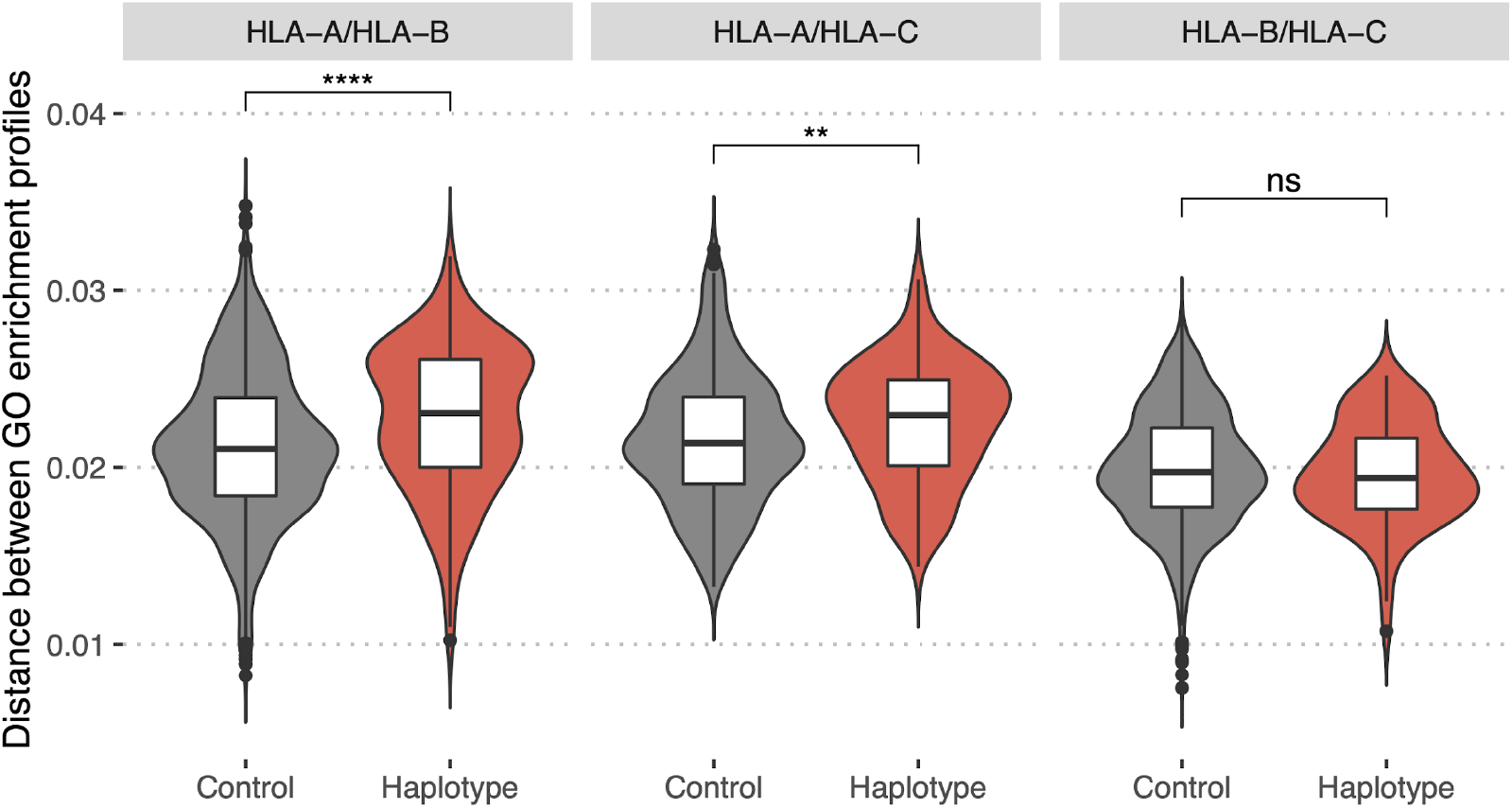
Bias in HLA presentation of proteins with different molecular functions is compensated in HLA-A/HLA-B and HLA-A/HLA-C haplotypes. Distance in GO category profiles between pairs of HLA alleles of different genes (HLA-A and HLA-B, left panel; HLA-A and HLA-C, central panel; HLA-B and HLA-C, right panel). “Haplotype” set is composed of pairs of alleles constituting haplotypes with high frequency in one of the populations, “Control” is composed of random pairs of alleles from the “Haplotype” set (for details see Methods section). Statistically significant differences between the groups are for HLA-A/HLA-B (p-value = 5e-07, Mann–Whitney U-test) and HLA-A/HLA-C genes (p-value = 0.004, Mann–Whitney U-test). Higher values of the distance for “Haplotype” group indicate that pairs of alleles composing frequent haplotypes tend to present proteins with distinct functions.

## Discussion

In this study, we performed a comparison of molecular functions of proteins preferentially presented by different HLA alleles. HLA restriction of peptide ligands shapes adaptive immune responses and is critical for protection against viruses and other pathogens, elimination of cancer cells and prevention of autoimmune diseases, moreover it directly shapes the repertoire of cognate T cells that form the backbone of adaptive immunity (1, 33–35). To investigate the effect of HLA restriction on the nature of the ligands presented to T cells, we studied peptides derived from the human proteome and predicted *in silico* binding for a set of HLA-I alleles having different recognition motifs. The number of predicted binders varied across HLA alleles in line with previous observations of different ligand length distributions (2) and sizes of experimental ligandomes of HLA-I variants (36). Ligand coverage of genes was not uniform -some genes gave rise to more or less predicted binders than was expected from the assumption of a random sampling of source genes. This difference in the number of ligands was used to select HLDPs and HLEPs, i.e. human proteins depleted and enriched in HLA ligands.

Enrichment in HLA-I ligands is biased towards proteins of shorter length: longer proteins constitute the largest fraction of HLDPs, while the HLEP set contains proteins of shorter lengths. This may be attributed to the different amino acid compositions of proteins of different lengths: shorter proteins have a higher frequency of hydrophobic, positively charged and tyrosine residues (which are favourable anchor residues for the surveyed HLAs) than longer ones. The observed length bias should also hold for other HLA-I alleles requiring hydrophobic, positively charged or tyrosine anchor residues, which constitute the majority of HLA-I alleles with a high frequency in the human population.

In order to comprehensively explore biases in features and functions of proteins preferentially presented by different HLA alleles we performed a gene ontology (GO) term enrichment analysis for sets of HLEPs and HLDPs. The profiles of GO categories corresponding to HLEPs and HLDPs differed between alleles but were similar for groups of alleles having similar anchor residue preferences (HLA-A* 02:01, HLA-C* 02:02, and HLA-C* 15:02; HLA-A* 11:01 and HLA-B* 27:05).

These patterns may account for preferences in HLA presentation of proteins with a higher fraction of amino acids which are good anchor residues for that particular allele. HLA-A* 02:01, HLA-C* 02:02, and HLA-C* 15:02 are more likely to present relatively hydrophobic proteins, thus GO terms corresponding to HLEPs for these alleles are mainly associated with membrane proteins. For HLA-A* 11:01 and HLA-B* 27:05 HLEPs have a higher fraction of positively charged amino acids and corresponding GO terms are associated with DNA binding functions. Some GO categories (e.g. associated with collagen) corresponded to HLDPs of all surveyed alleles but HLA-B* 07:02. Proteins constituting these categories have a relatively high proportion of glycine residues (which are disfavoured anchor residues for almost all HLA alleles) and prolines (which are conformationally rigid and may disturb peptide conformation suitable for HLA binding). Proline-rich proteins are also expected to be depleted in naturally processed HLA ligands for the reason that prolines are depleted in up-and downstream regions of proteasome cleavage sites (24). HLA-B* 07:02 is the exception as proline in P2 is strictly required for peptide binding by the allele variant.

In order to explore potential ligand length bias, for HLA-A* 11:01 we expanded our analysis for peptides of lengths 8-12 amino acids. Though general patterns of GO enrichment were similar for ligands of different lengths, some GO categories were associated with either shorter or longer ligands. This effect may be attributed to subtle differences in anchor residue preferences for ligands of different lengths, however such biases should be scrutinised to ensure that they are not an artifact of HLA binding prediction software.

Most of our initial analysis reported here was based on *in silico* HLA-I ligand predictions performed by NetMHCpan software and GO enrichment analysis based on a hypergeometric test. To exclude any bias coming from prediction methodology, we re-ran our analysis using an alternative algorithm, MHCflurry, and were able to reproduce our results. NetMHCpan software predicts only peptide-MHC binding and does not consider potential effects of proteasomal cleavage and TAP transport. To account for impact of antigen processing steps we reran our analysis using the “presentation score” feature of MHCflurry software, which is a combination of predictions of HLA binding and antigen processing and was demonstrated to perform better than either of the former when used individually for prediction of HLA-bound ligands (13). All the conclusions based on NetMHCpan predictions stayed true, indicating that antigen processing steps do not appear to exert significant additional biases on the HLA-I ligandome structure. Similarly, we tested an alternative GO enrichment analysis method implemented in the DAVID web tool, again arriving at similar results. Thus, our results appear to be robust and independent from the choice of bioinformatic tools. Moreover, our main result highlighting differences in functions of human proteins that are differentially presented by different HLAs also holds when using experimentally identified ligands from the HLA ligand atlas datasets and HLA-A* 02:01 and HLA-A* 11:01 ligandomes described in this study. HLA ligand prediction accuracy depends on the size of the training set, which in turn depends on the availability of experimental data, so predictions may be less accurate for some HLA alleles. In this study we observe similar groups of genes enriched and depleted in ligands both for extensively studied alleles (such as HLA-A* 02:01) and less studied ones (such as HLA-C* 15:02, which has a different binding motif to HLA-A* 02:01). Similarly, less preferred ligand lengths for the 6 HLA-alleles investigated in this study (8-, 11-and 12-mers) have lower binding prediction accuracy than more common ones (9-and 10-mers). We independently validated our results on human genes differentially presented by HLA ligands of different lengths using experimental data for the HLA-A* 11:01 ligandome. Thus, we suggest that our results should be robust to HLA ligand prediction accuracy differences.

It is necessary to note that in this study we did not consider potential effects of gene expression on HLA ligandomes. While gene expression is certainly associated with gene function and may alter gene ontology, the majority of our results were based on “expression-agnostic” proteomes (i.e. every protein was counted once independently of its expression). However, we believe that this factor deserves a more thorough investigation in follow-up studies. A recent study (26) reports that the subcellular localization of human proteins may influence HLA presentation and T cell priming in the case of cancer antigen-specific responses. Our results show that proteins with different cellular component (CC) gene ontology categories tend to be presented preferentially by different HLAs, further suggesting that CC-based biases in protein presentation may play a role in anti-tumor immune responses.

We also performed GO terms enrichment analysis for an experimental dataset from the HLA ligand atlas covering 51 HLA class I and 42 HLA class II alleles (15). The results obtained revealed co-clustering of various HLA class I and II alleles in terms of GO categories of genes they tend to present. Interestingly, PCA analysis shows a clear separation between HLA classes in terms of typically presented proteins, and HLA-DR alleles are clearly separated from HLA-DQ ones while there is little separation between HLA-A, -B and -C classes for HLA class I.

Different preferences in functions of presenting proteins between HLA alleles may be important for antiviral immunity. We examined the link between the presentation of proteins encoded by viral genes and self genes with specific functions. For all possible pairs of surveyed HLA alleles, we observed a substantial correlation between self-protein GO profiles and profiles of viral protein presentation (considering proteins from influenza virus, HIV-1 and SARS-CoV-1/2). We speculate that differences in the tendency of HLA alleles to present peptides from proteins with certain functions may be among the factors contributing to differential association of alleles with infection outcomes. Comparison of HLA-B* 57:01 (associated with efficient control of viraemia and good disease prognosis in HIV-1 infection) and HLA-B* 08:01 (associated with poor prognosis) (8) yielded a set of GO categories that differ between genes enriched in ligands presented by each of the alleles (**Supplementary Figure S7**). Interestingly, when comparing GO terms of genes providing a source of ligands that are preferentially presented by either HLA-B* 57:01 or HLA-B* 08:01 we found that membrane and ion transport-related functions that are assigned to HIV genes are enriched in HLA-B* 57:01-presented proteins.

Finally, we hypothesized that the reported HLA presentation bias may be compensated for in haplotypes to increase the size of the immunopeptidome presented in each individual. To test this, we compared distributions of distances between GO enrichment profiles for pairs of HLA alleles associated with frequent haplotypes and for random pairs of alleles. On average, HLA-A/HLA-B and HLA-A/HLA-C pairs showed higher within-haplotype group distance compared to control (randomly-selected) allele pairs. The absence of the effect for HLA-B/HLA-C pairs may be explained by the fact that different HLA-B and HLA-C alleles are more similar to each other than either of them are to HLA-A in terms of amino acid sequence, a result of the evolutionary origin of the HLA-C gene being from duplication of HLA-B (37). These results may suggest that haplotypes of HLA alleles with different preferences for presenting proteins with particular molecular functions are evolutionarily beneficial and have a greater chance of becoming fixed in the population. It should be noted that in a 2013 paper Rao *et al*. (38) reported that complementarity of binding motifs in frequent HLA-A/HLA-B haplotypes is not higher than in random HLA-A/HLA-B pairs. The difference between earlier results and our conclusions may be explained by the lower accuracy of the older version of NetMHCpan software which was used in the Rao et al. study. NetMHCpan v2.0 was released in 2009 and was trained on the limited *in vitro* binding affinity data then available, but v4.0 used in this study was released in 2017 and was trained on a much larger dataset that additionally incorporated MS eluted ligand data.

Thorough investigation of HLA presentation biases can lead to better understanding of mechanisms underlying the existence of both beneficial HLA alleles and those alleles leading to disease susceptibility in various scenarios ranging from infectious diseases to autoimmunity. The COVID-19 pandemic has highlighted the necessity of a rapid selection of vaccine targets. HLA binding preferences should evidently be taken into account together with the population frequency of HLA alleles during vaccine development. We hope that our findings can help to explain why certain HLAs are more likely to present peptides from specific viral proteins compared to others. Those presentation biases may arise due to evolutionary fine-tuning of the HLA presentation machinery optimizing selection of non-self peptides, a subject for future studies.

## Acknowledgements

This work was supported by RFBR grant No. 19-015-00551 (IZ), Ministry of Science and Higher Education of the Russian Federation grant No. 075-15-2019-1789 (MS, DMC and DS), Medical Research Council (MRC) programme grant MR/K012037 (PB) and National Institutes of Health grant R01 AI 118549 (PB). PB is a Jenner Institute Investigator. HK is funded by MRC Human Immunology Unit core funding. We would like to thank Masafumi Takaguchi for providing the CD4.721.221 cell line, Vasily Tsvetkov and Ekaterina Putintseva for their assistance with running HLA binding prediction algorithms.

## Author contributions

Experimental data: WP, TP, AN, SB, NT and PB. Data analysis: VK, WP, IBW, MS. Manuscript draft and editing: MS, VK,IBW, IZ, DMC, HK, WP, PB. Supervised the study: MS, IZ, DMC, HK, PB.

## References

1. La Gruta, N.L., Gras, S., Daley, S.R., Thomas, P.G. and Rossjohn, J. (2018) Understanding the drivers of MHC restriction of T cell receptors. Nat. Rev. Immunol., 18, 467–478.

2. Trolle, T., McMurtrey, C.P., Sidney, J., Bardet, W., Osborn, S.C., Kaever, T., Sette, A., Hildebrand, W.H., Nielsen, M. and Peters, B. (2016) The length distribution of class I restricted T cell epitopes is determined by both peptide supply and MHC allele specific binding preference. J. Immunol. Baltim. Md 1950, 196, 1480–1487.

3. Sarkizova, S., Klaeger, S., Le, P.M., Li, L.W., Oliveira, G., Keshishian, H., Hartigan, C.R., Zhang, W., Braun, D.A., Ligon, K.L., et al. (2020) A large peptidome dataset improves HLA class I epitope prediction across most of the human population. Nat. Biotechnol., 38, 199–209.

4. Racle, J., Michaux, J., Rockinger, G.A., Arnaud, M., Bobisse, S., Chong, C., Guillaume, P., Coukos, G., Harari, A., Jandus, C., et al. (2019) Robust prediction of HLA class II epitopes by deep motif deconvolution of immunopeptidomes. Nat. Biotechnol., 37, 1283–1286.

5. Gfeller, D. and Bassani-Sternberg, M. (2018) Predicting Antigen Presentation—What Could We Learn From a Million Peptides? Front. Immunol., 9.

6. Robinson, J., Barker, D.J., Georgiou, X., Cooper, M.A., Flicek, P. and Marsh, S.G.E. (2020) IPD-IMGT/HLA Database. Nucleic Acids Res., 48, D948–D955.

7. Wang, S.-F., Chen, K.-H., Chen, M., Li, W.-Y., Chen, Y.-J., Tsao, C.-H., Yen, M., Huang, J.C. and Chen, Y.-M.A. (2011) Human-Leukocyte Antigen Class I Cw 1502 and Class II DR 0301 Genotypes Are Associated with Resistance to Severe Acute Respiratory Syndrome (SARS) Infection. Viral Immunol., 24, 421–426.

8. Košmrlj, A., Read, E.L., Qi, Y., Allen, T.M., Altfeld, M., Deeks, S.G., Pereyra, F., Carrington, M., Walker, B.D. and Chakraborty, A.K. (2010) Effects of thymic selection of the T cell repertoire on HLA-class I associated control of HIV infection. Nature, 465, 350–354.

9. Vita, R., Mahajan, S., Overton, J.A., Dhanda, S.K., Martini, S., Cantrell, J.R., Wheeler, D.K., Sette, A. and Peters, B. (2019) The Immune Epitope Database (IEDB): 2018 update. Nucleic Acids Res., 47, D339–D343.

10. Nielsen, M., Andreatta, M., Peters, B. and Buus, S. (2020) Immunoinformatics: Predicting Peptide–MHC Binding. Annu. Rev. Biomed. Data Sci., 3, 191–215.

11. Peters, B., Nielsen, M. and Sette, A. (2020) T Cell Epitope Predictions. Annu. Rev. Immunol., 38, 123–145.

12. Jurtz, V., Paul, S., Andreatta, M., Marcatili, P., Peters, B. and Nielsen, M. (2017) NetMHCpan-4.0: Improved Peptide-MHC Class I Interaction Predictions Integrating Eluted Ligand and Peptide Binding Affinity Data. J. Immunol. Baltim. Md 1950, 199, 3360–3368.

13. O’Donnell, T.J., Rubinsteyn, A. and Laserson, U. (2020) MHCflurry 2.0: Improved Pan-Allele Prediction of MHC Class I-Presented Peptides by Incorporating Antigen Processing. Cell Syst., 11, 42-48.e7.

14. Mei, S., Li, F., Leier, A., Marquez-Lago, T.T., Giam, K., Croft, N.P., Akutsu, T., Smith, A.I., Li, J., Rossjohn, J., et al. (2020) A comprehensive review and performance evaluation of bioinformatics tools for HLA class I peptide-binding prediction. Brief. Bioinform., 21, 1119–1135.

15. The HLA Ligand Atlas. A resource of natural HLA ligands presented on benign tissues | bioRxiv.

16. Perez, M.A.S., Bassani-Sternberg, M., Coukos, G., Gfeller, D. and Zoete, V. (2019) Analysis of Secondary Structure Biases in Naturally Presented HLA-I Ligands. Front. Immunol., 10.

17. Bassani-Sternberg, M., Pletscher-Frankild, S., Jensen, L.J. and Mann, M. (2015) Mass Spectrometry of Human Leukocyte Antigen Class I Peptidomes Reveals Strong Effects of Protein Abundance and Turnover on Antigen Presentation. Mol. Cell. Proteomics, 14, 658–673.

18. Pearson, H., Daouda, T., Granados, D.P., Durette, C., Bonneil, E., Courcelles, M., Rodenbrock, A., Laverdure, J.-P., Côté, C., Mader, S., et al. (2016) MHC class I– associated peptides derive from selective regions of the human genome. J. Clin. Invest., 126, 4690–4701.

19. Juncker, A.S., Larsen, M.V., Weinhold, N., Nielsen, M., Brunak, S. and Lund, O. (2009) Systematic Characterisation of Cellular Localisation and Expression Profiles of Proteins Containing MHC Ligands. PLOS ONE, 4, e7448.

20. Scull, K.E., Dudek, N.L., Corbett, A.J., Ramarathinam, S.H., Gorasia, D.G., Williamson, N.A. and Purcell, A.W. (2012) Secreted HLA recapitulates the immunopeptidome and allows in-depth coverage of HLA A* 02:01 ligands. Mol. Immunol., 51, 136–142.

21. Schellens, I.M.M., Hoof, I., Meiring, H.D., Spijkers, S.N.M., Poelen, M.C.M., Brink, J.A.M. van G. den, Poel, K. van der, Costa, A.I., Els, C.A.C.M. van, Baarle, D. van, et al. (2015) Comprehensive Analysis of the Naturally Processed Peptide Repertoire: Differences between HLA-A and B in the Immunopeptidome. PLOS ONE, 10, e0136417.

22. McMurtrey, C., Bardet, W., Osborn, S., Jackson, K., Schafer, F. and Hildebrand, W. (2015) Comparison of HLA-A and HLA-B ligandomes. Hum. Immunol., 76, 149.

23. Müller, M., Gfeller, D., Coukos, G. and Bassani-Sternberg, M. (2017) ‘Hotspots’ of Antigen Presentation Revealed by Human Leukocyte Antigen Ligandomics for Neoantigen Prioritization. Front. Immunol., 8.

24. Abelin, J.G., Keskin, D.B., Sarkizova, S., Hartigan, C.R., Zhang, W., Sidney, J., Stevens, J., Lane, W., Zhang, G.L., Eisenhaure, T.M., et al. (2017) Mass Spectrometry Profiling of HLA-Associated Peptidomes in Mono-allelic Cells Enables More Accurate Epitope Prediction. Immunity, 46, 315–326.

25. Croft, N.P., Smith, S.A., Pickering, J., Sidney, J., Peters, B., Faridi, P., Witney, M.J., Sebastian, P., Flesch, I.E.A., Heading, S.L., et al. (2019) Most viral peptides displayed by class I MHC on infected cells are immunogenic. Proc. Natl. Acad. Sci., 116, 3112–3117.

26. Prota, G., Gileadi, U., Rei, M., Lechuga-Vieco, A.V., Chen, J.-L., Galiani, S., Bedard, M., Lau, V.W.C., Fanchi, L.F., Artibani, M., et al. (2020) Enhanced Immunogenicity of Mitochondrial-Localized Proteins in Cancer Cells. Cancer Immunol. Res., 8, 685–697.

27. Paes, W., Leonov, G., Partridge, T., Chikata, T., Murakoshi, H., Frangou, A., Brackenridge, S., Nicastri, A., Smith, A.G., Learn, G.H., et al. (2019) Contribution of proteasome-catalyzed peptide cis-splicing to viral targeting by CD8+ T cells in HIV-1 infection. Proc. Natl. Acad. Sci., 116, 24748–24759.

28. Ritchie, M.E., Phipson, B., Wu, D., Hu, Y., Law, C.W., Shi, W. and Smyth, G.K. (2015) limma powers differential expression analyses for RNA-sequencing and microarray studies. Nucleic Acids Res., 43, e47–e47.

29. Huang, D.W., Sherman, B.T. and Lempicki, R.A. (2009) Systematic and integrative analysis of large gene lists using DAVID bioinformatics resources. Nat. Protoc., 4, 44–57.

30. Gonzalez-Galarza, F.F., McCabe, A., Santos, E.J.M. dos Jones, J., Takeshita, L., Ortega-Rivera, N.D., Cid-Pavon, G.M.D., Ramsbottom, K., Ghattaoraya, G., Alfirevic, A., et al. (2020) Allele frequency net database (AFND) 2020 update: gold-standard data classification, open access genotype data and new query tools. Nucleic Acids Res., 48, D783–D788.

31. Burroughs, N.J., de Boer, R.J. and Keşmir, C. (2004) Discriminating self from nonself with short peptides from large proteomes. Immunogenetics, 56, 311–320.

32. Structures of HLA-A* 1101 Complexed with Immunodominant Nonamer and Decamer HIV-1 Epitopes Clearly Reveal the Presence of a Middle, Secondary Anchor Residue | The Journal of Immunology.

33. DeWitt, W.S., Smith, A., Schoch, G., Hansen, J.A., Matsen, F.A. and Bradley, P. (2018) Human T cell receptor occurrence patterns encode immune history, genetic background, and receptor specificity. eLife, 7.

34. Pogorelyy, M.V., Fedorova, A.D., McLaren, J.E., Ladell, K., Bagaev, D.V., Eliseev, A.V., Mikelov, A.I., Koneva, A.E., Zvyagin, I.V., Price, D.A., et al. (2018) Exploring the pre-immune landscape of antigen-specific T cells. Genome Med., 10, 68.

35. Zvyagin, I.V., Tsvetkov, V.O., Chudakov, D.M. and Shugay, M. (2020) An overview of immunoinformatics approaches and databases linking T cell receptor repertoires to their antigen specificity. Immunogenetics, 72, 77–84.

36. Paul, S., Weiskopf, D., Angelo, M.A., Sidney, J., Peters, B. and Sette, A. (2013) HLA class I alleles are associated with peptide-binding repertoires of different size, affinity, and immunogenicity. J. Immunol. Baltim. Md 1950, 191, 5831–5839.

37. Robinson, J., Guethlein, L.A., Cereb, N., Yang, S.Y., Norman, P.J., Marsh, S.G.E. and Parham, P. (2017) Distinguishing functional polymorphism from random variation in the sequences of >10,000 HLA-A, -B and -C alleles. PLOS Genet., 13, e1006862.

38. Rao, X., De Boer, R.J., van Baarle, D., Maiers, M. and Kesmir, C. (2013) Complementarity of Binding Motifs is a General Property of HLA-A and HLA-B Molecules and Does Not Seem to Effect HLA Haplotype Composition. Front. Immunol., 4.

