## Supplementary Figures and Tables for "HLA binding of self-peptides is biased towards proteins with specific molecular functions"

**Supplementary Table S1**. Number of predicted 9-mer HLA ligands, HLA ligand enriched (HLEP) and depleted (HLDP) proteins for every HLA allele used in this study.

| Allele | Number of enriched genes | Number of depleted genes | Ligand count |
|---|---|---|---|
| HLA-A*02:01 | 1946 | 1557 | 655638 |
| HLA-A*11:01 | 55 | 106 | 498485 |
| HLA-B*07:02 | 614 | 220 | 365507 |
| HLA-B*27:05 | 671 | 239 | 333662 |
| HLA-C*02:02 | 1015 | 770 | 657146 |
| HLA-C*15:02 | 964 | 500 | 553180 |

**Supplementary Table S2.** Number of predicted HLA-A*11:01 ligands, HLA ligand enriched (HLEP) and depleted (HLDP) proteins for 8-12mer peptides.

| Ligand length | Number of enriched genes | Number of depleted genes | Ligand count |
|---|---|---|---|
| 8 | 56 | 25 | 89334 |
| 9 | 55 | 106 | 498485 |
| 10 | 108 | 233 | 676915 |
| 11 | 306 | 295 | 354065 |
| 12 | 362 | 443 | 422447 |

**Supplementary Table S3. Extended set of 93 HLA alleles used for validation of the conclusions from a smaller set.** The list and frequencies of the alleles covering 95% of individuals worldwide were taken from Sarkizova et al. [https://www.nature.com/articles/s41587-019-0322-9#Sec33]. Anchor residues in the second (P2) and the ninth (P9) positions were inferred if the aggregated fraction of one of the groups of amino acids with similar physicochemical properties was greater than 0.5 for the corresponding position of netMHCpan binders for that allele.

| Allele | Freq. | P2 anchor | P9 anchor |
|---|---|---|---|
| HLA-A0101 | 0.048 | ST | |
| HLA-A0201 | 0.153 | LFIMV | LFIMV |
| HLA-A0202 | 0.008 | LFIMV | LFIMV |
| HLA-A0203 | 0.015 | LFIMV | LFIMV |
| HLA-A0204 | 0.003 | LFIMV | LFIMV |
| HLA-A0205 | 0.008 | LFIMV | LFIMV |
| HLA-A0206 | 0.035 | LFIMV | LFIMV |
| HLA-A0207 | 0.021 | LFIMV | LFIMV |
| HLA-A0211 | 0.003 | LFIMV | LFIMV |
| HLA-A0301 | 0.043 | LFIMV | RK |
| HLA-A1101 | 0.117 | | RK |
| HLA-A1102 | 0.007 | | RK |
| HLA-A2301 | 0.023 | | LFIMV |
| HLA-A2402 | 0.188 | | LFIMV |
| HLA-A2407 | 0.005 | | LFIMV |
| HLA-A2501 | 0.005 | | LFIMV |
| HLA-A2601 | 0.034 | | LFIMV |
| HLA-A2902 | 0.016 | | Y |
| HLA-A3001 | 0.025 | | |
| HLA-A3002 | 0.015 | | Y |
| HLA-A3101 | 0.041 | | RK |
| HLA-A3201 | 0.014 | LFIMV | LFIMV |
| HLA-A3301 | 0.012 | | RK |
| HLA-A3303 | 0.041 | | RK |
| HLA-A3401 | 0.016 | | RK |
| HLA-A3402 | 0.005 | | RK |
| HLA-A3601 | 0.004 | | Y |
| HLA-A6601 | 0.006 | | |
| HLA-A6801 | 0.023 | | RK |
| HLA-A6802 | 0.013 | | LFIMV |
| HLA-A7401 | 0.008 | LFIMV | RK |

| Allele | Freq. | P2 anchor | P9 anchor |
|---|---|---|---|
| HLA-B0702 | 0.041 | P | LFIMV |
| HLA-B0704 | 0.031 | P | LFIMV |
| HLA-B0801 | 0.03 | | LFIMV |
| HLA-B1301 | 0.026 | | LFIMV |
| HLA-B1302 | 0.014 | | LFIMV |
| HLA-B1402 | 0.012 | | LFIMV |
| HLA-B1501 | 0.034 | | LFIMV |
| HLA-B1502 | 0.013 | | LFIMV |
| HLA-B1503 | 0.012 | | LFIMV |
| HLA-B1510 | 0.006 | | LFIMV |
| HLA-B1517 | 0.003 | ST | LFIMV |
| HLA-B1801 | 0.023 | DE | |
| HLA-B2705 | 0.012 | RK | |
| HLA-B3501 | 0.055 | | LFIMV |
| HLA-B3503 | 0.009 | | LFIMV |
| HLA-B3507 | 0.013 | | |
| HLA-B3701 | 0.025 | DE | LFIMV |
| HLA-B3801 | 0.008 | | LFIMV |
| HLA-B3802 | 0.009 | | LFIMV |
| HLA-B4001 | 0.051 | DE | LFIMV |
| HLA-B4002 | 0.042 | DE | LFIMV |
| HLA-B4006 | 0.018 | DE | LFIMV |
| HLA-B4201 | 0.01 | P | LFIMV |
| HLA-B4402 | 0.022 | DE | LFIMV |
| HLA-B4403 | 0.045 | DE | LFIMV |
| HLA-B4501 | 0.01 | DE | |
| HLA-B4601 | 0.024 | | LFIMV |
| HLA-B4901 | 0.009 | DE | LFIMV |
| HLA-B5001 | 0.009 | DE | |
| HLA-B5101 | 0.052 | | LFIMV |
| HLA-B5201 | 0.023 | | LFIMV |
| HLA-B5301 | 0.016 | | LFIMV |
| HLA-B5401 | 0.014 | P | |
| HLA-B5501 | 0.006 | P | |
| HLA-B5502 | 0.01 | P | |
| HLA-B5601 | 0.014 | P | |
| HLA-B5701 | 0.01 | ST | LFIMV |
| HLA-B5703 | 0.005 | ST | LFIMV |
| HLA-B5801 | 0.029 | ST | LFIMV |
| HLA-B5802 | 0.008 | ST | LFIMV |

| Allele | Freq. | P2 anchor | P9 anchor |
|---|---|---|---|
| HLA-C0102 | 0.085 | | LFIMV |
| HLA-C0202 | 0.028 | | LFIMV |
| HLA-C0302 | 0.025 | | LFIMV |
| HLA-C0303 | 0.056 | | LFIMV |
| HLA-C0304 | 0.091 | | LFIMV |
| HLA-C0401 | 0.112 | | LFIMV |
| HLA-C0403 | 0.019 | | LFIMV |
| HLA-C0501 | 0.026 | | LFIMV |
| HLA-C0602 | 0.062 | | LFIMV |
| HLA-C0701 | 0.069 | | LFIMV |
| HLA-C0702 | 0.131 | | LFIMV |
| HLA-C0704 | 0.015 | | LFIMV |
| HLA-C0801 | 0.045 | | LFIMV |
| HLA-C0802 | 0.02 | | LFIMV |
| HLA-C1202 | 0.032 | | LFIMV |
| HLA-C1203 | 0.02 | | LFIMV |
| HLA-C1402 | 0.025 | | LFIMV |
| HLA-C1403 | 0.015 | | LFIMV |
| HLA-C1502 | 0.034 | | LFIMV |
| HLA-C1601 | 0.024 | | LFIMV |
| HLA-C1701 | 0.019 | | LFIMV |
| HLA-C1801 | 0.006 | | LFIMV |

**Supplementary Table S4. Populations included in the analysis of HLA presentation bias compensation in haplotypes.** Haplotypes and their frequencies were taken from http://allelefrequencies.net/.

| Population | Sample size | Number of haplotypes |
|---|---|---|
| USA NMDP Caribbean Indian | 14,339 | 89 |
| USA NMDP American Indian South or Central America | 5,926 | 65 |
| USA NMDP Hawaiian or other Pacific Islander | 11,499 | 63 |
| USA NMDP Filipino | 50,614 | 51 |
| USA NMDP Hispanic South or Central American | 146,714 | 45 |
| USA NMDP Chinese | 99,672 | 44 |
| USA NMDP Caribbean Hispanic | 115,374 | 42 |
| USA NMDP Mexican or Chicano | 261,235 | 42 |
| USA NMDP Southeast Asian | 27,978 | 39 |
| USA NMDP Japanese | 24,582 | 38 |
| USA NMDP Korean | 77,584 | 37 |
| USA NMDP North American Amerindian | 35,791 | 35 |
| USA NMDP Caribbean Black | 33,328 | 34 |
| USA NMDP Vietnamese | 43,540 | 34 |
| USA NMDP Middle Eastern or North Coast of Africa | 70,890 | 33 |
| USA NMDP African American pop 2 | 416,581 | 29 |
| USA NMDP European Caucasian | 1,242,890 | 29 |
| USA NMDP South Asian Indian | 185,391 | 29 |
| USA NMDP African | 28,557 | 28 |

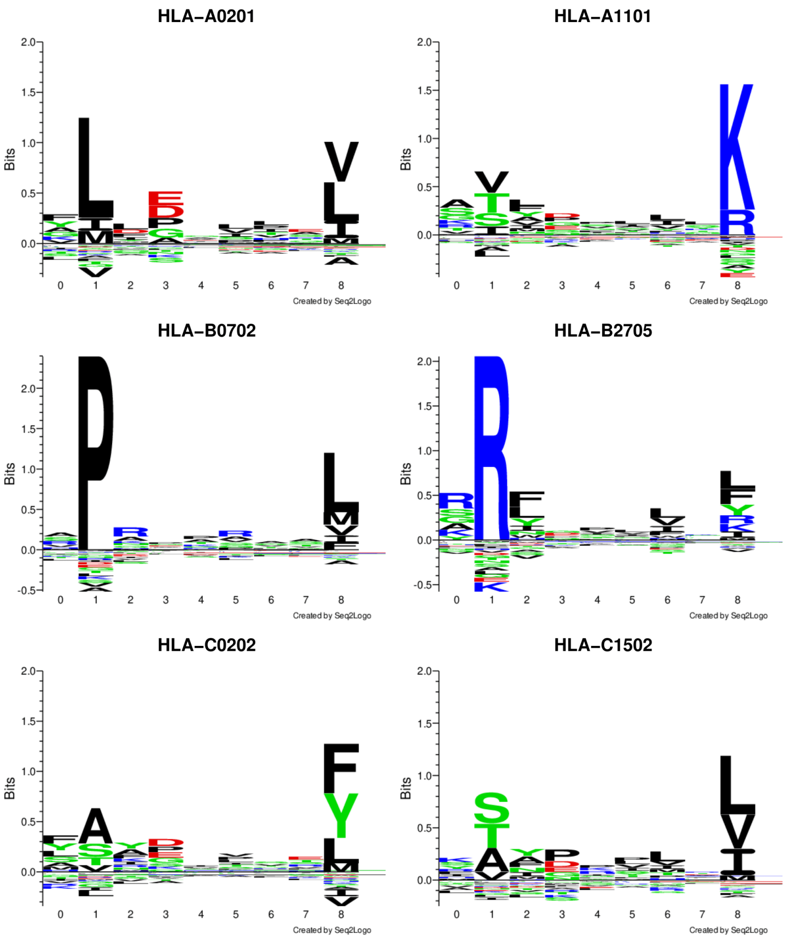


**Supplementary Figure S1. Binding motifs for surveyed HLA alleles**. Motifs for naturally presented HLA ligands for a sample set of alleles used in the study (data from http://www.cbs.dtu.dk/services/NetMHCpan/logos_ps.php).

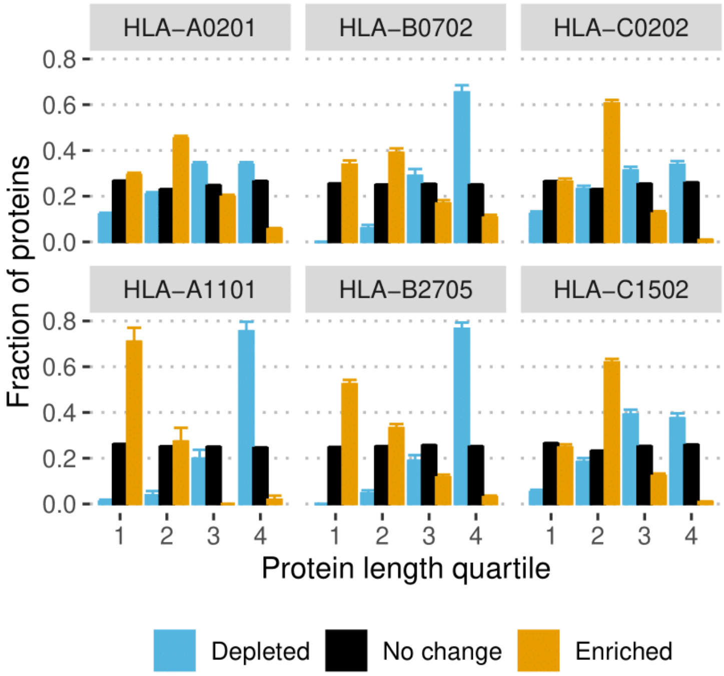


**Supplementary Figure S2. Length bias for proteins enriched (HLEPs) and depleted (HLDPs) in HLA ligands.** Plot shows the fraction of genes expressing proteins residing in four gene quartiles (from 1 - shortest to 4 - longest) in HLEP (Enriched) and HLDP (Depleted) sets compared to remaining proteins (No change). Error bars show 95% confidence interval for fractions. Note that genes enriched in HLA ligands are shifted towards shorter length while depleted ones are shifted towards longer lengths.

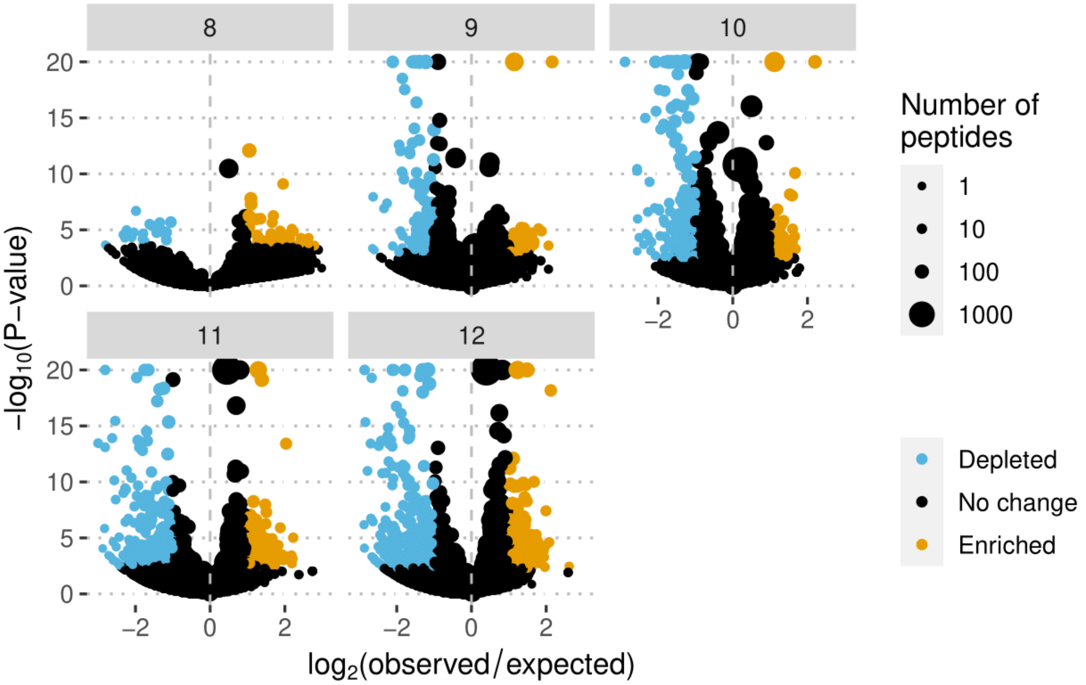


**Supplementary Figure S3. Ligand enrichment analysis for HLA-A*11:01 allele and different ligand lengths.** Volcano plots showing the log of the ratio of observed and expected HLA-A*11:01 ligands for each human gene plotted against enrichment P-value computed using binomial test. Point size shows number of predicted HLA ligands, point color highlights genes enriched and depleted in ligands according to 2 times odd differences and adjusted P-value of < 0.05. Data for different peptide lengths (8, 9, 10, 11 and 12 amino acids) are shown as separate plots.

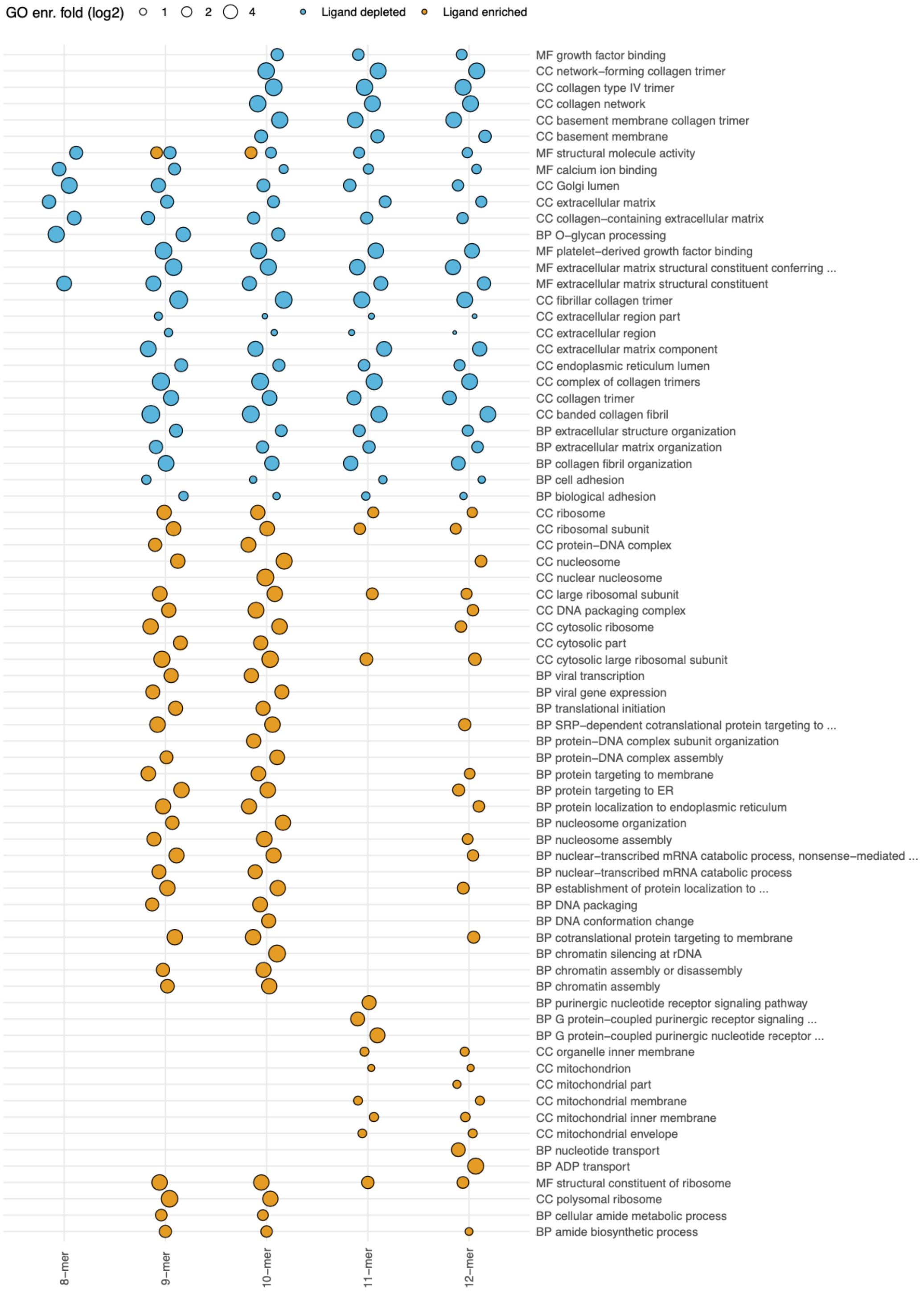


**Supplementary Figure S4. Gene ontology (GO) term enrichment analysis for human genes differentially enriched in ligands of various lengths by HLA-A*11:01.** Point size represents the GO enrichment fold for genes enriched (yellow) and depleted (blue) in HLA 8- to 12-mer HLA ligands. An adjusted P-value threshold of 0.01 was chosen and top 20 GO categories were chosen for each HLA allele.

**A**

| Ontology | Term | Alleles |
|---|---|---|
| MF | structural molecule activity | 89 (96%) |
| CC | supramolecular complex | 86 (92%) |
| CC | supramolecular polymer | 86 (92%) |
| MF | extracellular matrix structural constituent | 84 (90%) |
| CC | collagen trimer | 84 (90%) |
| CC | fibrillar collagen trimer | 84 (90%) |
| MF | extracellular matrix structural constituent conferring tensile strength | 84 (90%) |
| CC | banded collagen fibril | 84 (90%) |
| CC | complex of collagen trimers | 83 (89%) |
| MF | platelet–derived growth factor binding | 82 (88%) |
| CC | collagen type IV trimer | 81 (87%) |
| BP | extracellular matrix organization | 81 (87%) |
| BP | extracellular structure organization | 81 (87%) |
| CC | network–forming collagen trimer | 81 (87%) |
| CC | collagen network | 81 (87%) |
| CC | basement membrane collagen trimer | 81 (87%) |
| CC | extracellular matrix | 80 (86%) |
| BP | collagen–activated tyrosine kinase receptor signaling pathway | 80 (86%) |
| CC | collagen–containing extracellular matrix | 80 (86%) |
| CC | endoplasmic reticulum lumen | 76 (82%) |

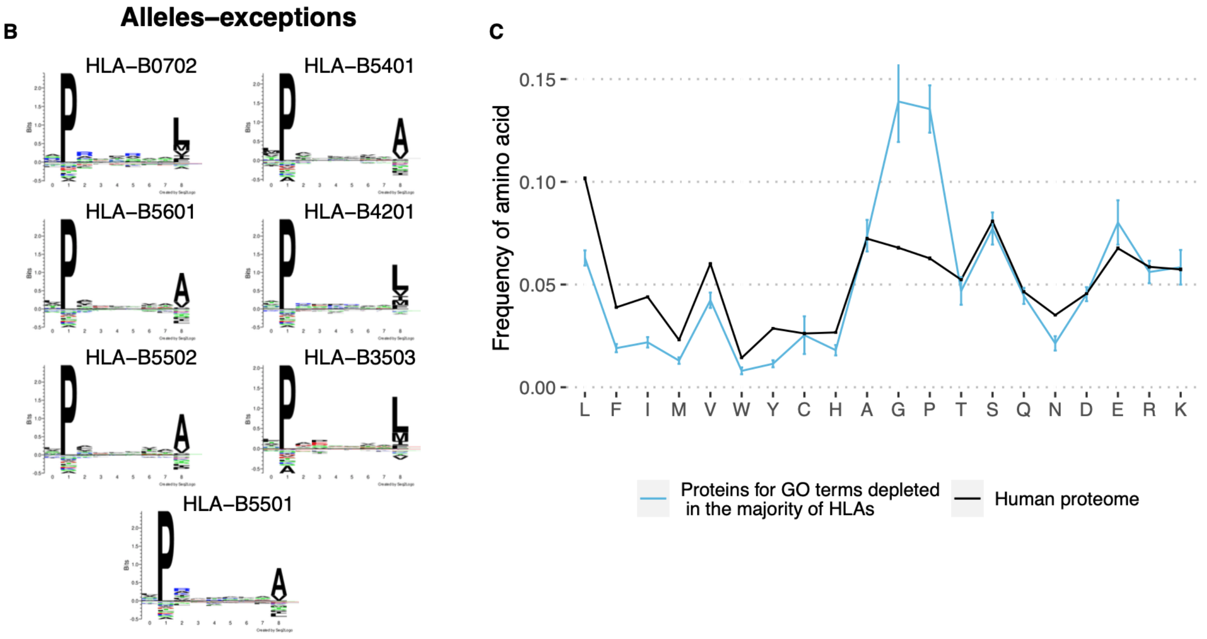


**Supplementary Figure S5. GO categories associated with HLDPs for most alleles in the extended dataset. A.** GO terms corresponding to HLDPs for more than 80% of the surveyed 93 HLA alleles. **B.** Motifs of naturally presented ligands for alleles-exceptions in which less than 15% of GO terms from table **A.** are depleted. Note that all these alleles require proline as anchor residue in P2. For all the alleles without proline anchors at least 70% of GO terms from table **A.** are depleted. **C.** Comparison of amino acid composition of proteins corresponding to GO categories from **A.** (consider only proteins which are in HLDPs for at least half of surveyed 93 alleles but not for any of the list from **B.**) and human proteome. Error bars show 95% confidence interval for the mean value. Note that glycines and prolines are enriched in these proteins as in **Figure 3C**.

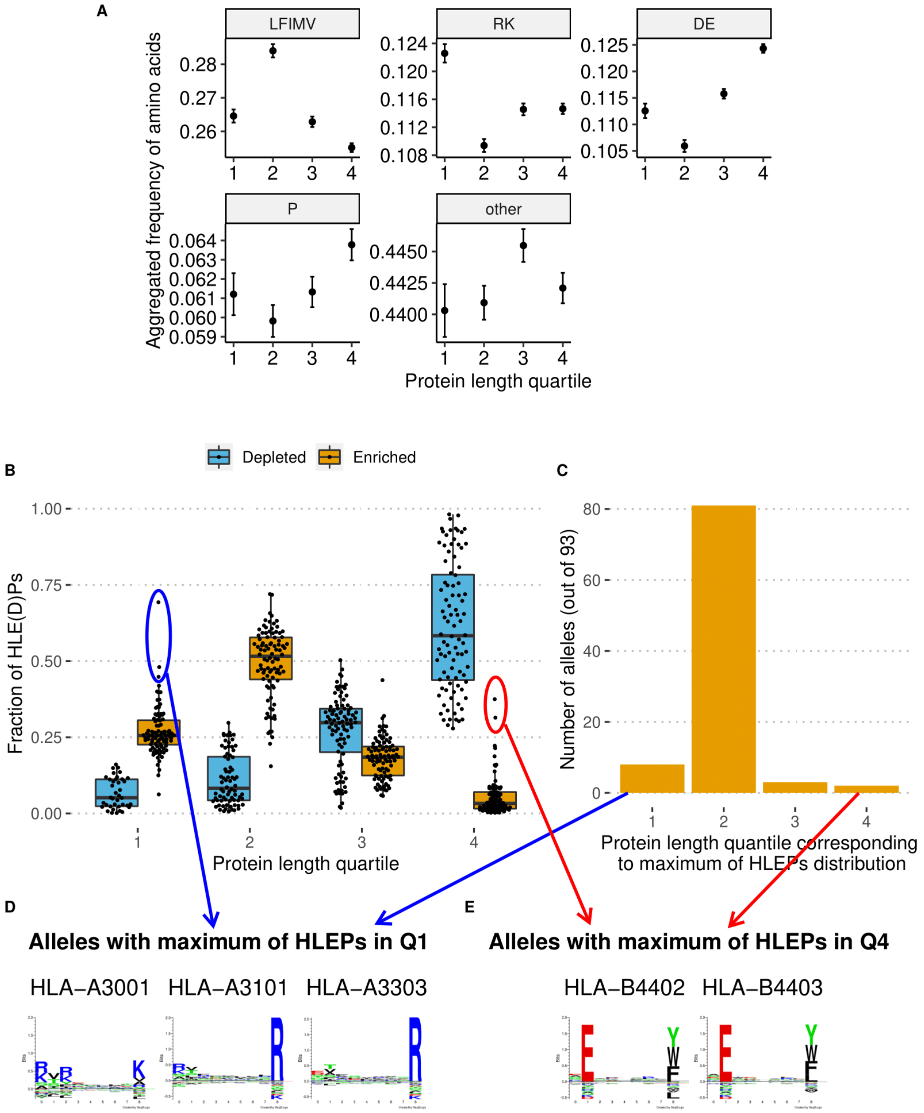


**Supplementary Figure S6. Analysis of length bias of HLE(D)Ps on an extended dataset from 93 HLA alleles. A.** Fractions of amino acids in proteins of different length quartiles. Amino acids are grouped as the following: "LFIMV" stands for hydrophobic amino acids (leucine, phenylalanine, isoleucine, methionine, and valine), "DE" - negatively charged amino acids (aspartic acid and glutamic acid), "RK" - positively charged amino acids (arginine and lysine), P - proline, "other" - the remaining 10 amino acids which are rarely used as anchor residues. Note that the highest fractions of hydrophobic, positively and negatively charged amino acids are for proteins from Q2, Q1, and Q4, respectively. Error bars show 95% confidence interval for the mean value. **B.** Distribution of the length (split to quartiles) of HLE(D)Ps for the surveyed 93 HLA alleles. Note that on average this distribution peaks in Q2 for HLEPs and in Q4 for HLDPs. **C.** Number of alleles for which HLEPs length distributions profiles from **B.** have maximums in Q1, Q2, Q3, and Q4. Note that for the majority of alleles (80 out of 93) the highest fraction of HLEPs correspond to Q2 proteins, while some alleles have preferences for presentation of proteins from Q1, Q3, or Q4. **D-E**. Motifs of naturally presented ligands for alleles, which have distributions of HLEPs with the maximum on Q1 (**D**) or Q4 (**E**). Note that "Q1-max" alleles require positively charged anchor residues which are enriched in Q1 proteins (see **A.**) while "Q4-max" ones require negatively charged anchor residues which are enriched in Q4 proteins. Motifs are taken from http://www.cbs.dtu.dk/services/NetMHCpan/logos_ps.php.

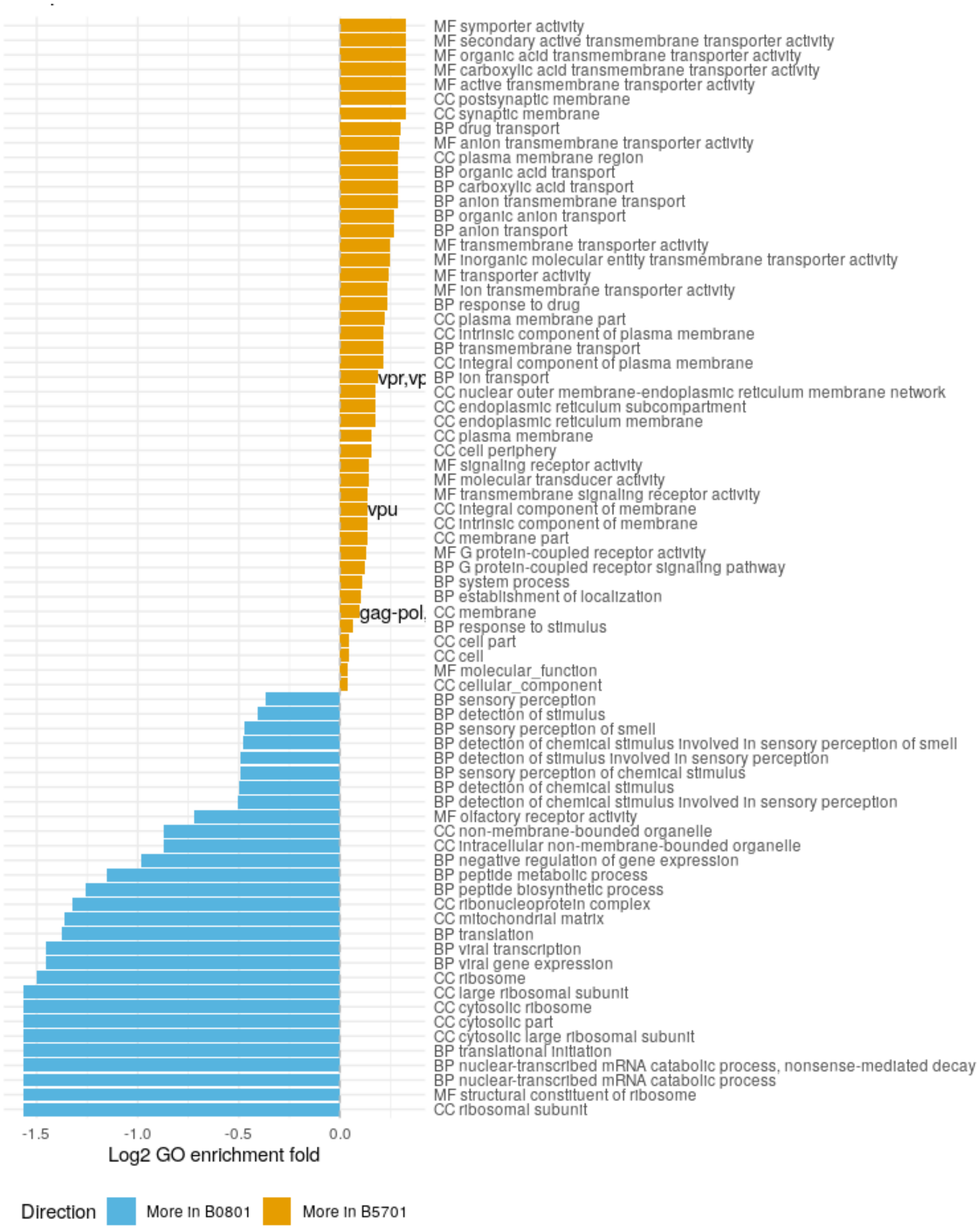


**Supplementary Figure S7. Analysis of GO categories enriched for HIV-protective and non-protective HLA alleles.** Bar plot shows GO category enrichment fold for HLA-B*57:01 (HIV-protective, yellow) and HLA-B*08:01 (non-protective, blue) HLA alleles. Only categories with an adjusted enrichment P-value less than 0.01 are shown, P-values were computed in a GO enrichment test where ligand-enriched genes for HLA-B*57:01 were used as a test set and HLA-B*08:01-ligand-enriched genes were used as control and vise versa. HIV protein names corresponding to GO categories are shown according to data from UniProt/QuickGO.
