## Supplementary Note 1 for "HLA binding of self-peptides is biased towards proteins with specific molecular functions"

#### Supplementary Note 1. Independent validation of HLA presentation biases using MHCflurry and DAVID software tools

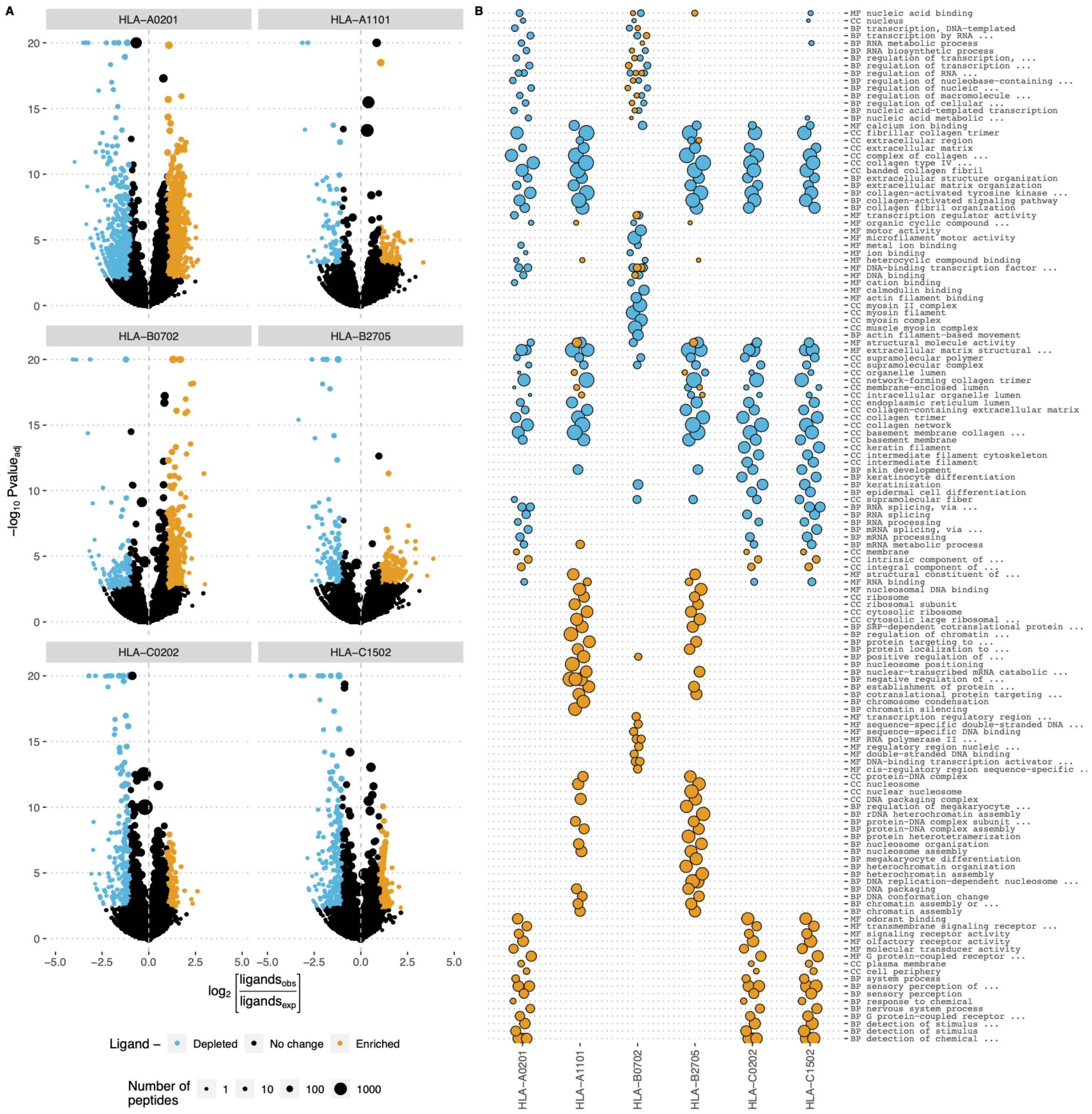


**Figure SN1. Human genes enriched and depleted in HLA ligands (HLEPs and HLDPs, respectively) and their associated Gene Ontology (GO) categories according to MHCflurry predictions.** Legend is the same as in Figure 2 of the main text. The analysis was based on the "Presentation score" of MHCflurry instead of MHC binding prediction ("Rank") of NetMHCpan. Using affinity predictions of MHCflurry ("Affinity percentile") resulted in qualitatively the same results (data not shown).

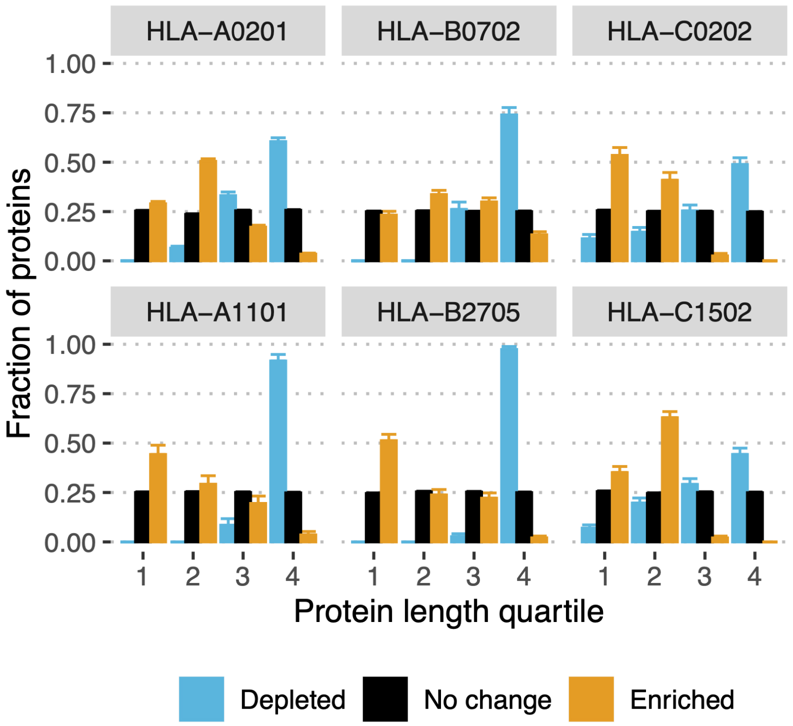


**Figure SN2. Length bias for proteins enriched and depleted in HLA ligands according to MHCflurry predictions.** Legend is the same as in Fig. S1 of the main text. The analysis was based on the “Presentation score” of MHCflurry instead of MHC binding prediction (“Rank”) of netMHCpan. Using affinity predictions of MHCflurry (“Affinity percentile”) resulted in qualitatively the same results (data not shown).

**Figure SN3. GO term enrichment analysis using DAVID web tool.** Analysis was performed for sets of HLEPs and HLDPs (denoted as "_enr" and "_depl" suffixes in the "Current gene list" section, respectively) for the 6 surveyed alleles as in Figure 2B and Figure SN1B but using DAVID web tool (https://david.ncifcrf.gov/). This analysis groups GO categories in annotation clusters describing similar sets of gene functions. Several clusters with the highest enrichment scores are presented.

#### Functional Annotation Clustering

Help and Manual

**Current Gene List: HLA-A0201_enr**
**Current Background: Homo sapiens**
**1946 DAVID IDs**

⊞ Options Classification Stringency Medium

Rerun using options | Create Sublist

**159 Cluster(s)** Download File

| Annotation Cluster 1 | Enrichment Score: ? | | Count | P_Value | Benjamini |
|---|---|---|---|---|---|
| UP_KEYWORDS | Transmembrane | RT | 1819 | 0.0E0 | 0.0E0 |
| UP_KEYWORDS | Transmembrane helix | RT | 1818 | 0.0E0 | 0.0E0 |
| UP_KEYWORDS | Membrane | RT | 1834 | 0.0E0 | 0.0E0 |
| UP_SEQ_FEATURE | transmembrane region | RT | 1775 | 0.0E0 | 0.0E0 |
| GOTERM_CC_DIRECT | integral component of membrane | RT | 1656 | 0.0E0 | 0.0E0 |
| **Annotation Cluster 2** | **Enrichment Score: ?** | | **Count** | **P_Value** | **Benjamini** |
| UP_KEYWORDS | G-protein coupled receptor | RT | 683 | 0.0E0 | 0.0E0 |
| GOTERM_MF_DIRECT | olfactory receptor activity | RT | 380 | 0.0E0 | 0.0E0 |
| UP_KEYWORDS | Receptor | RT | 743 | 0.0E0 | 0.0E0 |
| UP_KEYWORDS | Transducer | RT | 683 | 0.0E0 | 0.0E0 |
| INTERPRO | Olfactory receptor | RT | 379 | 0.0E0 | 0.0E0 |
| INTERPRO | G protein-coupled receptor, rhodopsin-like | RT | 626 | 0.0E0 | 0.0E0 |
| UP_KEYWORDS | Olfaction | RT | 381 | 0.0E0 | 0.0E0 |
| GOTERM_BP_DIRECT | G-protein coupled receptor signaling pathway | RT | 597 | 0.0E0 | 0.0E0 |
| GOTERM_BP_DIRECT | detection of chemical stimulus involved in sensory perception of smell | RT | 379 | 0.0E0 | 0.0E0 |
| INTERPRO | GPCR, rhodopsin-like, 7TM | RT | 639 | 0.0E0 | 0.0E0 |
| UP_SEQ_FEATURE | topological domain:Extracellular | RT | 1059 | 0.0E0 | 0.0E0 |
| UP_SEQ_FEATURE | topological domain:Cytoplasmic | RT | 1181 | 0.0E0 | 0.0E0 |
| GOTERM_MF_DIRECT | G-protein coupled receptor activity | RT | 571 | 0.0E0 | 0.0E0 |
| UP_KEYWORDS | Cell membrane | RT | 974 | 4.5E-321 | 8.1E-319 |
| KEGG_PATHWAY | Olfactory transduction | RT | 368 | 5.4E-300 | 1.0E-297 |
| UP_KEYWORDS | Sensory transduction | RT | 424 | 6.2E-299 | 7.4E-297 |
| UP_SEQ_FEATURE | glycosylation site:N-linked (GlcNAc...) | RT | 1074 | 6.8E-281 | 8.1E-278 |
| UP_KEYWORDS | Glycoprotein | RT | 1109 | 7.9E-275 | 7.0E-273 |
| GOTERM_CC_DIRECT | plasma membrane | RT | 1081 | 3.1E-251 | 6.0E-249 |
| UP_SEQ_FEATURE | disulfide bond | RT | 628 | 4.1E-103 | 3.3E-100 |
| UP_KEYWORDS | Disulfide bond | RT | 692 | 2.4E-101 | 1.5E-99 |
| **Annotation Cluster 3** | **Enrichment Score: 40.79** | | **Count** | **P_Value** | **Benjamini** |
| UP_KEYWORDS | Symport | RT | 94 | 7.8E-76 | 4.0E-74 |
| UP_KEYWORDS | Sodium | RT | 67 | 3.8E-35 | 1.5E-33 |
| UP_KEYWORDS | Sodium transport | RT | 63 | 5.9E-33 | 2.1E-31 |
| GOTERM_BP_DIRECT | sodium ion transport | RT | 44 | 4.0E-22 | 1.1E-19 |

#### Functional Annotation Clustering

Help and Manual

**Current Gene List: HLA-A0201_depl**
**Current Background: Homo sapiens**
**1556 DAVID IDs**

Options Classification Stringency Medium

Rerun using options | Create Sublist

**129 Cluster(s)**

Download File

| Annotation Cluster 1 | | Enrichment Score: 153.09 | | | Count | P_Value | Benjamini |
|---|---|---|---|---|---|---|---|
| ☐ | INTERPRO | Zinc finger C2H2-type/integrase DNA-binding domain | RT | | 375 | 6.3E-237 | 6.3E-234 |
| ☐ | INTERPRO | Zinc finger, C2H2-like | RT | | 385 | 6.8E-235 | 3.4E-232 |
| ☐ | INTERPRO | Zinc finger, C2H2 | RT | | 393 | 1.8E-234 | 6.0E-232 |
| ☐ | UP_SEQ_FEATURE | zinc finger region:C2H2-type 5 | RT | | 323 | 3.6E-223 | 1.3E-219 |
| ☐ | UP_SEQ_FEATURE | zinc finger region:C2H2-type 6 | RT | | 308 | 7.8E-221 | 1.4E-217 |
| ☐ | UP_SEQ_FEATURE | zinc finger region:C2H2-type 4 | RT | | 329 | 4.0E-218 | 4.8E-215 |
| ☐ | UP_SEQ_FEATURE | zinc finger region:C2H2-type 7 | RT | | 287 | 1.9E-206 | 1.7E-203 |
| ☐ | UP_SEQ_FEATURE | zinc finger region:C2H2-type 8 | RT | | 276 | 1.7E-205 | 1.2E-202 |
| ☐ | UP_SEQ_FEATURE | zinc finger region:C2H2-type 3 | RT | | 329 | 2.0E-203 | 1.2E-200 |
| ☐ | SMART | ZnF_C2H2 | RT | | 385 | 1.1E-196 | 2.7E-194 |
| ☐ | UP_SEQ_FEATURE | zinc finger region:C2H2-type 9 | RT | | 257 | 3.8E-196 | 2.0E-193 |
| ☐ | UP_KEYWORDS | Zinc-finger | RT | | 528 | 1.8E-194 | 6.3E-192 |
| ☐ | UP_SEQ_FEATURE | zinc finger region:C2H2-type 10 | RT | | 240 | 2.6E-191 | 1.2E-188 |
| ☐ | UP_SEQ_FEATURE | zinc finger region:C2H2-type 2 | RT | | 310 | 1.7E-186 | 6.8E-184 |
| ☐ | UP_SEQ_FEATURE | zinc finger region:C2H2-type 11 | RT | | 220 | 1.0E-180 | 3.7E-178 |
| ☐ | UP_KEYWORDS | Nucleus | RT | | 899 | 4.5E-175 | 7.8E-173 |
| ☐ | UP_KEYWORDS | Transcription regulation | RT | | 580 | 1.3E-174 | 1.6E-172 |
| ☐ | UP_KEYWORDS | Transcription | RT | | 584 | 3.9E-171 | 3.4E-169 |
| ☐ | GOTERM_MF_DIRECT | nucleic acid binding | RT | | 378 | 8.3E-171 | 4.8E-168 |
| ☐ | INTERPRO | Krueppel-associated box | RT | | 242 | 2.7E-167 | 6.8E-165 |
| ☐ | GOTERM_BP_DIRECT | regulation of transcription, DNA-templated | RT | | 453 | 6.1E-166 | 1.7E-162 |
| ☐ | GOTERM_BP_DIRECT | transcription, DNA-templated | RT | | 515 | 6.2E-164 | 8.5E-161 |
| ☐ | UP_SEQ_FEATURE | zinc finger region:C2H2-type 12 | RT | | 192 | 1.2E-160 | 3.8E-158 |
| ☐ | UP_KEYWORDS | DNA-binding | RT | | 520 | 2.6E-157 | 1.8E-155 |
| ☐ | UP_KEYWORDS | Zinc | RT | | 557 | 7.4E-156 | 4.3E-154 |
| ☐ | UP_SEQ_FEATURE | domain:KRAB | RT | | 211 | 9.9E-149 | 3.0E-146 |
| ☐ | UP_SEQ_FEATURE | zinc finger region:C2H2-type 1 | RT | | 259 | 7.5E-144 | 2.1E-141 |
| ☐ | SMART | KRAB | RT | | 237 | 2.8E-138 | 3.5E-136 |

| Category | Term | RT | Count | P_Value | Benjamini |
|---|---|---|---|---|---|
| GOTERM_CC_DIRECT | nucleus | RT | 840 | 1.8E-135 | 9.3E-133 |
| UP_SEQ_FEATURE | zinc finger region:C2H2-type 13 | RT | 156 | 4.3E-132 | 1.1E-129 |
| UP_SEQ_FEATURE | zinc finger region:C2H2-type 14 | RT | 126 | 1.2E-111 | 3.0E-109 |
| GOTERM_MF_DIRECT | DNA binding | RT | 408 | 8.8E-107 | 2.6E-104 |
| GOTERM_MF_DIRECT | metal ion binding | RT | 453 | 1.7E-102 | 3.3E-100 |
| UP_SEQ_FEATURE | zinc finger region:C2H2-type 15 | RT | 107 | 3.6E-94 | 8.2E-92 |
| UP_KEYWORDS | Metal-binding | RT | 586 | 4.9E-85 | 2.5E-83 |
| GOTERM_MF_DIRECT | transcription factor activity, sequence-specific DNA binding | RT | 265 | 2.2E-78 | 3.3E-76 |
| UP_SEQ_FEATURE | zinc finger region:C2H2-type 16 | RT | 82 | 3.5E-71 | 7.6E-69 |
| UP_SEQ_FEATURE | zinc finger region:C2H2-type 17 | RT | 64 | 2.2E-54 | 4.2E-52 |
| UP_SEQ_FEATURE | zinc finger region:C2H2-type 18 | RT | 51 | 2.0E-43 | 3.4E-41 |
| GOTERM_CC_DIRECT | intracellular | RT | 235 | 3.9E-36 | 1.0E-33 |
| UP_SEQ_FEATURE | zinc finger region:C2H2-type 19 | RT | 40 | 2.5E-32 | 4.2E-30 |
| **Annotation Cluster 2** | **Enrichment Score: 18.29** | | **Count** | **P_Value** | **Benjamini** |
| UP_KEYWORDS | mRNA processing | RT | 92 | 2.5E-28 | 7.4E-27 |
| UP_KEYWORDS | mRNA splicing | RT | 80 | 5.5E-28 | 1.5E-26 |
| KEGG_PATHWAY | Spliceosome | RT | 36 | 1.6E-21 | 2.5E-19 |
| GOTERM_BP_DIRECT | RNA splicing | RT | 55 | 1.4E-20 | 1.3E-17 |
| GOTERM_BP_DIRECT | mRNA processing | RT | 57 | 2.2E-20 | 1.5E-17 |
| GOTERM_BP_DIRECT | mRNA splicing, via spliceosome | RT | 57 | 1.4E-15 | 7.3E-13 |
| UP_KEYWORDS | Spliceosome | RT | 32 | 3.7E-9 | 6.9E-8 |
| GOTERM_CC_DIRECT | catalytic step 2 spliceosome | RT | 26 | 1.3E-8 | 1.2E-6 |
| **Annotation Cluster 3** | **Enrichment Score: 16.39** | | **Count** | **P_Value** | **Benjamini** |
| INTERPRO | Nucleotide-binding, alpha-beta plait | RT | 84 | 2.8E-30 | 5.7E-28 |
| INTERPRO | RNA recognition motif domain | RT | 77 | 5.6E-30 | 9.3E-28 |
| SMART | RRM | RT | 75 | 1.0E-22 | 8.4E-21 |
| GOTERM_MF_DIRECT | nucleotide binding | RT | 87 | 6.5E-22 | 6.4E-20 |
| UP_SEQ_FEATURE | domain:RRM | RT | 49 | 7.7E-21 | 9.7E-19 |
| UP_KEYWORDS | RNA-binding | RT | 122 | 3.4E-20 | 7.9E-19 |
| GOTERM_MF_DIRECT | RNA binding | RT | 88 | 3.9E-10 | 2.6E-8 |
| UP_SEQ_FEATURE | domain:RRM 1 | RT | 27 | 1.7E-6 | 7.1E-5 |
| UP_SEQ_FEATURE | domain:RRM 2 | RT | 27 | 1.7E-6 | 7.1E-5 |
| UP_SEQ_FEATURE | domain:RRM 3 | RT | 10 | 3.8E-2 | 5.5E-1 |
| **Annotation Cluster 4** | **Enrichment Score: 14.94** | | **Count** | **P_Value** | **Benjamini** |
| INTERPRO | Transcription regulator SCAN | RT | 30 | 3.1E-17 | 4.4E-15 |
| UP_SEQ_FEATURE | domain:SCAN box | RT | 29 | 2.4E-16 | 2.1E-14 |
| INTERPRO | Retrovirus capsid, C-terminal | RT | 30 | 5.6E-15 | 7.1E-13 |
| SMART | SCAN | RT | 30 | 4.1E-14 | 2.6E-12 |
| **Annotation Cluster 5** | **Enrichment Score: 12.7** | | **Count** | **P_Value** | **Benjamini** |
| UP_SEQ_FEATURE | repeat:7 | RT | 49 | 3.7E-21 | 4.8E-19 |
| UP_SEQ_FEATURE | repeat:3 | RT | 64 | 1.2E-20 | 1.4E-18 |
| UP_SEQ_FEATURE | repeat:1 | RT | 68 | 4.9E-20 | 5.7E-18 |
| UP_SEQ_FEATURE | repeat:6 | RT | 50 | 7.2E-20 | 8.2E-18 |

#### Functional Annotation Clustering

Help and Manual

**Current Gene List: HLA-A1101_enr**
**Current Background: Homo sapiens**
**55 DAVID IDs**

Options Classification Stringency Medium

Rerun using options | Create Sublist

**8 Cluster(s)**

Download File

| | Annotation Cluster 1 | Enrichment Score: 7.04 | | | Count | P_Value | Benjamini |
|---|---|---|---|---|---|---|---|
| | GOTERM_MF_DIRECT | structural constituent of ribosome | RT | | 12 | 3.0E-11 | 2.1E-9 |
| | UP_KEYWORDS | Ribosomal protein | RT | | 11 | 4.6E-11 | 4.8E-9 |
| | KEGG_PATHWAY | Ribosome | RT | | 11 | 9.5E-11 | 1.0E-8 |
| | GOTERM_BP_DIRECT | SRP-dependent cotranslational protein targeting to membrane | RT | | 9 | 1.6E-10 | 4.7E-8 |
| | GOTERM_BP_DIRECT | viral transcription | RT | | 9 | 6.4E-10 | 9.7E-8 |
| | GOTERM_CC_DIRECT | cytosolic large ribosomal subunit | RT | | 8 | 7.5E-10 | 5.5E-8 |
| | GOTERM_CC_DIRECT | ribosome | RT | | 10 | 7.6E-10 | 2.8E-8 |
| | GOTERM_BP_DIRECT | nuclear-transcribed mRNA catabolic process, nonsense-mediated decay | RT | | 9 | 1.0E-9 | 1.1E-7 |
| | GOTERM_BP_DIRECT | translation | RT | | 11 | 1.3E-9 | 9.6E-8 |
| | GOTERM_BP_DIRECT | translational initiation | RT | | 9 | 3.2E-9 | 1.9E-7 |
| | UP_KEYWORDS | Ribonucleoprotein | RT | | 11 | 4.5E-9 | 2.3E-7 |
| | GOTERM_BP_DIRECT | rRNA processing | RT | | 10 | 5.5E-9 | 2.8E-7 |
| | GOTERM_MF_DIRECT | poly(A) RNA binding | RT | | 14 | 1.4E-5 | 4.7E-4 |
| | GOTERM_CC_DIRECT | focal adhesion | RT | | 5 | 2.4E-2 | 2.0E-1 |
| | GOTERM_CC_DIRECT | cytosol | RT | | 14 | 1.2E-1 | 5.3E-1 |
| | GOTERM_MF_DIRECT | RNA binding | RT | | 4 | 2.1E-1 | 9.3E-1 |
| | GOTERM_CC_DIRECT | membrane | RT | | 9 | 2.7E-1 | 7.9E-1 |
| | **Annotation Cluster 2** | **Enrichment Score: 4.06** | | | **Count** | **P_Value** | **Benjamini** |
| | SMART | H15 | RT | | 6 | 1.7E-11 | 2.6E-10 |
| | INTERPRO | Histone H5 | RT | | 6 | 2.0E-11 | 1.7E-9 |
| | INTERPRO | Histone H1/H5 | RT | | 6 | 3.1E-10 | 1.4E-8 |
| | UP_KEYWORDS | Citrullination | RT | | 7 | 2.0E-7 | 7.0E-6 |
| | GOTERM_CC_DIRECT | nucleosome | RT | | 6 | 6.4E-6 | 1.6E-4 |
| | GOTERM_BP_DIRECT | nucleosome assembly | RT | | 6 | 1.8E-5 | 6.7E-4 |
| | GOTERM_MF_DIRECT | chromatin DNA binding | RT | | 5 | 2.4E-5 | 5.4E-4 |
| | GOTERM_BP_DIRECT | histone H3-K27 trimethylation | RT | | 3 | 1.1E-4 | 3.7E-3 |
| | GOTERM_BP_DIRECT | nucleosome positioning | RT | | 3 | 2.0E-4 | 5.6E-3 |
| | INTERPRO | Winged helix-turn-helix DNA-binding domain | RT | | 6 | 5.5E-4 | 1.2E-2 |
| | GOTERM_BP_DIRECT | histone H3-K4 trimethylation | RT | | 3 | 7.5E-4 | 1.7E-2 |
| | UP_SEQ_FEATURE | region of interest:Globular | RT | | 3 | 8.0E-4 | 1.2E-1 |
| | GOTERM_CC_DIRECT | nuclear euchromatin | RT | | 3 | 2.6E-3 | 4.7E-2 |
| | UP_KEYWORDS | Chromosome | RT | | 6 | 3.9E-3 | 5.7E-2 |
| | GOTERM_CC_DIRECT | nuclear chromatin | RT | | 3 | 1.0E-1 | 4.8E-1 |
| | GOTERM_BP_DIRECT | negative regulation of transcription from RNA polymerase II promoter | RT | | 5 | 1.3E-1 | 8.5E-1 |
| | UP_KEYWORDS | Isopeptide bond | RT | | 5 | 3.5E-1 | 9.0E-1 |
| | UP_KEYWORDS | DNA-binding | RT | | 6 | 6.4E-1 | 9.8E-1 |

| Annotation Cluster 3 | Enrichment Score: 2.94 | | Count | P_Value | Benjamini |
|---|---|---|---|---|---|
| GOTERM_BP_DIRECT | very long-chain fatty acid biosynthetic process | RT | 4 | 5.4E-6 | 2.3E-4 |
| KEGG_PATHWAY | Fatty acid elongation | RT | 4 | 1.5E-4 | 7.8E-3 |
| GOTERM_BP_DIRECT | fatty acid elongation, saturated fatty acid | RT | 3 | 1.5E-4 | 4.6E-3 |
| GOTERM_BP_DIRECT | fatty acid elongation, monounsaturated fatty acid | RT | 3 | 1.5E-4 | 4.6E-3 |
| GOTERM_BP_DIRECT | fatty acid elongation, polyunsaturated fatty acid | RT | 3 | 1.5E-4 | 4.6E-3 |
| INTERPRO | GNS1/SUR4 membrane protein | RT | 3 | 1.7E-4 | 4.9E-3 |
| GOTERM_MF_DIRECT | 3-oxo-arachidoyl-CoA synthase activity | RT | 3 | 1.7E-4 | 3.0E-3 |
| GOTERM_MF_DIRECT | 3-oxo-cerotoyl-CoA synthase activity | RT | 3 | 1.7E-4 | 3.0E-3 |
| GOTERM_MF_DIRECT | 3-oxo-lignoceronyl-CoA synthase activity | RT | 3 | 1.7E-4 | 3.0E-3 |
| GOTERM_MF_DIRECT | fatty acid elongase activity | RT | 3 | 1.7E-4 | 3.0E-3 |
| GOTERM_BP_DIRECT | sphingolipid biosynthetic process | RT | 4 | 3.1E-4 | 7.7E-3 |
| UP_KEYWORDS | Fatty acid biosynthesis | RT | 4 | 3.4E-4 | 8.9E-3 |
| UP_KEYWORDS | Lipid biosynthesis | RT | 5 | 7.5E-4 | 1.3E-2 |
| GOTERM_BP_DIRECT | unsaturated fatty acid biosynthetic process | RT | 3 | 7.5E-4 | 1.7E-2 |
| GOTERM_CC_DIRECT | integral component of endoplasmic reticulum membrane | RT | 4 | 3.1E-3 | 4.5E-2 |
| UP_KEYWORDS | Fatty acid metabolism | RT | 4 | 4.2E-3 | 4.8E-2 |
| GOTERM_BP_DIRECT | long-chain fatty-acyl-CoA biosynthetic process | RT | 3 | 5.9E-3 | 9.0E-2 |
| UP_KEYWORDS | Lipid metabolism | RT | 5 | 2.7E-2 | 2.3E-1 |
| UP_KEYWORDS | Endoplasmic reticulum | RT | 7 | 6.0E-2 | 3.9E-1 |
| GOTERM_CC_DIRECT | endoplasmic reticulum membrane | RT | 6 | 9.2E-2 | 4.8E-1 |
| UP_KEYWORDS | Transferase | RT | 8 | 1.6E-1 | 7.0E-1 |
| GOTERM_CC_DIRECT | endoplasmic reticulum | RT | 4 | 4.1E-1 | 9.1E-1 |
| **Annotation Cluster 4** | **Enrichment Score: 1.22** | | **Count** | **P_Value** | **Benjamini** |
| GOTERM_MF_DIRECT | cAMP-dependent protein kinase activity | RT | 3 | 1.7E-4 | 3.0E-3 |
| KEGG_PATHWAY | Taste transduction | RT | 4 | 8.5E-4 | 3.0E-2 |
| GOTERM_BP_DIRECT | activation of protein kinase A activity | RT | 3 | 1.1E-3 | 2.3E-2 |
| GOTERM_CC_DIRECT | ciliary base | RT | 3 | 2.6E-3 | 4.7E-2 |
| KEGG_PATHWAY | Parkinson's disease | RT | 5 | 2.8E-3 | 7.1E-2 |
| GOTERM_BP_DIRECT | renal water homeostasis | RT | 3 | 3.5E-3 | 6.3E-2 |
| SMART | S_TK_X | RT | 3 | 3.9E-3 | 2.9E-2 |
| UP_KEYWORDS | cAMP | RT | 3 | 4.0E-3 | 5.1E-2 |
| GOTERM_BP_DIRECT | lipoprotein metabolic process | RT | 3 | 4.8E-3 | 8.3E-2 |
| GOTERM_BP_DIRECT | cellular response to glucagon stimulus | RT | 3 | 5.4E-3 | 8.6E-2 |
| KEGG_PATHWAY | Hedgehog signaling pathway | RT | 3 | 5.6E-3 | 1.1E-1 |
| KEGG_PATHWAY | Prion diseases | RT | 3 | 8.9E-3 | 1.5E-1 |
| UP_SEQ_FEATURE | domain:AGC-kinase C-terminal | RT | 3 | 9.6E-3 | 5.5E-1 |
| KEGG_PATHWAY | Amoebiasis | RT | 4 | 9.7E-3 | 1.4E-1 |
| INTERPRO | AGC-kinase, C-terminal | RT | 3 | 1.1E-2 | 1.8E-1 |
| KEGG_PATHWAY | Vasopressin-regulated water reabsorption | RT | 3 | 1.5E-2 | 1.8E-1 |
| KEGG_PATHWAY | Endocrine and other factor-regulated calcium reabsorption | RT | 3 | 1.5E-2 | 1.7E-1 |
| KEGG_PATHWAY | Cocaine addiction | RT | 3 | 1.8E-2 | 1.8E-1 |
| KEGG_PATHWAY | Ovarian steroidogenesis | RT | 3 | 1.8E-2 | 1.8E-1 |
| KEGG_PATHWAY | Vibrio cholerae infection | RT | 3 | 2.0E-2 | 1.8E-1 |

#### Functional Annotation Clustering

Help and Manual

**Current Gene List: HLA-A1101_depl**
**Current Background: Homo sapiens**
**106 DAVID IDs**

Options  Classification Stringency Medium

Rerun using options  Create Sublist

**22 Cluster(s)**  Download File

| Annotation Cluster 1 | Enrichment Score: 12.89 | | | Count | P_Value | Benjamini |
|---|---|---|---|---|---|---|
| UP_SEQ_FEATURE | signal peptide | RT | | 59 | 2.9E-20 | 2.4E-17 |
| UP_KEYWORDS | Signal | RT | | 61 | 7.3E-17 | 4.0E-15 |
| UP_KEYWORDS | Secreted | RT | | 42 | 5.9E-16 | 1.6E-14 |
| GOTERM_CC_DIRECT | extracellular region | RT | | 34 | 5.4E-12 | 1.5E-10 |
| UP_KEYWORDS | Glycoprotein | RT | | 56 | 1.1E-11 | 2.0E-10 |
| UP_KEYWORDS | Disulfide bond | RT | | 48 | 1.4E-11 | 2.3E-10 |
| UP_SEQ_FEATURE | glycosylation site:N-linked (GlcNAc...) | RT | | 53 | 2.4E-11 | 2.3E-9 |
| UP_SEQ_FEATURE | disulfide bond | RT | | 40 | 3.2E-9 | 1.3E-7 |

| Annotation Cluster 2 | Enrichment Score: 9.09 | | | Count | P_Value | Benjamini |
|---|---|---|---|---|---|---|
| GOTERM_MF_DIRECT | extracellular matrix structural constituent | RT | | 20 | 5.0E-29 | 6.1E-27 |
| UP_KEYWORDS | Extracellular matrix | RT | | 24 | 2.3E-22 | 3.3E-20 |
| UP_KEYWORDS | Hydroxylation | RT | | 16 | 1.1E-18 | 5.5E-17 |
| GOTERM_BP_DIRECT | extracellular matrix organization | RT | | 19 | 1.3E-17 | 7.2E-15 |
| GOTERM_CC_DIRECT | proteinaceous extracellular matrix | RT | | 20 | 2.3E-16 | 2.5E-14 |
| INTERPRO | Collagen triple helix repeat | RT | | 14 | 2.7E-16 | 5.9E-15 |
| UP_SEQ_FEATURE | region of interest:Triple-helical region | RT | | 10 | 1.2E-15 | 3.5E-13 |
| KEGG_PATHWAY | Protein digestion and absorption | RT | | 13 | 1.6E-15 | 5.1E-14 |
| UP_KEYWORDS | Collagen | RT | | 14 | 2.8E-15 | 6.7E-14 |
| KEGG_PATHWAY | ECM-receptor interaction | RT | | 12 | 7.2E-14 | 1.2E-12 |
| GOTERM_CC_DIRECT | endoplasmic reticulum lumen | RT | | 16 | 1.1E-13 | 5.9E-12 |
| GOTERM_BP_DIRECT | collagen catabolic process | RT | | 11 | 1.4E-12 | 4.0E-10 |
| UP_SEQ_FEATURE | domain:Fibrillar collagen NC1 | RT | | 7 | 6.7E-12 | 7.2E-10 |
| INTERPRO | Fibrillar collagen, C-terminal | RT | | 7 | 7.4E-12 | 1.3E-10 |
| SMART | COLFI | RT | | 7 | 4.9E-11 | 6.4E-10 |
| GOTERM_CC_DIRECT | collagen trimer | RT | | 11 | 6.3E-11 | 1.4E-9 |
| GOTERM_BP_DIRECT | skeletal system development | RT | | 12 | 1.7E-10 | 3.2E-8 |
| KEGG_PATHWAY | Amoebiasis | RT | | 10 | 7.3E-10 | 8.0E-9 |
| KEGG_PATHWAY | Focal adhesion | RT | | 12 | 9.9E-10 | 8.2E-9 |
| GOTERM_BP_DIRECT | collagen fibril organization | RT | | 8 | 1.3E-9 | 1.9E-7 |
| UP_SEQ_FEATURE | propeptide:C-terminal propeptide | RT | | 5 | 4.3E-8 | 1.2E-6 |
| KEGG_PATHWAY | PI3K-Akt signaling pathway | RT | | 12 | 2.1E-7 | 1.4E-6 |
| UP_SEQ_FEATURE | domain:VWFC | RT | | 5 | 1.8E-6 | 2.9E-5 |
| UP_SEQ_FEATURE | propeptide:N-terminal propeptide | RT | | 4 | 2.5E-6 | 3.8E-5 |
| GOTERM_MF_DIRECT | platelet-derived growth factor binding | RT | | 4 | 2.0E-5 | 6.0E-4 |
| KEGG_PATHWAY | Platelet activation | RT | | 7 | 2.5E-5 | 1.4E-4 |
| UP_KEYWORDS | Ehlers-Danlos syndrome | RT | | 4 | 4.5E-5 | 5.4E-4 |

| | | | | | Count | P_Value | Benjamini |
|---|---|---|---|---|---|---|---|
| ☐ | UP_SEQ_FEATURE | region of interest:Nonhelical region (C-terminal) | RT | | 3 | 1.5E-4 | 1.9E-3 |
| ☐ | UP_SEQ_FEATURE | domain:TSP N-terminal | RT | | 4 | 2.1E-4 | 2.5E-3 |
| ☐ | INTERPRO | Laminin G domain | RT | | 5 | 2.6E-4 | 2.5E-3 |
| ☐ | SMART | TSPN | RT | | 4 | 5.6E-4 | 2.7E-3 |
| ☐ | UP_KEYWORDS | Disease mutation | RT | | 26 | 8.8E-4 | 6.7E-3 |
| ☐ | UP_SEQ_FEATURE | glycosylation site:O-linked (Gal...) | RT | | 3 | 9.0E-4 | 9.8E-3 |
| ☐ | GOTERM_BP_DIRECT | cellular response to amino acid stimulus | RT | | 4 | 2.0E-3 | 1.1E-1 |
| ☐ | INTERPRO | Concanavalin A-like lectin/glucanase, subgroup | RT | | 6 | 7.1E-3 | 4.1E-2 |
| ☐ | GOTERM_BP_DIRECT | skin development | RT | | 3 | 1.4E-2 | 3.8E-1 |
| ☐ | GOTERM_BP_DIRECT | sensory perception of sound | RT | | 3 | 1.6E-1 | 9.6E-1 |
| ☐ | GOTERM_BP_DIRECT | regulation of immune response | RT | | 3 | 2.5E-1 | 9.9E-1 |
| | **Annotation Cluster 3** | **Enrichment Score: 6.56** | G | | **Count** | **P_Value** | **Benjamini** |
| ☐ | UP_SEQ_FEATURE | domain:EGF-like 3 | RT | | 11 | 5.3E-12 | 6.5E-10 |
| ☐ | UP_SEQ_FEATURE | domain:EGF-like 6 | RT | | 9 | 5.1E-11 | 4.0E-9 |
| ☐ | UP_SEQ_FEATURE | domain:EGF-like 19 | RT | | 5 | 4.3E-8 | 1.2E-6 |
| ☐ | UP_SEQ_FEATURE | domain:EGF-like 5 | RT | | 7 | 6.7E-8 | 1.8E-6 |
| ☐ | UP_SEQ_FEATURE | domain:EGF-like 9 | RT | | 6 | 1.2E-7 | 2.7E-6 |
| ☐ | UP_SEQ_FEATURE | domain:EGF-like 7 | RT | | 5 | 1.2E-5 | 1.6E-4 |
| ☐ | INTERPRO | EGF-like, laminin | RT | | 4 | 1.1E-3 | 8.1E-3 |
| ☐ | SMART | EGF_Lam | RT | | 3 | 2.6E-2 | 7.3E-2 |

### Functional Annotation Clustering

Help and Manual

**Current Gene List: HLA-B0702_enr**
**Current Background: Homo sapiens**
**614 DAVID IDs**

⊞ Options   Classification Stringency [Medium]

[Rerun using options] [Create Sublist]

**64 Cluster(s)**   Download File

| Category | Term | | Count | P_Value | Benjamini |
|---|---|---|---|---|---|
| **Annotation Cluster 1** | **Enrichment Score: 21.3** | | **Count** | **P_Value** | **Benjamini** |
| INTERPRO | Homeobox, conserved site | RT | 43 | 3.0E-26 | 1.9E-23 |
| GOTERM_MF_DIRECT | sequence-specific DNA binding | RT | 67 | 3.9E-25 | 1.9E-22 |
| SMART | HOX | RT | 47 | 1.2E-24 | 1.6E-22 |
| INTERPRO | Homeodomain | RT | 47 | 1.4E-24 | 4.5E-22 |
| UP_SEQ_FEATURE | DNA-binding region:Homeobox | RT | 41 | 3.0E-23 | 1.8E-20 |
| UP_KEYWORDS | Homeobox | RT | 47 | 1.3E-22 | 4.2E-20 |
| INTERPRO | Homeodomain-like | RT | 49 | 3.6E-21 | 7.7E-19 |
| INTERPRO | Homeodomain, metazoa | RT | 25 | 2.7E-17 | 4.3E-15 |
| UP_KEYWORDS | Developmental protein | RT | 73 | 2.8E-13 | 1.8E-11 |
| **Annotation Cluster 2** | **Enrichment Score: 11.43** | | **Count** | **P_Value** | **Benjamini** |
| UP_KEYWORDS | DNA-binding | RT | 142 | 4.3E-22 | 7.1E-20 |
| UP_KEYWORDS | Transcription regulation | RT | 139 | 1.0E-15 | 1.1E-13 |
| UP_KEYWORDS | Transcription | RT | 140 | 3.9E-15 | 3.2E-13 |
| GOTERM_MF_DIRECT | transcription factor activity, sequence-specific DNA binding | RT | 67 | 2.4E-11 | 5.6E-9 |
| GOTERM_BP_DIRECT | transcription, DNA-templated | RT | 106 | 3.1E-11 | 5.2E-8 |
| UP_KEYWORDS | Nucleus | RT | 216 | 5.5E-8 | 2.6E-6 |
| GOTERM_CC_DIRECT | nucleus | RT | 214 | 2.3E-7 | 6.3E-5 |
| GOTERM_MF_DIRECT | DNA binding | RT | 81 | 2.5E-6 | 2.4E-4 |
| **Annotation Cluster 3** | **Enrichment Score: 10.36** | | **Count** | **P_Value** | **Benjamini** |
| INTERPRO | Transcription factor, fork head, conserved site | RT | 15 | 9.9E-14 | 1.3E-11 |
| INTERPRO | Transcription factor, fork head | RT | 16 | 2.1E-12 | 2.2E-10 |
| SMART | FH | RT | 16 | 3.8E-12 | 2.6E-10 |
| UP_SEQ_FEATURE | DNA-binding region:Fork-head | RT | 16 | 6.1E-12 | 1.8E-9 |
| GOTERM_MF_DIRECT | RNA polymerase II transcription factor activity, sequence-specific DNA binding | RT | 24 | 6.5E-10 | 1.0E-7 |
| INTERPRO | Winged helix-turn-helix DNA-binding domain | RT | 22 | 2.3E-6 | 1.6E-4 |
| **Annotation Cluster 4** | **Enrichment Score: 6.96** | | **Count** | **P_Value** | **Benjamini** |
| GOTERM_BP_DIRECT | transcription from RNA polymerase II promoter | RT | 44 | 1.7E-10 | 1.5E-7 |
| GOTERM_BP_DIRECT | positive regulation of transcription from RNA polymerase II promoter | RT | 55 | 1.8E-6 | 4.4E-4 |
| GOTERM_MF_DIRECT | transcriptional activator activity, RNA polymerase II core promoter proximal region sequence-specific binding | RT | 22 | 4.2E-6 | 3.4E-4 |
| **Annotation Cluster 5** | **Enrichment Score: 4.56** | | **Count** | **P_Value** | **Benjamini** |
| INTERPRO | Myc-type, basic helix-loop-helix (bHLH) domain | RT | 16 | 6.7E-7 | 6.2E-5 |
| SMART | HLH | RT | 16 | 8.2E-7 | 3.8E-5 |
| UP_SEQ_FEATURE | DNA-binding region:Basic motif | RT | 19 | 1.2E-6 | 2.4E-4 |
| UP_SEQ_FEATURE | domain:Helix-loop-helix motif | RT | 15 | 5.9E-6 | 1.0E-3 |
| INTERPRO | Orange | RT | 6 | 1.0E-5 | 6.0E-4 |

#### Functional Annotation Clustering

Help and Manual

**Current Gene List: HLA-B0702_depl**
**Current Background: Homo sapiens**
**220 DAVID IDs**

Options    Classification Stringency [Medium]

[Rerun using options] [Create Sublist]

**39 Cluster(s)**

Download File

| Annotation Cluster 1 | Enrichment Score: 5.73 | | Count | P_Value | Benjamini |
|---|---|---|---|---|---|
| UP_SEQ_FEATURE | zinc finger region:C2H2-type 10 | RT | 23 | 2.2E-11 | 2.2E-8 |
| UP_SEQ_FEATURE | zinc finger region:C2H2-type 6 | RT | 27 | 4.0E-11 | 2.0E-8 |
| UP_SEQ_FEATURE | zinc finger region:C2H2-type 7 | RT | 26 | 4.2E-11 | 1.4E-8 |
| UP_SEQ_FEATURE | zinc finger region:C2H2-type 11 | RT | 21 | 1.1E-10 | 2.2E-8 |
| UP_SEQ_FEATURE | domain:KRAB | RT | 22 | 1.7E-10 | 2.8E-8 |
| UP_SEQ_FEATURE | zinc finger region:C2H2-type 9 | RT | 23 | 2.3E-10 | 3.2E-8 |
| UP_SEQ_FEATURE | zinc finger region:C2H2-type 8 | RT | 24 | 2.8E-10 | 3.4E-8 |
| INTERPRO | Krueppel-associated box | RT | 23 | 5.0E-10 | 7.9E-8 |
| UP_SEQ_FEATURE | zinc finger region:C2H2-type 12 | RT | 18 | 3.0E-9 | 3.2E-7 |
| UP_SEQ_FEATURE | zinc finger region:C2H2-type 5 | RT | 25 | 7.3E-9 | 7.2E-7 |
| INTERPRO | Zinc finger C2H2-type/integrase DNA-binding domain | RT | 28 | 1.5E-8 | 1.2E-6 |
| INTERPRO | Zinc finger, C2H2 | RT | 29 | 4.2E-8 | 2.7E-6 |
| INTERPRO | Zinc finger, C2H2-like | RT | 28 | 6.0E-8 | 3.2E-6 |
| SMART | KRAB | RT | 23 | 9.4E-8 | 9.2E-6 |
| GOTERM_MF_DIRECT | nucleic acid binding | RT | 31 | 1.4E-7 | 3.0E-5 |
| UP_SEQ_FEATURE | zinc finger region:C2H2-type 13 | RT | 14 | 3.6E-7 | 2.4E-5 |
| UP_SEQ_FEATURE | zinc finger region:C2H2-type 14 | RT | 12 | 1.1E-6 | 6.3E-5 |
| UP_SEQ_FEATURE | zinc finger region:C2H2-type 3 | RT | 23 | 1.7E-6 | 8.3E-5 |
| UP_SEQ_FEATURE | zinc finger region:C2H2-type 4 | RT | 22 | 1.8E-6 | 8.3E-5 |
| UP_SEQ_FEATURE | zinc finger region:C2H2-type 16 | RT | 10 | 1.8E-6 | 8.0E-5 |
| UP_KEYWORDS | Zinc-finger | RT | 42 | 1.9E-6 | 4.5E-5 |
| GOTERM_MF_DIRECT | DNA binding | RT | 39 | 4.1E-6 | 3.0E-4 |
| UP_SEQ_FEATURE | zinc finger region:C2H2-type 15 | RT | 10 | 1.5E-5 | 5.5E-4 |
| GOTERM_BP_DIRECT | regulation of transcription, DNA-templated | RT | 34 | 2.4E-5 | 2.0E-2 |
| SMART | ZnF_C2H2 | RT | 28 | 2.5E-5 | 6.1E-4 |
| UP_KEYWORDS | Zinc | RT | 47 | 2.9E-5 | 4.4E-4 |
| UP_SEQ_FEATURE | zinc finger region:C2H2-type 2 | RT | 20 | 4.2E-5 | 1.5E-3 |
| UP_KEYWORDS | Metal-binding | RT | 62 | 1.2E-4 | 1.7E-3 |
| UP_KEYWORDS | Nucleus | RT | 81 | 1.8E-4 | 2.2E-3 |
| GOTERM_MF_DIRECT | metal ion binding | RT | 40 | 2.0E-4 | 4.8E-3 |
| UP_SEQ_FEATURE | zinc finger region:C2H2-type 17 | RT | 7 | 2.6E-4 | 7.8E-3 |
| UP_SEQ_FEATURE | zinc finger region:C2H2-type 1 | RT | 16 | 1.1E-3 | 2.7E-2 |
| UP_KEYWORDS | DNA-binding | RT | 37 | 1.8E-3 | 1.8E-2 |
| UP_SEQ_FEATURE | zinc finger region:C2H2-type 18 | RT | 5 | 5.4E-3 | 9.5E-2 |
| GOTERM_CC_DIRECT | intracellular | RT | 26 | 5.9E-3 | 7.9E-2 |
| GOTERM_BP_DIRECT | transcription, DNA-templated | RT | 32 | 9.5E-3 | 3.7E-1 |
| UP_KEYWORDS | Transcription regulation | RT | 37 | 1.4E-2 | 8.2E-2 |

| Annotation Cluster 2 | Enrichment Score: 5.37 | G | | Count | P_Value | Benjamini |
|---|---|---|---|---|---|---|
| INTERPRO | Myosin, N-terminal, SH3-like | RT | | 8 | 9.4E-11 | 3.0E-8 |
| GOTERM_CC_DIRECT | muscle myosin complex | RT | | 8 | 1.8E-10 | 4.5E-8 |
| GOTERM_CC_DIRECT | myosin filament | RT | | 8 | 1.8E-10 | 4.5E-8 |
| UP_KEYWORDS | Thick filament | RT | | 8 | 2.5E-10 | 1.8E-8 |
| INTERPRO | Myosin tail | RT | | 8 | 7.1E-10 | 7.6E-8 |
| INTERPRO | Myosin-like IQ motif-containing domain | RT | | 8 | 7.1E-10 | 7.6E-8 |
| UP_SEQ_FEATURE | region of interest:Actin-binding | RT | | 8 | 2.0E-8 | 1.8E-6 |
| UP_SEQ_FEATURE | domain:Myosin head-like | RT | | 8 | 1.2E-7 | 1.0E-5 |
| INTERPRO | Myosin head, motor domain | RT | | 8 | 2.2E-7 | 9.9E-6 |
| UP_KEYWORDS | Motor protein | RT | | 12 | 2.2E-7 | 9.2E-6 |
| UP_SEQ_FEATURE | domain:IQ | RT | | 9 | 3.0E-7 | 2.1E-5 |
| UP_KEYWORDS | Myosin | RT | | 8 | 1.1E-6 | 3.3E-5 |
| GOTERM_MF_DIRECT | microfilament motor activity | RT | | 6 | 1.7E-6 | 1.8E-4 |
| SMART | MYSc | RT | | 8 | 1.9E-6 | 9.4E-5 |
| GOTERM_CC_DIRECT | sarcomere | RT | | 7 | 3.9E-6 | 3.2E-4 |
| UP_KEYWORDS | Muscle protein | RT | | 8 | 4.2E-6 | 8.1E-5 |
| INTERPRO | IQ motif, EF-hand binding site | RT | | 9 | 4.6E-6 | 1.5E-4 |
| UP_KEYWORDS | Actin-binding | RT | | 14 | 8.1E-6 | 1.3E-4 |
| KEGG_PATHWAY | Tight junction | RT | | 7 | 1.4E-5 | 1.1E-3 |
| GOTERM_BP_DIRECT | muscle filament sliding | RT | | 6 | 4.2E-5 | 1.7E-2 |
| GOTERM_MF_DIRECT | motor activity | RT | | 7 | 6.8E-5 | 3.0E-3 |
| INTERPRO | P-loop containing nucleoside triphosphate hydrolase | RT | | 24 | 8.4E-5 | 2.4E-3 |
| GOTERM_BP_DIRECT | muscle contraction | RT | | 8 | 1.2E-4 | 3.2E-2 |
| GOTERM_MF_DIRECT | ATPase activity | RT | | 10 | 1.4E-4 | 4.4E-3 |
| GOTERM_MF_DIRECT | calmodulin binding | RT | | 10 | 1.8E-4 | 4.9E-3 |
| GOTERM_CC_DIRECT | myosin complex | RT | | 6 | 2.1E-4 | 8.8E-3 |
| UP_KEYWORDS | Calmodulin-binding | RT | | 9 | 2.3E-4 | 2.7E-3 |
| GOTERM_BP_DIRECT | ATP metabolic process | RT | | 5 | 3.1E-4 | 6.2E-2 |
| GOTERM_MF_DIRECT | myosin phosphatase activity | RT | | 3 | 6.5E-4 | 1.2E-2 |
| GOTERM_BP_DIRECT | skeletal muscle contraction | RT | | 4 | 1.9E-3 | 1.6E-1 |
| SMART | IQ | RT | | 5 | 7.7E-3 | 8.1E-2 |
| GOTERM_MF_DIRECT | actin binding | RT | | 9 | 9.3E-3 | 1.2E-1 |
| GOTERM_CC_DIRECT | myofibril | RT | | 3 | 3.8E-2 | 3.2E-1 |
| GOTERM_BP_DIRECT | protein dephosphorylation | RT | | 4 | 1.4E-1 | 9.3E-1 |
| **Annotation Cluster 3** | **Enrichment Score: 4.86** | **G** | | **Count** | **P_Value** | **Benjamini** |
| INTERPRO | Intermediate filament protein, conserved site | RT | | 9 | 4.3E-7 | 1.7E-5 |
| UP_SEQ_FEATURE | region of interest:Coil 2 | RT | | 9 | 7.0E-7 | 4.3E-5 |
| UP_SEQ_FEATURE | region of interest:Linker 12 | RT | | 9 | 7.0E-7 | 4.3E-5 |
| UP_SEQ_FEATURE | region of interest:Coil 1A | RT | | 9 | 1.3E-6 | 7.3E-5 |
| UP_SEQ_FEATURE | region of interest:Coil 1B | RT | | 9 | 1.3E-6 | 7.3E-5 |
| UP_SEQ_FEATURE | region of interest:Linker 1 | RT | | 9 | 1.3E-6 | 7.3E-5 |
| UP_SEQ_FEATURE | region of interest:Rod | RT | | 9 | 1.5E-6 | 7.7E-5 |
| UP_KEYWORDS | Intermediate filament | RT | | 9 | 1.6E-6 | 4.3E-5 |
| UP_SEQ_FEATURE | region of interest:Head | RT | | 9 | 1.8E-6 | 7.8E-5 |

### Functional Annotation Clustering

Help and Manual

**Current Gene List: HLA-B2705_enr**
**Current Background: Homo sapiens**
**671 DAVID IDs**

⊞ Options   Classification Stringency [Medium]

[Rerun using options] [Create Sublist]

**63 Cluster(s)**   Download File

| Annotation Cluster 1 | Enrichment Score: 29.33 | | Count | P_Value | Benjamini |
|---|---|---|---|---|---|
| UP_KEYWORDS | Ribosomal protein | RT | 67 | 1.1E-50 | 3.3E-48 |
| KEGG_PATHWAY | Ribosome | RT | 56 | 1.9E-45 | 2.7E-43 |
| GOTERM_MF_DIRECT | structural constituent of ribosome | RT | 66 | 1.7E-43 | 8.2E-41 |
| GOTERM_CC_DIRECT | ribosome | RT | 58 | 2.7E-43 | 7.6E-41 |
| UP_KEYWORDS | Ribonucleoprotein | RT | 73 | 2.5E-42 | 3.9E-40 |
| GOTERM_BP_DIRECT | translation | RT | 62 | 1.0E-36 | 1.6E-33 |
| GOTERM_BP_DIRECT | SRP-dependent cotranslational protein targeting to membrane | RT | 39 | 1.4E-32 | 5.3E-30 |
| GOTERM_CC_DIRECT | cytosolic large ribosomal subunit | RT | 33 | 5.5E-30 | 7.8E-28 |
| GOTERM_BP_DIRECT | nuclear-transcribed mRNA catabolic process, nonsense-mediated decay | RT | 40 | 2.6E-29 | 6.7E-27 |
| GOTERM_BP_DIRECT | viral transcription | RT | 39 | 3.1E-29 | 6.9E-27 |
| GOTERM_BP_DIRECT | rRNA processing | RT | 50 | 1.6E-28 | 3.2E-26 |
| GOTERM_BP_DIRECT | translational initiation | RT | 40 | 1.0E-26 | 1.6E-24 |
| GOTERM_MF_DIRECT | RNA binding | RT | 49 | 8.2E-10 | 6.4E-8 |
| GOTERM_CC_DIRECT | focal adhesion | RT | 20 | 4.8E-2 | 3.5E-1 |
| GOTERM_CC_DIRECT | cytosol | RT | 71 | 1.0E0 | 1.0E0 |
| **Annotation Cluster 2** | **Enrichment Score: 14.45** | | **Count** | **P_Value** | **Benjamini** |
| GOTERM_BP_DIRECT | telomere organization | RT | 26 | 5.7E-36 | 4.4E-33 |
| GOTERM_BP_DIRECT | chromatin silencing at rDNA | RT | 29 | 4.4E-35 | 2.3E-32 |
| UP_KEYWORDS | Nucleosome core | RT | 39 | 4.4E-33 | 4.6E-31 |
| UP_KEYWORDS | Citrullination | RT | 40 | 1.1E-32 | 8.4E-31 |
| GOTERM_BP_DIRECT | negative regulation of gene expression, epigenetic | RT | 29 | 2.1E-29 | 6.6E-27 |
| GOTERM_BP_DIRECT | DNA replication-dependent nucleosome assembly | RT | 24 | 4.0E-28 | 7.0E-26 |
| GOTERM_CC_DIRECT | nucleosome | RT | 35 | 4.3E-27 | 4.0E-25 |
| GOTERM_BP_DIRECT | positive regulation of gene expression, epigenetic | RT | 29 | 5.8E-26 | 8.2E-24 |
| GOTERM_BP_DIRECT | protein heterotetramerization | RT | 25 | 1.2E-25 | 1.5E-23 |
| INTERPRO | Histone-fold | RT | 36 | 1.3E-25 | 1.0E-22 |
| GOTERM_CC_DIRECT | nuclear chromosome | RT | 26 | 1.0E-23 | 7.1E-22 |
| GOTERM_BP_DIRECT | nucleosome assembly | RT | 35 | 1.7E-23 | 2.0E-21 |
| SMART | H3 | RT | 17 | 2.4E-23 | 3.6E-21 |
| KEGG_PATHWAY | Systemic lupus erythematosus | RT | 37 | 1.4E-22 | 9.9E-21 |
| SMART | H4 | RT | 15 | 6.5E-22 | 4.9E-20 |
| INTERPRO | Histone H3 | RT | 17 | 6.7E-22 | 2.7E-19 |
| INTERPRO | Histone H4 | RT | 15 | 1.2E-20 | 3.2E-18 |
| INTERPRO | Histone H4, conserved site | RT | 14 | 3.5E-19 | 7.2E-17 |
| UP_SEQ_FEATURE | chain:Histone H4 | RT | 14 | 3.8E-19 | 4.4E-16 |
| SMART | TAF | RT | 14 | 9.4E-19 | 4.7E-17 |

| | | | | | |
|---|---|---|---|---|---|
| UP_SEQ_FEATURE | chain:Histone H3.1 | RT | 13 | 7.7E-17 | 6.6E-14 |
| UP_SEQ_FEATURE | chain:Histone H2A type 1-D | RT | 13 | 7.7E-17 | 6.6E-14 |
| GOTERM_BP_DIRECT | DNA replication-independent nucleosome assembly | RT | 16 | 1.4E-16 | 1.1E-14 |
| KEGG_PATHWAY | Alcoholism | RT | 35 | 1.9E-16 | 1.0E-14 |
| GOTERM_BP_DIRECT | negative regulation of megakaryocyte differentiation | RT | 14 | 2.0E-16 | 2.0E-14 |
| UP_SEQ_FEATURE | cross-link:Glycyl lysine isopeptide (Lys-Gly) (interchain with G-Cter in ubiquitin); alternate | RT | 14 | 2.0E-16 | 8.7E-14 |
| GOTERM_BP_DIRECT | telomere capping | RT | 15 | 5.1E-16 | 4.8E-14 |
| GOTERM_CC_DIRECT | nuclear chromosome, telomeric region | RT | 29 | 8.2E-16 | 4.4E-14 |
| GOTERM_CC_DIRECT | nuclear nucleosome | RT | 18 | 9.7E-15 | 4.6E-13 |
| UP_KEYWORDS | Chromosome | RT | 48 | 2.6E-14 | 1.6E-12 |
| GOTERM_BP_DIRECT | DNA-templated transcription, initiation | RT | 16 | 7.6E-14 | 5.7E-12 |
| GOTERM_MF_DIRECT | nucleosomal DNA binding | RT | 17 | 6.9E-13 | 8.0E-11 |
| GOTERM_BP_DIRECT | regulation of gene silencing | RT | 10 | 1.5E-12 | 1.1E-10 |
| INTERPRO | Histone core | RT | 18 | 2.3E-11 | 3.2E-9 |
| GOTERM_BP_DIRECT | CENP-A containing nucleosome assembly | RT | 15 | 2.8E-11 | 1.9E-9 |
| GOTERM_BP_DIRECT | beta-catenin-TCF complex assembly | RT | 15 | 2.8E-11 | 1.9E-9 |
| GOTERM_BP_DIRECT | double-strand break repair via nonhomologous end joining | RT | 15 | 7.4E-9 | 4.4E-7 |
| GOTERM_MF_DIRECT | protein heterodimerization activity | RT | 41 | 3.8E-8 | 2.6E-6 |
| GOTERM_BP_DIRECT | blood coagulation | RT | 19 | 2.2E-5 | 1.2E-3 |
| KEGG_PATHWAY | Transcriptional misregulation in cancer | RT | 19 | 2.4E-5 | 8.6E-4 |
| GOTERM_CC_DIRECT | protein complex | RT | 31 | 2.7E-5 | 5.9E-4 |
| GOTERM_CC_DIRECT | extracellular region | RT | 81 | 5.1E-5 | 1.0E-3 |
| GOTERM_CC_DIRECT | extracellular matrix | RT | 24 | 8.9E-5 | 1.7E-3 |
| KEGG_PATHWAY | Viral carcinogenesis | RT | 20 | 1.2E-4 | 3.3E-3 |
| UP_KEYWORDS | DNA-binding | RT | 95 | 4.1E-4 | 7.9E-3 |
| GOTERM_MF_DIRECT | protein domain specific binding | RT | 17 | 1.6E-3 | 6.0E-2 |
| UP_KEYWORDS | Methylation | RT | 50 | 3.0E-3 | 5.0E-2 |
| UP_KEYWORDS | Chromosomal rearrangement | RT | 22 | 3.2E-3 | 5.2E-2 |
| UP_SEQ_FEATURE | cross-link:Glycyl lysine isopeptide (Lys-Gly) (interchain with G-Cter in ubiquitin) | RT | 17 | 7.4E-3 | 4.2E-1 |
| GOTERM_BP_DIRECT | cell-cell adhesion | RT | 17 | 1.2E-2 | 3.1E-1 |
| GOTERM_MF_DIRECT | cadherin binding involved in cell-cell adhesion | RT | 17 | 3.3E-2 | 5.4E-1 |
| GOTERM_CC_DIRECT | cell-cell adherens junction | RT | 17 | 5.7E-2 | 3.9E-1 |
| UP_KEYWORDS | Ubl conjugation | RT | 62 | 2.3E-1 | 8.3E-1 |
| UP_KEYWORDS | Isopeptide bond | RT | 37 | 5.8E-1 | 9.8E-1 |
| GOTERM_CC_DIRECT | extracellular exosome | RT | 81 | 8.9E-1 | 1.0E0 |
| **Annotation Cluster 3** | **Enrichment Score: 11.01** | G | **Count** | **P_Value** | **Benjamini** |
| GOTERM_BP_DIRECT | mitochondrial translational elongation | RT | 23 | 1.2E-14 | 1.0E-12 |
| GOTERM_BP_DIRECT | mitochondrial translational termination | RT | 23 | 1.6E-14 | 1.3E-12 |
| GOTERM_CC_DIRECT | mitochondrial inner membrane | RT | 49 | 1.2E-13 | 4.9E-12 |
| GOTERM_CC_DIRECT | mitochondrial large ribosomal subunit | RT | 17 | 8.8E-13 | 3.1E-11 |
| UP_KEYWORDS | Mitochondrion | RT | 82 | 1.1E-11 | 5.5E-10 |
| UP_KEYWORDS | Transit peptide | RT | 43 | 1.6E-7 | 6.0E-6 |
| UP_SEQ_FEATURE | transit peptide:Mitochondrion | RT | 39 | 2.5E-7 | 5.9E-5 |

#### Functional Annotation Clustering

Help and Manual

**Current Gene List: HLA-B2705_depl**
**Current Background: Homo sapiens**
**239 DAVID IDs**

Options Classification Stringency Medium

Rerun using options | Create Sublist

**42 Cluster(s)**

Download File

| Annotation Cluster 1 | Enrichment Score: 9.42 | G | | | Count | P_Value | Benjamini |
|---|---|---|---|---|---|---|---|
| INTERPRO | Collagen triple helix repeat | RT | | | 28 | 4.2E-32 | 1.5E-29 |
| UP_KEYWORDS | Collagen | RT | | | 28 | 5.4E-30 | 1.2E-27 |
| GOTERM_MF_DIRECT | extracellular matrix structural constituent | RT | | | 24 | 1.3E-27 | 3.4E-25 |
| UP_KEYWORDS | Hydroxylation | RT | | | 26 | 2.8E-27 | 3.1E-25 |
| UP_KEYWORDS | Extracellular matrix | RT | | | 33 | 9.3E-24 | 6.9E-22 |
| KEGG_PATHWAY | Protein digestion and absorption | RT | | | 21 | 8.5E-22 | 6.4E-20 |
| GOTERM_CC_DIRECT | collagen trimer | RT | | | 22 | 4.1E-21 | 1.1E-18 |
| UP_SEQ_FEATURE | region of interest:Triple-helical region | RT | | | 14 | 5.4E-20 | 7.3E-17 |
| GOTERM_BP_DIRECT | collagen catabolic process | RT | | | 19 | 6.2E-20 | 6.5E-17 |
| GOTERM_CC_DIRECT | endoplasmic reticulum lumen | RT | | | 27 | 8.5E-20 | 1.1E-17 |
| GOTERM_BP_DIRECT | extracellular matrix organization | RT | | | 26 | 2.7E-18 | 1.4E-15 |
| GOTERM_CC_DIRECT | proteinaceous extracellular matrix | RT | | | 28 | 3.6E-17 | 3.0E-15 |
| KEGG_PATHWAY | ECM-receptor interaction | RT | | | 16 | 1.5E-14 | 5.5E-13 |
| UP_SEQ_FEATURE | domain:Fibrillar collagen NC1 | RT | | | 9 | 4.9E-14 | 6.5E-12 |
| INTERPRO | Fibrillar collagen, C-terminal | RT | | | 9 | 5.7E-14 | 9.9E-12 |
| SMART | COLFI | RT | | | 9 | 3.4E-13 | 3.6E-11 |
| KEGG_PATHWAY | Amoebiasis | RT | | | 15 | 5.5E-12 | 1.4E-10 |
| UP_SEQ_FEATURE | propeptide:C-terminal propeptide | RT | | | 7 | 6.5E-11 | 4.6E-9 |
| KEGG_PATHWAY | Focal adhesion | RT | | | 17 | 5.2E-10 | 9.7E-9 |
| UP_SEQ_FEATURE | domain:Collagen IV NC1 | RT | | | 6 | 1.2E-9 | 7.5E-8 |
| INTERPRO | Collagen IV, non-collagenous | RT | | | 6 | 1.4E-9 | 1.6E-7 |
| GOTERM_CC_DIRECT | collagen type IV trimer | RT | | | 6 | 1.6E-9 | 1.0E-7 |
| SMART | C4 | RT | | | 6 | 4.2E-9 | 2.2E-7 |
| UP_KEYWORDS | Basement membrane | RT | | | 9 | 1.6E-8 | 5.0E-7 |
| GOTERM_BP_DIRECT | collagen fibril organization | RT | | | 9 | 2.4E-8 | 8.4E-6 |
| UP_KEYWORDS | Secreted | RT | | | 51 | 7.2E-8 | 2.0E-6 |
| GOTERM_MF_DIRECT | platelet-derived growth factor binding | RT | | | 6 | 1.2E-7 | 1.1E-5 |
| BIOCARTA | Regulators of Bone Mineralization | RT | | | 6 | 1.4E-7 | 6.1E-6 |
| GOTERM_BP_DIRECT | skeletal system development | RT | | | 13 | 1.6E-7 | 3.4E-5 |
| BIOCARTA | Vitamin C in the Brain | RT | | | 6 | 2.5E-7 | 5.5E-6 |
| UP_SEQ_FEATURE | propeptide:N-terminal propeptide | RT | | | 5 | 2.7E-7 | 9.4E-6 |
| GOTERM_CC_DIRECT | extracellular region | RT | | | 45 | 4.0E-7 | 1.7E-5 |
| BIOCARTA | Angiotensin-converting enzyme 2 regulates heart function | RT | | | 6 | 6.7E-7 | 1.0E-5 |
| KEGG_PATHWAY | PI3K-Akt signaling pathway | RT | | | 17 | 7.8E-7 | 1.2E-5 |
| BIOCARTA | Platelet Amyloid Precursor Protein Pathway | RT | | | 6 | 1.5E-6 | 1.7E-5 |
| GOTERM_BP_DIRECT | cellular response to amino acid stimulus | RT | | | 8 | 1.8E-6 | 3.2E-4 |
| UP_SEQ_FEATURE | region of interest:Nonhelical region (C-terminal) | RT | | | 4 | 6.2E-6 | 1.8E-4 |

| Annotation Cluster 2 | Enrichment Score: 8.31 | | Count | P_Value | Benjamini |
|---|---|---|---|---|---|
| UP_SEQ_FEATURE | repeat:9 | RT | 19 | 3.6E-16 | 2.2E-13 |
| UP_SEQ_FEATURE | repeat:10 | RT | 18 | 1.5E-15 | 7.0E-13 |
| UP_SEQ_FEATURE | repeat:13 | RT | 17 | 2.2E-15 | 6.0E-13 |
| UP_SEQ_FEATURE | repeat:12 | RT | 17 | 3.2E-15 | 7.2E-13 |
| UP_SEQ_FEATURE | repeat:11 | RT | 17 | 5.8E-15 | 1.1E-12 |
| UP_SEQ_FEATURE | repeat:7 | RT | 19 | 2.0E-14 | 3.4E-12 |
| UP_SEQ_FEATURE | repeat:5 | RT | 20 | 4.8E-14 | 7.1E-12 |
| UP_SEQ_FEATURE | repeat:8 | RT | 18 | 8.7E-14 | 1.1E-11 |
| UP_SEQ_FEATURE | repeat:6 | RT | 19 | 1.2E-13 | 1.3E-11 |
| UP_SEQ_FEATURE | repeat:15 | RT | 15 | 1.4E-13 | 1.5E-11 |
| UP_SEQ_FEATURE | repeat:14 | RT | 15 | 4.5E-13 | 4.3E-11 |
| UP_SEQ_FEATURE | repeat:4 | RT | 19 | 1.2E-11 | 1.1E-9 |
| UP_SEQ_FEATURE | repeat:3 | RT | 20 | 2.0E-11 | 1.7E-9 |
| UP_SEQ_FEATURE | repeat:1 | RT | 21 | 2.5E-11 | 2.0E-9 |
| UP_SEQ_FEATURE | repeat:17 | RT | 12 | 4.0E-11 | 3.0E-9 |
| UP_SEQ_FEATURE | repeat:20 | RT | 11 | 1.1E-10 | 7.3E-9 |
| UP_SEQ_FEATURE | repeat:2 | RT | 20 | 2.4E-10 | 1.5E-8 |
| UP_SEQ_FEATURE | repeat:21 | RT | 10 | 1.4E-9 | 8.0E-8 |
| UP_SEQ_FEATURE | repeat:16 | RT | 11 | 2.1E-9 | 1.2E-7 |
| UP_SEQ_FEATURE | repeat:19 | RT | 10 | 3.9E-9 | 2.0E-7 |
| UP_SEQ_FEATURE | repeat:18 | RT | 10 | 4.7E-9 | 2.3E-7 |
| UP_SEQ_FEATURE | repeat:28 | RT | 8 | 1.5E-8 | 6.8E-7 |
| UP_SEQ_FEATURE | repeat:22 | RT | 9 | 2.1E-8 | 9.3E-7 |
| UP_SEQ_FEATURE | repeat:27 | RT | 8 | 3.4E-8 | 1.4E-6 |
| UP_SEQ_FEATURE | repeat:25 | RT | 8 | 1.1E-7 | 4.4E-6 |
| UP_SEQ_FEATURE | repeat:23 | RT | 8 | 2.1E-7 | 7.7E-6 |
| UP_SEQ_FEATURE | repeat:26 | RT | 7 | 1.4E-6 | 4.5E-5 |
| UP_SEQ_FEATURE | repeat:30 | RT | 6 | 1.6E-6 | 5.0E-5 |
| UP_SEQ_FEATURE | repeat:32 | RT | 5 | 3.6E-6 | 1.1E-4 |
| UP_SEQ_FEATURE | repeat:29 | RT | 6 | 7.3E-6 | 2.1E-4 |
| UP_SEQ_FEATURE | repeat:31 | RT | 5 | 1.2E-5 | 3.2E-4 |
| UP_SEQ_FEATURE | repeat:37 | RT | 4 | 3.1E-5 | 8.0E-4 |
| UP_SEQ_FEATURE | repeat:38 | RT | 4 | 3.1E-5 | 8.0E-4 |
| UP_SEQ_FEATURE | repeat:24 | RT | 6 | 4.4E-5 | 1.1E-3 |
| UP_SEQ_FEATURE | repeat:40 | RT | 4 | 5.3E-5 | 1.3E-3 |
| UP_SEQ_FEATURE | repeat:35 | RT | 4 | 8.4E-5 | 2.0E-3 |
| UP_SEQ_FEATURE | repeat:34 | RT | 4 | 1.2E-4 | 2.9E-3 |
| INTERPRO | SEA domain | RT | 5 | 1.7E-4 | 6.7E-3 |
| UP_SEQ_FEATURE | repeat:33 | RT | 4 | 1.8E-4 | 4.1E-3 |
| UP_SEQ_FEATURE | repeat:36; approximate | RT | 3 | 4.0E-4 | 9.1E-3 |
| UP_SEQ_FEATURE | domain:SEA | RT | 4 | 7.8E-4 | 1.7E-2 |
| UP_SEQ_FEATURE | repeat:39 | RT | 3 | 2.0E-3 | 3.9E-2 |
| UP_SEQ_FEATURE | repeat:41 | RT | 3 | 2.0E-3 | 3.9E-2 |
| **Annotation Cluster 3** | **Enrichment Score: 6.34** | | **Count** | **P_Value** | **Benjamini** |
| UP_SEQ_FEATURE | domain:TSP N-terminal | RT | 8 | 5.6E-9 | 2.7E-7 |
| SMART | TSPN | RT | 8 | 3.0E-8 | 1.0E-6 |

#### Functional Annotation Clustering

Help and Manual

**Current Gene List: HLA-C0202_enr**
**Current Background: Homo sapiens**
**1015 DAVID IDs**

Options Classification Stringency Medium

Rerun using options | Create Sublist

**72 Cluster(s)**

Download File

| Annotation Cluster 1 | Enrichment Score: ? | | Count | P_Value | Benjamini |
|---|---|---|---|---|---|
| UP_SEQ_FEATURE | transmembrane region | RT | 939 | 0.0E0 | 0.0E0 |
| GOTERM_CC_DIRECT | integral component of membrane | RT | 898 | 0.0E0 | 0.0E0 |
| UP_KEYWORDS | Transmembrane helix | RT | 954 | 0.0E0 | 0.0E0 |
| UP_KEYWORDS | Membrane | RT | 965 | 0.0E0 | 0.0E0 |
| UP_KEYWORDS | Transmembrane | RT | 954 | 0.0E0 | 0.0E0 |
| **Annotation Cluster 2** | **Enrichment Score: ?** | | **Count** | **P_Value** | **Benjamini** |
| UP_KEYWORDS | Transducer | RT | 511 | 0.0E0 | 0.0E0 |
| INTERPRO | G protein-coupled receptor, rhodopsin-like | RT | 483 | 0.0E0 | 0.0E0 |
| KEGG_PATHWAY | Olfactory transduction | RT | 358 | 0.0E0 | 0.0E0 |
| UP_KEYWORDS | Olfaction | RT | 368 | 0.0E0 | 0.0E0 |
| INTERPRO | Olfactory receptor | RT | 367 | 0.0E0 | 0.0E0 |
| UP_KEYWORDS | Receptor | RT | 529 | 0.0E0 | 0.0E0 |
| GOTERM_MF_DIRECT | olfactory receptor activity | RT | 368 | 0.0E0 | 0.0E0 |
| INTERPRO | GPCR, rhodopsin-like, 7TM | RT | 491 | 0.0E0 | 0.0E0 |
| GOTERM_BP_DIRECT | G-protein coupled receptor signaling pathway | RT | 460 | 0.0E0 | 0.0E0 |
| GOTERM_BP_DIRECT | detection of chemical stimulus involved in sensory perception of smell | RT | 367 | 0.0E0 | 0.0E0 |
| GOTERM_MF_DIRECT | G-protein coupled receptor activity | RT | 459 | 0.0E0 | 0.0E0 |
| UP_KEYWORDS | G-protein coupled receptor | RT | 510 | 0.0E0 | 0.0E0 |
| UP_KEYWORDS | Sensory transduction | RT | 395 | 0.0E0 | 0.0E0 |
| UP_SEQ_FEATURE | topological domain:Extracellular | RT | 639 | 2.6E-314 | 1.6E-311 |
| UP_SEQ_FEATURE | topological domain:Cytoplasmic | RT | 678 | 9.2E-295 | 3.8E-292 |
| UP_KEYWORDS | Cell membrane | RT | 608 | 2.4E-246 | 3.2E-244 |
| UP_SEQ_FEATURE | glycosylation site:N-linked (GlcNAc...) | RT | 627 | 2.1E-189 | 6.3E-187 |
| GOTERM_CC_DIRECT | plasma membrane | RT | 648 | 2.3E-189 | 2.6E-187 |
| UP_KEYWORDS | Glycoprotein | RT | 637 | 8.3E-181 | 7.5E-179 |
| UP_SEQ_FEATURE | disulfide bond | RT | 448 | 8.8E-125 | 2.2E-122 |
| UP_KEYWORDS | Disulfide bond | RT | 473 | 7.7E-116 | 5.2E-114 |
| **Annotation Cluster 3** | **Enrichment Score: 10.27** | | **Count** | **P_Value** | **Benjamini** |
| GOTERM_MF_DIRECT | bitter taste receptor activity | RT | 18 | 1.0E-16 | 1.6E-14 |
| INTERPRO | Mammalian taste receptor | RT | 18 | 2.7E-16 | 2.3E-14 |
| UP_KEYWORDS | Taste | RT | 18 | 5.4E-14 | 1.6E-12 |
| GOTERM_BP_DIRECT | detection of chemical stimulus involved in sensory perception of bitter taste | RT | 18 | 8.5E-12 | 2.2E-9 |
| KEGG_PATHWAY | Taste transduction | RT | 18 | 9.3E-8 | 5.0E-6 |
| GOTERM_BP_DIRECT | sensory perception of taste | RT | 10 | 1.6E-5 | 9.2E-4 |

#### Functional Annotation Clustering

Help and Manual

**Current Gene List: HLA-C0202_depl**
**Current Background: Homo sapiens**
**770 DAVID IDs**

Options Classification Stringency Medium

Rerun using options | Create Sublist

**91 Cluster(s)**

Download File

| | | | | | | Count | P_Value | Benjamini |
|---|---|---|---|---|---|---|---|---|
| Annotation Cluster 1 | | Enrichment Score: 19.72 | | | | Count | P_Value | Benjamini |
| | UP_KEYWORDS | mRNA splicing | RT | | | 54 | 1.8E-24 | 1.8E-22 |
| | UP_KEYWORDS | mRNA processing | RT | | | 58 | 2.4E-22 | 1.2E-20 |
| | GOTERM_BP_DIRECT | RNA splicing | RT | | | 38 | 8.2E-20 | 1.5E-16 |
| | GOTERM_BP_DIRECT | mRNA processing | RT | | | 34 | 3.7E-15 | 3.4E-12 |
| Annotation Cluster 2 | | Enrichment Score: 19.47 | | | | Count | P_Value | Benjamini |
| | UP_SEQ_FEATURE | repeat:7 | RT | | | 44 | 4.6E-29 | 1.0E-25 |
| | UP_SEQ_FEATURE | repeat:8 | RT | | | 42 | 3.3E-28 | 3.7E-25 |
| | UP_SEQ_FEATURE | repeat:6 | RT | | | 43 | 5.3E-26 | 3.9E-23 |
| | UP_SEQ_FEATURE | repeat:10 | RT | | | 36 | 1.4E-25 | 6.4E-23 |
| | UP_SEQ_FEATURE | repeat:3 | RT | | | 51 | 3.1E-25 | 1.2E-22 |
| | UP_SEQ_FEATURE | repeat:9 | RT | | | 37 | 3.2E-25 | 1.0E-22 |
| | UP_SEQ_FEATURE | repeat:13 | RT | | | 33 | 1.6E-24 | 4.6E-22 |
| | UP_SEQ_FEATURE | repeat:5 | RT | | | 43 | 2.7E-24 | 6.7E-22 |
| | UP_SEQ_FEATURE | repeat:1 | RT | | | 53 | 3.6E-24 | 7.9E-22 |
| | UP_SEQ_FEATURE | repeat:12 | RT | | | 33 | 3.9E-24 | 7.9E-22 |
| | UP_SEQ_FEATURE | repeat:14 | RT | | | 32 | 4.7E-24 | 8.8E-22 |
| | UP_SEQ_FEATURE | repeat:15 | RT | | | 31 | 5.2E-24 | 9.0E-22 |
| | UP_SEQ_FEATURE | repeat:2 | RT | | | 53 | 6.3E-24 | 1.0E-21 |
| | UP_SEQ_FEATURE | repeat:4 | RT | | | 46 | 7.6E-24 | 1.1E-21 |
| | UP_SEQ_FEATURE | repeat:11 | RT | | | 33 | 1.4E-23 | 1.8E-21 |
| | UP_SEQ_FEATURE | repeat:18 | RT | | | 26 | 2.6E-23 | 3.2E-21 |
| | UP_SEQ_FEATURE | repeat:17 | RT | | | 27 | 7.7E-23 | 9.0E-21 |
| | UP_SEQ_FEATURE | repeat:19 | RT | | | 25 | 3.6E-22 | 4.1E-20 |
| | UP_SEQ_FEATURE | repeat:16 | RT | | | 27 | 1.2E-21 | 1.2E-19 |
| | UP_SEQ_FEATURE | repeat:20 | RT | | | 24 | 2.7E-21 | 2.7E-19 |
| | UP_SEQ_FEATURE | repeat:21 | RT | | | 23 | 1.0E-20 | 1.0E-18 |
| | UP_SEQ_FEATURE | repeat:22 | RT | | | 22 | 7.7E-20 | 7.1E-18 |
| | UP_SEQ_FEATURE | repeat:23 | RT | | | 20 | 4.2E-18 | 3.7E-16 |
| | UP_SEQ_FEATURE | repeat:25 | RT | | | 18 | 4.4E-16 | 3.8E-14 |
| | UP_SEQ_FEATURE | repeat:28 | RT | | | 16 | 1.4E-15 | 1.2E-13 |
| | UP_SEQ_FEATURE | repeat:24 | RT | | | 17 | 1.2E-14 | 8.0E-13 |
| | UP_SEQ_FEATURE | repeat:27 | RT | | | 16 | 1.3E-14 | 8.8E-13 |
| | UP_SEQ_FEATURE | repeat:26 | RT | | | 16 | 4.7E-14 | 3.1E-12 |
| | UP_SEQ_FEATURE | repeat:29 | RT | | | 14 | 3.6E-13 | 2.3E-11 |
| | INTERPRO | High sulphur keratin-associated protein | RT | | | 16 | 1.3E-9 | 1.3E-7 |
| | UP_SEQ_FEATURE | repeat:30 | RT | | | 10 | 4.3E-9 | 2.6E-7 |
| | GOTERM_CC_DIRECT | keratin filament | RT | | | 19 | 2.9E-8 | 1.8E-6 |
| | UP_KEYWORDS | Keratin | RT | | | 19 | 1.7E-5 | 2.3E-4 |

| Annotation Cluster 2 | Enrichment Score: 19.47 | | Count | P_Value | Benjamini |
|---|---|---|---|---|---|
| UP_SEQ_FEATURE | repeat:7 | RT | 44 | 4.6E-29 | 1.0E-25 |
| UP_SEQ_FEATURE | repeat:8 | RT | 42 | 3.3E-28 | 3.7E-25 |
| UP_SEQ_FEATURE | repeat:6 | RT | 43 | 5.3E-26 | 3.9E-23 |
| UP_SEQ_FEATURE | repeat:10 | RT | 36 | 1.4E-25 | 6.4E-23 |
| UP_SEQ_FEATURE | repeat:3 | RT | 51 | 3.1E-25 | 1.2E-22 |
| UP_SEQ_FEATURE | repeat:9 | RT | 37 | 3.2E-25 | 1.0E-22 |
| UP_SEQ_FEATURE | repeat:13 | RT | 33 | 1.6E-24 | 4.6E-22 |
| UP_SEQ_FEATURE | repeat:5 | RT | 43 | 2.7E-24 | 6.7E-22 |
| UP_SEQ_FEATURE | repeat:1 | RT | 53 | 3.6E-24 | 7.9E-22 |
| UP_SEQ_FEATURE | repeat:12 | RT | 33 | 3.9E-24 | 7.9E-22 |
| UP_SEQ_FEATURE | repeat:14 | RT | 32 | 4.7E-24 | 8.8E-22 |
| UP_SEQ_FEATURE | repeat:15 | RT | 31 | 5.2E-24 | 9.0E-22 |
| UP_SEQ_FEATURE | repeat:2 | RT | 53 | 6.3E-24 | 1.0E-21 |
| UP_SEQ_FEATURE | repeat:4 | RT | 46 | 7.6E-24 | 1.1E-21 |
| UP_SEQ_FEATURE | repeat:11 | RT | 33 | 1.4E-23 | 1.8E-21 |
| UP_SEQ_FEATURE | repeat:18 | RT | 26 | 2.6E-23 | 3.2E-21 |
| UP_SEQ_FEATURE | repeat:17 | RT | 27 | 7.7E-23 | 9.0E-21 |
| UP_SEQ_FEATURE | repeat:19 | RT | 25 | 3.6E-22 | 4.1E-20 |
| UP_SEQ_FEATURE | repeat:16 | RT | 27 | 1.2E-21 | 1.2E-19 |
| UP_SEQ_FEATURE | repeat:20 | RT | 24 | 2.7E-21 | 2.7E-19 |
| UP_SEQ_FEATURE | repeat:21 | RT | 23 | 1.0E-20 | 1.0E-18 |
| UP_SEQ_FEATURE | repeat:22 | RT | 22 | 7.7E-20 | 7.1E-18 |
| UP_SEQ_FEATURE | repeat:23 | RT | 20 | 4.2E-18 | 3.7E-16 |
| UP_SEQ_FEATURE | repeat:25 | RT | 18 | 4.4E-16 | 3.8E-14 |
| UP_SEQ_FEATURE | repeat:28 | RT | 16 | 1.4E-15 | 1.2E-13 |
| UP_SEQ_FEATURE | repeat:24 | RT | 17 | 1.2E-14 | 8.0E-13 |
| UP_SEQ_FEATURE | repeat:27 | RT | 16 | 1.3E-14 | 8.8E-13 |
| UP_SEQ_FEATURE | repeat:26 | RT | 16 | 4.7E-14 | 3.1E-12 |
| UP_SEQ_FEATURE | repeat:29 | RT | 14 | 3.6E-13 | 2.3E-11 |
| INTERPRO | High sulphur keratin-associated protein | RT | 16 | 1.3E-9 | 1.3E-7 |
| UP_SEQ_FEATURE | repeat:30 | RT | 10 | 4.3E-9 | 2.6E-7 |
| GOTERM_CC_DIRECT | keratin filament | RT | 19 | 2.9E-8 | 1.8E-6 |
| UP_KEYWORDS | Keratin | RT | 19 | 1.7E-5 | 2.3E-4 |
| **Annotation Cluster 3** | **Enrichment Score: 12.62** | | **Count** | **P_Value** | **Benjamini** |
| INTERPRO | RNA recognition motif domain | RT | 40 | 5.5E-16 | 1.3E-13 |
| INTERPRO | Nucleotide-binding, alpha-beta plait | RT | 43 | 7.5E-16 | 1.3E-13 |
| UP_SEQ_FEATURE | domain:RRM | RT | 30 | 1.0E-14 | 7.3E-13 |
| SMART | RRM | RT | 38 | 3.5E-14 | 6.0E-12 |
| GOTERM_MF_DIRECT | nucleotide binding | RT | 45 | 1.1E-12 | 1.5E-10 |
| UP_KEYWORDS | RNA-binding | RT | 66 | 1.4E-12 | 3.8E-11 |
| GOTERM_MF_DIRECT | RNA binding | RT | 47 | 2.1E-7 | 1.4E-5 |
| **Annotation Cluster 4** | **Enrichment Score: 11.62** | | **Count** | **P_Value** | **Benjamini** |
| UP_KEYWORDS | mRNA splicing | RT | 54 | 1.8E-24 | 1.8E-22 |
| GOTERM_BP_DIRECT | mRNA splicing, via spliceosome | RT | 33 | 1.0E-11 | 3.8E-9 |
| UP_KEYWORDS | Spliceosome | RT | 19 | 1.2E-6 | 2.2E-5 |
| GOTERM_CC_DIRECT | catalytic step 2 spliceosome | RT | 16 | 1.6E-6 | 6.8E-5 |

#### Functional Annotation Clustering

Help and Manual

**Current Gene List: HLA-C1507_enr**
**Current Background: Homo sapiens**
**1170 DAVID IDs**

Options   Classification Stringency Medium

Rerun using options   Create Sublist

**86 Cluster(s)**

Download File

| Annotation Cluster 1 | Enrichment Score: ? | | | Count | P_Value | Benjamini |
|---|---|---|---|---|---|---|
| UP_SEQ_FEATURE | transmembrane region | RT | | 1072 | 0.0E0 | 0.0E0 |
| UP_KEYWORDS | Membrane | RT | | 1105 | 0.0E0 | 0.0E0 |
| GOTERM_CC_DIRECT | integral component of membrane | RT | | 1011 | 0.0E0 | 0.0E0 |
| UP_KEYWORDS | Transmembrane | RT | | 1098 | 0.0E0 | 0.0E0 |
| UP_KEYWORDS | Transmembrane helix | RT | | 1098 | 0.0E0 | 0.0E0 |
| **Annotation Cluster 2** | **Enrichment Score: ?** | | | **Count** | **P_Value** | **Benjamini** |
| INTERPRO | Olfactory receptor | RT | | 372 | 0.0E0 | 0.0E0 |
| GOTERM_BP_DIRECT | G-protein coupled receptor signaling pathway | RT | | 494 | 0.0E0 | 0.0E0 |
| GOTERM_MF_DIRECT | olfactory receptor activity | RT | | 373 | 0.0E0 | 0.0E0 |
| UP_KEYWORDS | Olfaction | RT | | 373 | 0.0E0 | 0.0E0 |
| UP_SEQ_FEATURE | topological domain:Cytoplasmic | RT | | 767 | 0.0E0 | 0.0E0 |
| KEGG_PATHWAY | Olfactory transduction | RT | | 362 | 0.0E0 | 0.0E0 |
| UP_KEYWORDS | Sensory transduction | RT | | 408 | 0.0E0 | 0.0E0 |
| UP_SEQ_FEATURE | topological domain:Extracellular | RT | | 720 | 0.0E0 | 0.0E0 |
| UP_KEYWORDS | Transducer | RT | | 557 | 0.0E0 | 0.0E0 |
| INTERPRO | G protein-coupled receptor, rhodopsin-like | RT | | 523 | 0.0E0 | 0.0E0 |
| UP_KEYWORDS | Receptor | RT | | 585 | 0.0E0 | 0.0E0 |
| INTERPRO | GPCR, rhodopsin-like, 7TM | RT | | 533 | 0.0E0 | 0.0E0 |
| UP_KEYWORDS | G-protein coupled receptor | RT | | 557 | 0.0E0 | 0.0E0 |
| GOTERM_BP_DIRECT | detection of chemical stimulus involved in sensory perception of smell | RT | | 372 | 0.0E0 | 0.0E0 |
| GOTERM_MF_DIRECT | G-protein coupled receptor activity | RT | | 492 | 0.0E0 | 0.0E0 |
| UP_KEYWORDS | Cell membrane | RT | | 674 | 9.0E-261 | 1.4E-258 |
| UP_SEQ_FEATURE | glycosylation site:N-linked (GlcNAc...) | RT | | 702 | 3.2E-205 | 2.3E-202 |
| GOTERM_CC_DIRECT | plasma membrane | RT | | 725 | 4.2E-200 | 6.1E-198 |
| UP_KEYWORDS | Glycoprotein | RT | | 717 | 4.7E-196 | 4.7E-194 |
| UP_SEQ_FEATURE | disulfide bond | RT | | 481 | 3.0E-121 | 1.5E-118 |
| UP_KEYWORDS | Disulfide bond | RT | | 509 | 1.6E-110 | 1.2E-108 |
| **Annotation Cluster 3** | **Enrichment Score: 15.64** | | | **Count** | **P_Value** | **Benjamini** |
| UP_KEYWORDS | Symport | RT | | 47 | 5.3E-28 | 2.7E-26 |
| UP_KEYWORDS | Ion transport | RT | | 87 | 8.2E-14 | 2.5E-12 |
| GOTERM_BP_DIRECT | sodium ion transport | RT | | 28 | 2.1E-13 | 4.5E-11 |
| UP_KEYWORDS | Sodium | RT | | 33 | 2.6E-13 | 7.1E-12 |
| UP_KEYWORDS | Sodium transport | RT | | 32 | 2.8E-13 | 7.1E-12 |

#### Functional Annotation Clustering

[Help and Manual](#)

**Current Gene List: HLA-C1507_depl**
**Current Background: Homo sapiens**
**674 DAVID IDs**

**Options** **Classification Stringency** Medium

Rerun using options | Create Sublist

**85 Cluster(s)**

**Download File**

| Annotation Cluster 1 | Enrichment Score: 17.66 | | Count | P_Value | Benjamini |
|---|---|---|---|---|---|
| INTERPRO | Nucleotide-binding, alpha-beta plait | RT | 50 | 7.4E-24 | 2.4E-21 |
| INTERPRO | RNA recognition motif domain | RT | 46 | 2.7E-23 | 5.7E-21 |
| SMART | RRM | RT | 45 | 2.4E-21 | 4.1E-19 |
| UP_KEYWORDS | RNA-binding | RT | 75 | 1.4E-20 | 6.6E-19 |
| UP_SEQ_FEATURE | domain:RRM | RT | 33 | 4.2E-19 | 4.1E-16 |
| GOTERM_MF_DIRECT | nucleotide binding | RT | 50 | 1.8E-18 | 2.5E-16 |
| GOTERM_MF_DIRECT | RNA binding | RT | 57 | 1.6E-14 | 1.3E-12 |
| GOTERM_MF_DIRECT | nucleic acid binding | RT | 59 | 6.3E-6 | 2.9E-4 |
| **Annotation Cluster 2** | **Enrichment Score: 17.34** | | **Count** | **P_Value** | **Benjamini** |
| UP_KEYWORDS | mRNA splicing | RT | 56 | 4.5E-29 | 6.5E-27 |
| UP_KEYWORDS | mRNA processing | RT | 62 | 1.4E-28 | 1.4E-26 |
| KEGG_PATHWAY | Spliceosome | RT | 29 | 3.7E-20 | 5.3E-18 |
| GOTERM_BP_DIRECT | RNA splicing | RT | 36 | 6.8E-20 | 1.3E-16 |
| GOTERM_BP_DIRECT | mRNA splicing, via spliceosome | RT | 38 | 2.2E-17 | 2.1E-14 |
| GOTERM_BP_DIRECT | mRNA processing | RT | 33 | 5.4E-16 | 3.4E-13 |
| GOTERM_CC_DIRECT | catalytic step 2 spliceosome | RT | 16 | 2.6E-7 | 1.1E-5 |
| UP_KEYWORDS | Spliceosome | RT | 17 | 4.0E-6 | 6.8E-5 |
| **Annotation Cluster 3** | **Enrichment Score: 12.3** | | **Count** | **P_Value** | **Benjamini** |
| INTERPRO | Collagen triple helix repeat | RT | 31 | 3.0E-24 | 1.9E-21 |
| UP_KEYWORDS | Collagen | RT | 32 | 1.7E-22 | 9.8E-21 |
| GOTERM_MF_DIRECT | extracellular matrix structural constituent | RT | 26 | 1.3E-20 | 2.7E-18 |
| UP_KEYWORDS | Hydroxylation | RT | 27 | 2.4E-17 | 8.8E-16 |
| KEGG_PATHWAY | Protein digestion and absorption | RT | 22 | 1.7E-16 | 1.6E-14 |
| UP_SEQ_FEATURE | region of interest:Triple-helical region | RT | 15 | 7.5E-16 | 3.9E-13 |
| GOTERM_CC_DIRECT | collagen trimer | RT | 25 | 1.3E-15 | 2.9E-13 |
| UP_KEYWORDS | Extracellular matrix | RT | 40 | 1.3E-15 | 3.9E-14 |
| GOTERM_BP_DIRECT | collagen catabolic process | RT | 21 | 1.9E-15 | 9.3E-13 |
| GOTERM_CC_DIRECT | endoplasmic reticulum lumen | RT | 33 | 2.4E-14 | 3.5E-12 |
| INTERPRO | Fibrillar collagen, C-terminal | RT | 10 | 1.7E-12 | 1.8E-10 |
| GOTERM_BP_DIRECT | extracellular matrix organization | RT | 30 | 1.8E-12 | 6.6E-10 |
| UP_SEQ_FEATURE | domain:Fibrillar collagen NC1 | RT | 10 | 1.9E-12 | 3.0E-10 |
| KEGG_PATHWAY | ECM-receptor interaction | RT | 18 | 5.3E-12 | 2.5E-10 |
| SMART | COLFI | RT | 10 | 6.4E-12 | 5.5E-10 |
| KEGG_PATHWAY | Amoebiasis | RT | 18 | 1.4E-10 | 5.1E-9 |
| GOTERM_CC_DIRECT | proteinaceous extracellular matrix | RT | 33 | 2.3E-10 | 1.4E-8 |
| KEGG_PATHWAY | Focal adhesion | RT | 21 | 2.8E-8 | 8.0E-7 |
| UP_SEQ_FEATURE | propeptide:C-terminal propeptide | RT | 7 | 3.0E-8 | 1.4E-6 |
| GOTERM_BP_DIRECT | collagen fibril organization | RT | 10 | 2.0E-6 | 3.7E-4 |
